# The proteome is a terminal electron acceptor

**DOI:** 10.1101/2024.01.31.578293

**Authors:** Avi I. Flamholz, Akshit Goyal, Woodward W. Fischer, Dianne K. Newman, Rob Phillips

## Abstract

Microbial metabolism is impressively flexible, enabling growth even when available nutrients differ greatly from biomass in redox state. *E. coli*, for example, rearranges its physiology to grow on reduced and oxidized carbon sources through several forms of fermentation and respiration. To understand the limits on and evolutionary consequences of metabolic flexibility, we developed a mathematical model coupling redox chemistry with principles of cellular resource allocation. Our integrated model clarifies key phenomena, including demonstrating that autotrophs grow slower than heterotrophs because of constraints imposed by intracellular production of reduced carbon. Our model further indicates that growth is improved by adapting the redox state of biomass to nutrients, revealing an unexpected mode of evolution where proteins accumulate mutations benefiting organismal redox balance.

**One sentence summary:** Microbial proteins adapt their composition on physiological and evolutionary timescales to ensure organismal redox balance.

## Main Text

Many microbes have a remarkable capacity for metabolic chemistry, growing on a diversity of environmentally-supplied nutrients (e.g. carbon, nitrogen and sulfur sources) in both oxic and anoxic settings. *E. coli* strains, for example, can grow on relatively reduced (e.g. fatty acids) and oxidized carbon (C) sources (e.g. organic acids) through several forms of fermentation and respiration (Fig. 1A). Some microbes can even switch their fundamental metabolic mode (*1*, *^2^*), eating organic molecules in some conditions (i.e. performing heterotrophy) while fixing CO_2_ into organics to build biomass in others (i.e. performing autotrophy). Not all microbes are so flexible, yet this widespread metabolic flexibility is noteworthy because it is essentially absent from animals.

**Figure 1:**
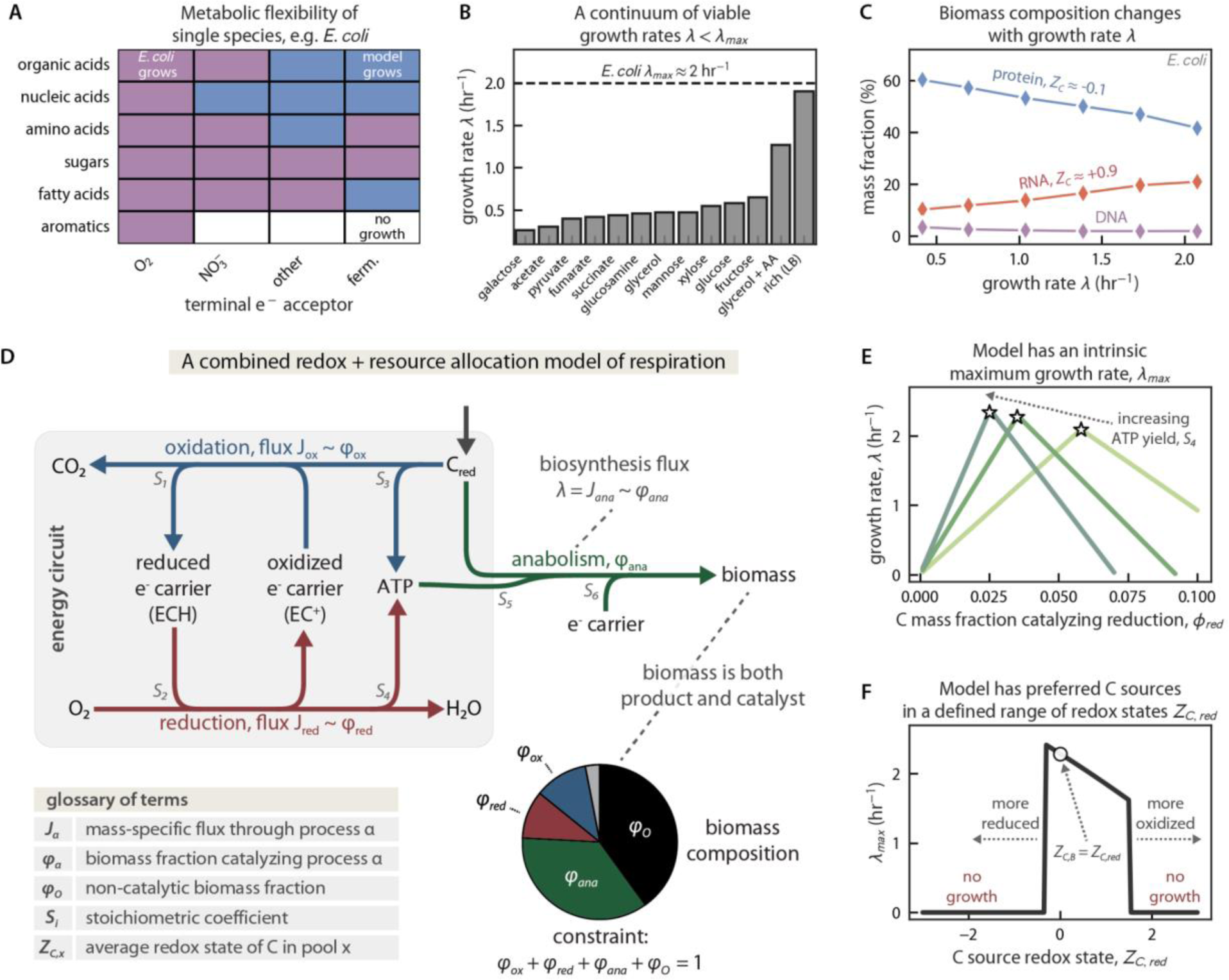
Integrating redox chemistry with principles of cellular resource allocation captures key features of microbial physiology. (A) Many microbes can grow on a diversity of C sources (Y axis) and e^-^ acceptors (X axis), with “other” denoting trimethylamine N-oxide or dimethyl sulfoxide and “ferm.” fermentation. Purple squares indicate observed growth of *E. coli* strains, blue predictions from a metabolic model, and white no growth in the lab or in silico (see *Methods*). (B) Microbes have an intrinsic maximum growth rate, *λ*_max_, that arises in nutrient surplus, but can rearrange their internal physiology to grow at nearly any growth rate *λ* < *λ*_max_. Illustrated here with *E. coli* grown on various C sources (*51*), but also observed with variable nutrient concentrations. (C) As ribosomes are ≈⅔ RNA, faster-growing cells typically have more RNA (C redox state *Z_C_* ≈ + 0.9) and less protein (*Z_C_* ≈ -0.1). *E. coli* data from (*12*). (D) An integrated model of heterotrophic respiration comprises 3 redox reactions. Oxidation (blue) withdraws e^-^ from the reduced C source (*C_red_*), generating reduced e^-^ carrier (ECH). Reduction (red) oxidizes ECH to EC^+^ while reducing a favorable e^-^ acceptor (here O_2_), coupling this favorable e^-^ transfer to ATP synthesis. Anabolism (green) uses ATP and, potentially, ECH to convert *C_red_* into biomass. The mass-specific flux *J*_⍺_ through process ⍺ is proportional to the C mass fraction of catalyst, *φ*_⍺_, and *λ* is determined by the anabolic flux *J_ana_*. Reactions are written per C atom with stoichiometries (*S_i_*) as labeled. To sustain growth, cells must allocate resources to synthesis of oxidative (*φ*_ox_), reductive (*φ*_red_), and anabolic (*φ*_ana_) enzymes so that flux (*J_ox_*, *J_red_*, and *J_ana_*) balance to conserve C atoms and e^-^. The remaining non-catalytic mass, encompassing structural components like lipids and cell walls, is termed *φ*_o_. (E) This model has an intrinsic maximum growth rate, *λ*_max_, arising because *φ*_⍺_trade-off with each other (pie chart in panel D). When *φ*_red_ is small, for example, increasing its value increases *λ* by providing ATP used in anabolism. When *φ*_red_ is large, increasing its value decreases *λ* by displacing *φ*_ana_. Stars mark *λ*_max_ on each curve. (F) Our model includes the C redox states of *C_red_* (*Z_C,red_*) and biomass (*Z_C,B_*), with negative values connoting reduced molecules (more e^-^/C) and positive values oxidized (fewer e^-^/C). When *Z_C,red_* and *Z_C,B_* were very different, growth was infeasible (“no growth” where **λ*_max_* = 0). The largest **λ** values arise when *Z_C,red_* is somewhat more reduced than *Z_C,B_*, consistent with the dual roles of carbon as a source of e^-^ and C atoms.

The language of redox chemistry — intermolecular electron (e^-^) transfers — is often used to describe the consequences of metabolic flexibility. Redox logic is useful because all organisms extract energy from their environments by coupling thermodynamically-favorable e^-^ transfers to the synthesis of energy carriers like adenosine triphosphate (ATP). Using redox chemistry to “coarse-grain” metabolism is advantageous because it can describe any metabolism, enforces conservation of atoms and e^-^, and provides an opportunity to incorporate thermodynamic limits via redox potentials (*3*, *4*). Yet redox is not a sufficient description of microbial physiology. For example, microbes have intrinsic maximum growth rates that arise in nutrient surplus (*λ*_max_, Fig. 1B), but conservation of atoms and e^-^ only requires that fluxes of production and consumption balance — the fluxes, and, therefore, the growth rate, can be arbitrarily large.

In recent years, several groups have developed a theory of resource allocation to explain how microbes rearrange their internal machinery to achieve different growth rates (*5–8*). These models treat microbes as self-replicating catalysts whose growth rate, *λ*, is determined by the manner in which they apportion nutrients (e.g. carbon) and catalytic activity (e.g. protein synthesis by ribosomes) to produce the enzymes that perform core cellular tasks like catabolism and biosynthesis.

Considering cells as resource-allocating self-replicators can explain the existence of an intrinsic maximum growth rate, *λ*_max_, which arises when resources are optimally divided between catabolism and biosynthesis so that influxes of precursors are matched by the anabolic processes consuming them (*5–8*). Furthermore, because biological macromolecules play different cellular roles — metabolic reactions are catalyzed by proteins, but ribosomes are 60-70% RNA by mass (*9*) — resource allocation can also explain observed changes in biomass composition that accompany increases in *λ* (*5–8*), including increases in the RNA:protein ratio (*10–12*).

In addition to its central role in ribosomes, RNA is also fairly oxidized (e^-^ poor), with a formal carbon redox state of *Z_C_* ≈ +0.9 charge units per carbon (Fig. 1C). In contrast, values for proteins are about 1 e^-^/C more reduced (e^-^ rich), ranging from -0.1 to -0.3 (*4*, *13*). As such, increasing RNA:protein ratios entail the oxidation of biomass during faster growth (Fig. 1C). To understand how concerted changes in biomass redox should affect growth, we developed an integrated redox chemical model of microbial resource allocation to study intrinsic limits on growth and metabolic rates (Fig. 1D). As microbial communities are pivotal contributors to global recycling of carbon, nitrogen, phosphorus and sulfur (*14*), a theory that describes the intrinsic limits of microbial activities would offer predictive value (*15*). Here we describe such a theory, draw lessons and predictions from it, and offer empirical support for those predictions from the genomes and proteomes of diverse microbes.

## Results

### A redox chemical model of microbial self-replication

To merge redox and resource-economic descriptions, we track the redox state of carbon (*Z_C_*) in growth substrates (e.g., an organic C source like glucose) and products (e.g., CO_2_, biomass). *Z_C_* counts the average number of valence e^-^ associated with C atoms in a molecule (*4*). Electrons are counted by their charge, with negative *Z_C_* denoting excess e^-^ compared to neutrally-charged C atoms (4 valence e^-^) and positive *Z_C_* denoting paucity.

Consider heterotrophic respiration — the net transfer of e^-^ from organic molecules to a terminal acceptor like O_2_ as depicted in Fig. 1D. This metabolism is described by three coupled redox reactions: (i) oxidation, (ii) reduction, and (iii) anabolism. Oxidation extracts e^-^ from the reduced C source (*C_red_*) while reduction drives respiratory ATP production by harnessing the flow of e^-^ to O_2_ to generate a proton-motive force driving an ATP synthase (*16*). Not all C atoms can be oxidized to make ATP, however, or there will be no growth or maintenance of biomass. Rather, some organic C is used in anabolism to make new biomass, with a mass-specific flux of

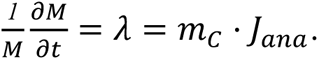

Here *M* is the carbon mass of cells, **λ** is the growth rate (C mass doublings per hour), *J_ana_* the anabolic flux (mol C/g C/hr), and *m_C_* is the molar mass of C. In cells, e^-^ are carried between these processes by soluble molecules like nicotinamides (e.g. NADH), flavins, and quinones which we represent as a generic redox couple ECH/EC^+^ for “electron carrier,” ECH being the reduced (e^-^ carrying) form. Reduction of the terminal e^-^ acceptor therefore exchanges *S_2_* ECH for *S_4_* ATP where *S_i_* are stoichiometric coefficients labeled in Fig. 1D. Because C sources typically differ from biomass in *Z_C_*, anabolism consumes *S_6_* ≠ 0 ECH per C atom.

Conservation of mass requires conservation of electrons, which occurs when

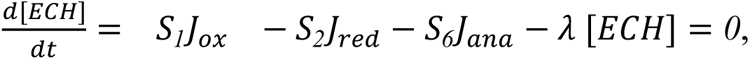

where **λ**[ECH] accounts for dilution of the e^-^ carrier due to growth. *S_1_* = ½ (*Z_C,ox_* - *Z_C,red_*) and *S_6_* = ½ (*Z_C,B_* - *Z_C,red_*) are calculated from *Z_C_* values for biomass (*Z_C,B_*), the C source (*C_red_* with *Z_C,red_*) and the oxidized product (e.g. *Z_C,ox_* = +4 for CO_2_). So a model tracking *Z_C_* conserves e^-^ in addition to C atoms and biosynthetic activity.

In addition to balancing redox reactions, cells must also extract sufficient ATP for maintenance (*b*) and biosynthesis (*S_5_ J_ana_*):

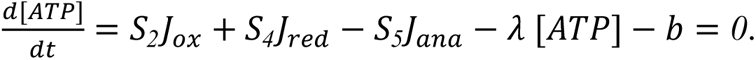

One fundamental aspect of growth missing from this representation is autocatalysis: biomass is both the product and the catalyst of growth (*5–8*). As such, each of the mass-specific fluxes *J_ox_*, *J_red_*, and *J_ana,_* (generically *J_α_*) depend on the amount of cellular catalyst (C mass *Φ_α_M*) and its mass-specific activity (*γ_α_*, mol/g C/hr)

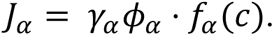

Cells manipulate *Φ_α_* by regulating the synthesis of enzymes and macromolecules. Here *f_α_*(**c**) is relative catalytic activity as a function of reactant concentrations (via the vector **c**), accounting for substrate saturation, product inhibition, and concentration-dependent regulation, as described in the supplement.

Recognizing the autocatalytic constraint on growth, we subdivide biomass into its catalytic (*Φ_ox_, Φ_red_, and Φ_ana_*) and non-catalytic (*Φ_O_*) components so that a proportionality between, for example, *J_red_* and *Φ_red_* can be enforced. An allocation constraint then requires that all of biomass is accounted for:

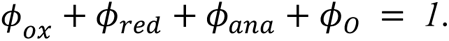

As a result of this constraint, and in contrast with purely redox descriptions of metabolism, our unified model displays a maximum growth rate **λ**_max_ because allocation of more biomass to one process (e.g. increasing *Φ_red_*) displaces another (e.g. *Φ_ana_*, Fig. 1E).

### Chemical limits on growth and metabolic rates

Since cells must apportion finite catalytic capacity (i.e. enzyme mass) between C source oxidation, acceptor reduction, and biosynthesis, a maximum growth rate *λ*_max_ arises when increasing the anabolic allocation *Φ_ana_* no longer increases the biosynthetic flux *m_C_*·*J_ana_* = *λ*. This occurs because increasing *Φ_ana_* further displaces oxidation or reduction, lowering production of ATP or ECH beneath the level needed to support *J_ana_* and, therefore, *λ*.

Given a set of model parameters and limits on *Φ_O_*, one can calculate *λ*_max_ and *J_red_*\*, a flux beyond which additional respiratory CO_2_ production no longer increases *λ* (Fig. 2A and Fig. S6). These values depend on biochemical kinetics (*γ_α_*), stoichiometries (e.g., *S_4_* ATP produced per O_2_ reduced) and the redox states of nutrients and biomass (*Z_C,red_* and *Z_C,B_*, Figs. 2B-D). As such, parameterization of our model permits principled calculation of microbial carbon use efficiencies — values that represent the fraction of carbon taken up from the environment that is incorporated into biomass, which are vital for understanding and predicting microbial carbon cycling in soils and sediments (*17–20*).

**Figure 2.**
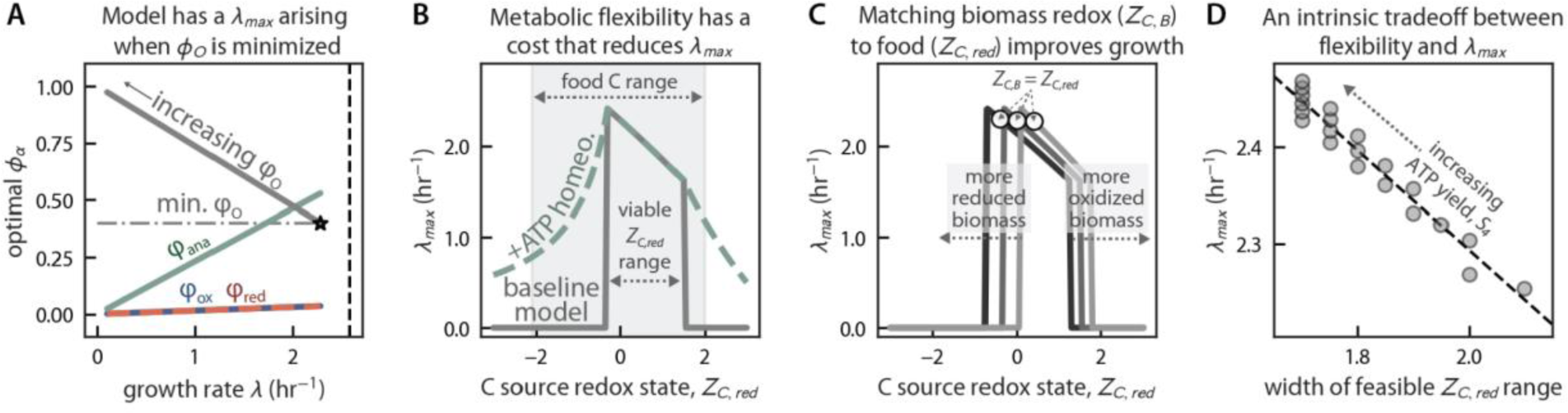
Predictions about heterotrophic physiology from an integrated redox and resource allocation model. (A) Like microbial populations, our integrated model has an intrinsic maximum growth rate, *λ*_max_ (star), arising when *φ*_O_ achieves its minimum allowed value (horizontal gray line). As such, *φ*_O_ increases as *λ* decreases. The vertical dashed line gives a simple estimate of *λ*_max_ ≈ *γ*_ana_(1-*φ*_o,min_) made by assuming all catalytic mass is anabolic. (B) Varying *Z_C,red_*, the redox state of the C source, affects both oxidative yields of reduced e^-^ carriers (*S_1_*) and anabolic consumption of reduced carriers (*S_6_*, see Fig. 1D). When the C source is too reduced or oxidized, growth becomes infeasible (“baseline model” in gray). This failure is due to an inability to simultaneously balance flows of e^-^, ATP, and C. Enabling balancing of ATP and e^-^ carriers by hydrolysis of excess ATP permitted growth at any *Z_C,red_* (dashed green). The gray range gives the approximate span of *Z_C,red_* for organic C sources from (*4*). (C) Each gray curve represents a model with a distinct biomass redox state, *Z_C,B_*. Altering *Z_C,B_* shifts the range of C sources that can be consumed, but does not alter the width of the viable *Z_C,red_* range marked on panel B. As in Fig. 1F, consuming C sources that are slightly more reduced than biomass maximizes growth. (D) Varying the reductive ATP yield, *S_4_*, reveals a tradeoff between metabolic flexibility and maximum growth rate. High ATP yields (top left) produced the fastest maximum growth rates, but over the narrowest range of *Z_C,red_* values. Lower *S_4_* values (bottom right) reduced *λ*_max_ but permitted growth on a wider range of C sources (see Fig. S11).

### Understanding the role of non-catalytic compounds in heterotrophic growth

While fluxes *J_α_* are potentially nonlinear functions of substrate and product concentrations, the above model can be simplified by assuming reactions are irreversible and substrate-saturated, i.e. *f_α_*(**c**) = 1 (*21*, *22*). We study this simplified model to gain intuition about the chemical requirements of growth.

Presuming ATP and e^-^ carrier (ECH/EC^+^) concentrations are constrained to typical physiological values (≈1-10 mM), our model involves three equations and four free variables *Φ_ox_, Φ_red_, Φ_ana_,* and *Φ_O_* (Fig. 1D). When the growth rate is maximized, i.e. when *λ* = *λ*_max_, the non-catalytic biomass fraction, *Φ_O_*, must reach the minimum allowed value *Φ_O,min_* (Fig. 2A). Otherwise cells could grow faster by reducing *Φ_O_* and redistributing that mass to catalytic tasks. It follows that *Φ_O_* > *Φ_O,min_* when *λ* < *λ*_max_, i.e. that cells make more non-catalytic biomass when forced to grow at submaximal rates due to nutrient limitation, for example. In the linearized model, therefore, accepting variable *Φ_O_* ≥ *Φ_O,min_* permits a continuum of sub-maximal growth rates *λ* < *λ*_max_, with *Φ_O_* increasing as *λ* decreases (*6*). This observation reveals how non-catalytic C storage molecules like glycogen (*23*) can serve as a means of balancing fluxes of carbon, energy, and reductant to enable growth at a rate set by exogenous limitations. Consistent with this view, microbes typically accumulate C storage molecules in nutrient limited conditions, with the amount of glycogen, for example, inversely related to *λ* (*23–28*).

### A need for regulatory mechanisms balancing flows of energy, reductant and carbon

Because microbes can typically grow on a variety of carbon sources, we considered heterotrophic growth as a function of C source redox, *Z_C,red_*, which affects oxidative e^-^ yield (stoichiometric coefficient *S_1_*) and anabolic demand for e^-^ (*S_6_*). We found that growth was only feasible when *Z_C,red_* fell in a defined range; outside this range *λ_max_* = 0 (Fig. 1F). These limits on growth result from the assumption that anabolism consumes ATP and ECH in a defined ratio (*S_5_*/*S_6_* per C, Fig. 1D). If the C source is too reduced (e.g. *Z_C,red_* ≈ -2, as for fatty acids (*4*)) its oxidation yields more ECH (*S_1_* increases) but anabolism requires less (-*S_6_* decreases). These conflicting trends make it impossible for the modeled cell to grow on very reduced substrates as there is no sink for excess reductant other than producing unneeded ATP. Likewise, problems arise if the C source is too oxidized, and these challenges persist in a non-linear model where fluxes *J__α__* are saturating functions of concentrations (Figs. S7 and S8).

In photosynthetic organisms, imbalances between the supply of and demand for ATP and ECH are addressed by various kinds of light-powered cyclic e^-^ flow (*29*, *30*). In contrast to linear e^-^ flow, which produces reductant and ATP in a characteristic ratio, cyclic flow produces only ATP and can therefore be used to adjust the ATP:ECH flux ratio to match anabolic needs (e.g. the Calvin-Benson cycle consumes 3 ATP and 4 e^-^ per CO_2_ fixed). Based on the *Z_C,red_* limits of our simplified model, it is apparent that heterotrophy also requires some mechanism of ATP:ECH flux ratio balancing. Such a mechanism could include regulation of ATP production stoichiometry (i.e. altering *S_3_* or *S_4_*), a shift in biomass composition affecting ATP or ECH demand (altering *S_5_* or *S_6_*), regulated ATP hydrolysis (breakdown to ADP), or some additional sink for ECH (e.g. CO_2_ fixation as in (*31*)). Regardless of the underlying mechanisms, regulated ATP homeostasis appears to be a substantial cost for bacteria (*32*).

After adding a regulated ATP hydrolysis reaction (flux *J_H_*), growth became feasible across the full range of *Z_C,red_* values (green dashed lines in Fig. 2B), and such homeostasis was not required for growth within the above limits. Yet, outside those limits, ATP homeostasis imposes a substantial cost: *λ*_max_ decreases roughly quadratically as *Z_C,red_* diverges from the preferred range (Figs. 2B and S9). This result highlights that metabolic flexibility has a cost: heterotrophs can arrange their metabolic stoichiometry to optimize growth on a preferred range of C sources and e^-^ acceptors, but switching to a substantially different metabolism (e.g. with a very different *Z_C,red_*) requires a regulatory mechanism that ensures balanced flows of ATP, e^-^, and carbon. Moreover, if the goal is to maintain relatively fast growth rates without regulatory overhead, it is preferable to make biomass that is similar to nutrients in redox state (*Z_C,B_* ≈ *Z_C,red_*, Fig. 2C).

Thus far, our model results revealed two valuable lessons regarding physiology: (i) carbon storage molecules enable a continuum of sub-maximal growth rates (Fig. 2A), and (ii) flux-balancing regulatory mechanisms enable heterotrophic growth on carbon sources spanning a wide range of redox states (Fig. 2B). This latter point might provide context for observations that light-harvesting proteins called rhodopsins are abundantly expressed by heterotrophic microbes across Earth’s oceans (*33*), as these proteins provide a light-dependent source of ATP synthesis — an analog of cyclic e^-^ flow for heterotrophy.

### Metabolic flexibility and fast growth are conflicting goals

We saw that regulatory mechanisms are needed to permit heterotrophic growth when there is a large difference between the redox state of biomass carbon (*Z_C,B_*) and that of environmentally-supplied reduced carbon sources (*Z_C,red_*). These mechanisms can take various forms, but inevitably take up “space” in biomass and consume resources (e.g. ATP) to achieve flux balance. Such homeostasis is required only when *Z_C,red_* falls outside a preferred range. As shown in Fig. 2B-D, this range depends on the stoichiometric coefficients in the model, e.g. the ATP yield of reduction (*S_4_*) and the reductant (ECH) requirements of anabolism (*S_6_*).

Altering any of these stoichiometries – e.g. by switching oxidative (*34*) or reductive pathways (*35*, *36*) – shifts the range of carbon sources for which respiratory growth is feasible without expending biomass or energy on flux-balancing homeostatic mechanisms (marked on Fig. 2B). This range can be widened by, for example, decreasing the reductive ATP yield (*S_4_*), but this comes at the expense of decreasing *λ*_max_ (Fig 2D). As above, addition of a regulated ATP homeostasis reaction permitted growth across the entire range (Figs. 2B-D and S10). This emphasizes an equivalence between regulated pathway switching, which can alter stoichiometries (*7*, *34–36*), and other forms of flux-balancing, e.g. regulated ATP hydrolysis or cyclic e^-^ flow.

### Autotrophy is more constrained than heterotrophy

Our redox-based model naturally captures the full range of microbial metabolism, including heterotrophic respiration (Fig. 1D), photosynthesis (Fig. 3A), chemolithoautotrophy and fermentation (Fig. S2). Indeed, equations for photosynthesis (or chemoautotrophy) are nearly identical to those for heterotrophy, containing one additional mass balance constraint because autotrophs produce reduced carbon (*C_red_*) intracellularly and use it to make biomass.

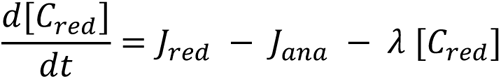

**Figure 3:**
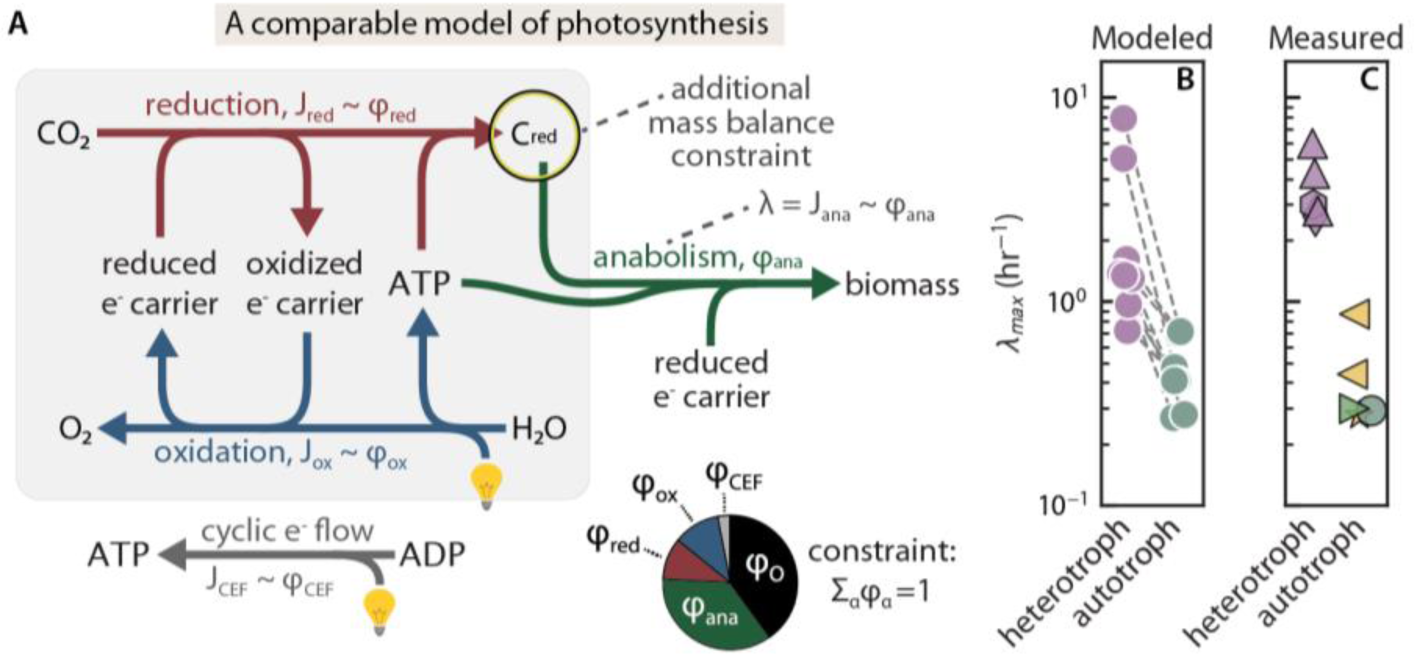
Constraints imposed only on autotrophs reduce their maximum growth rates. (A) A model of photosynthesis comprises nearly identical equations to respiration, but with two key differences. Due to the use of light energy, phototrophs can balance ATP and redox carriers with new ATP synthesis, i.e. via cyclic e^-^ flow (*J_CEF_* ∼ *φ*_CEF_). Second, because autotrophs produce and consume organic C intracellularly, fluxes of *C_red_* production, consumption and dilution must be balanced. Heterotrophs, in contrast, acquire *C_red_* from outside the cell and can deplete it. (B) Comparing models of photosynthesis and respiration with identical parameters, we found that respiratory *λ*_max_ values exceeded photosynthetic ones by approximately threefold (compare green and purple dots). (C) This result is qualitatively consistent with measurements: the fastest-growing photosynthetic microbes known have maximum growth rates that are 5-10 times slower than heterotrophic counterparts. Purple markers denote heterotrophs, green photoautotrophs, and yellow chemoautotrophs. Heterotrophic organisms: ▴ *V. natrigens* (*γ*-proteobacteria) ⬢ *E. coli* (*γ*-proteobacteria), ♦ *C. perfingens* (clostridia). Autotrophs: ⬤ *C. ohadii* (green algae) ▶ *S. elongatus* PCC 11801 (cyanobacterium), ◀ *T. crunogena* (*γ*-proteobacterial chemoautotroph).

Due to this additional constraint, models of autotrophy are defined by at least four equations. Four variables (*Φ_ox_, Φ_red_, Φ_ana_,* and *Φ_O_*) do not permit a continuum of growth rates **λ** ≤ **λ**_max_; it was natural to include cyclic e^-^ flow as a fifth process in models of photosynthesis (*J_CEF_* ∼ *Φ_CEF_*).

These additional constraints on autotrophy can only decrease **λ**_max_. The effect on **λ**_max_ depends on the *C_red_* concentration via the dilution term **λ**[*C_red_*] (Fig. S11), but this additional constraint reduces autotrophic **λ**_max_ values even when [*C_red_*] is negligible because *J_red_* ≈ *J_ana_* is still required in this limit (see Fig. S13 for a geometric explanation). To estimate the magnitude of this effect on **λ**_max_ we randomly sampled parameters to construct comparable models of photosynthesis and respiration, observing a roughly threefold reduction in **λ**_max_ (Fig. 3B), with the exact difference depending on the sampling procedure. Our results therefore indicate that constraints absent from heterotrophy could explain why the fastest-growing autotrophs have longer generation times than the fastest heterotrophs (Fig. 3C), even when grown in nutrient surfeit (*37–39*).

### Energy economy impacts the redox state of proteins

Above we saw that large differences between biomass and nutrient redox state (|*Z_C,B_* - *Z_C,red_*|) challenge heterotrophic growth (Fig 2B-D), raising the prospect that microbes may adapt the composition of their biomass over physiological and evolutionary timescales to reduce the magnitude of the biochemical task. Fig. 2C gave us a sense of the expected direction of adaptation: in nutrient surplus, microbial growth is improved by making biomass with *Z_C,B_* similar to *Z_C,red_*.

Figure 4A illustrates how the maximum growth rate, *λ*_max_, varies as a function of *Z_C,B_* and *Z_C,red_*. The largest values fall just above the line of equality (*Z_C,B_* = *Z_C,red_*), indicating that the **λ**-maximizing *Z_C,B_* value is somewhat more oxidized than the C source. This is due to the need to use a fraction of the e^-^ from C substrates to produce ATP. The strength of this effect depends intracellular fluxes *J_α_* and, therefore, on all model parameters. The optimal *Z_C,B_* can be expressed as

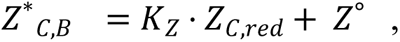

**Figure 4:**
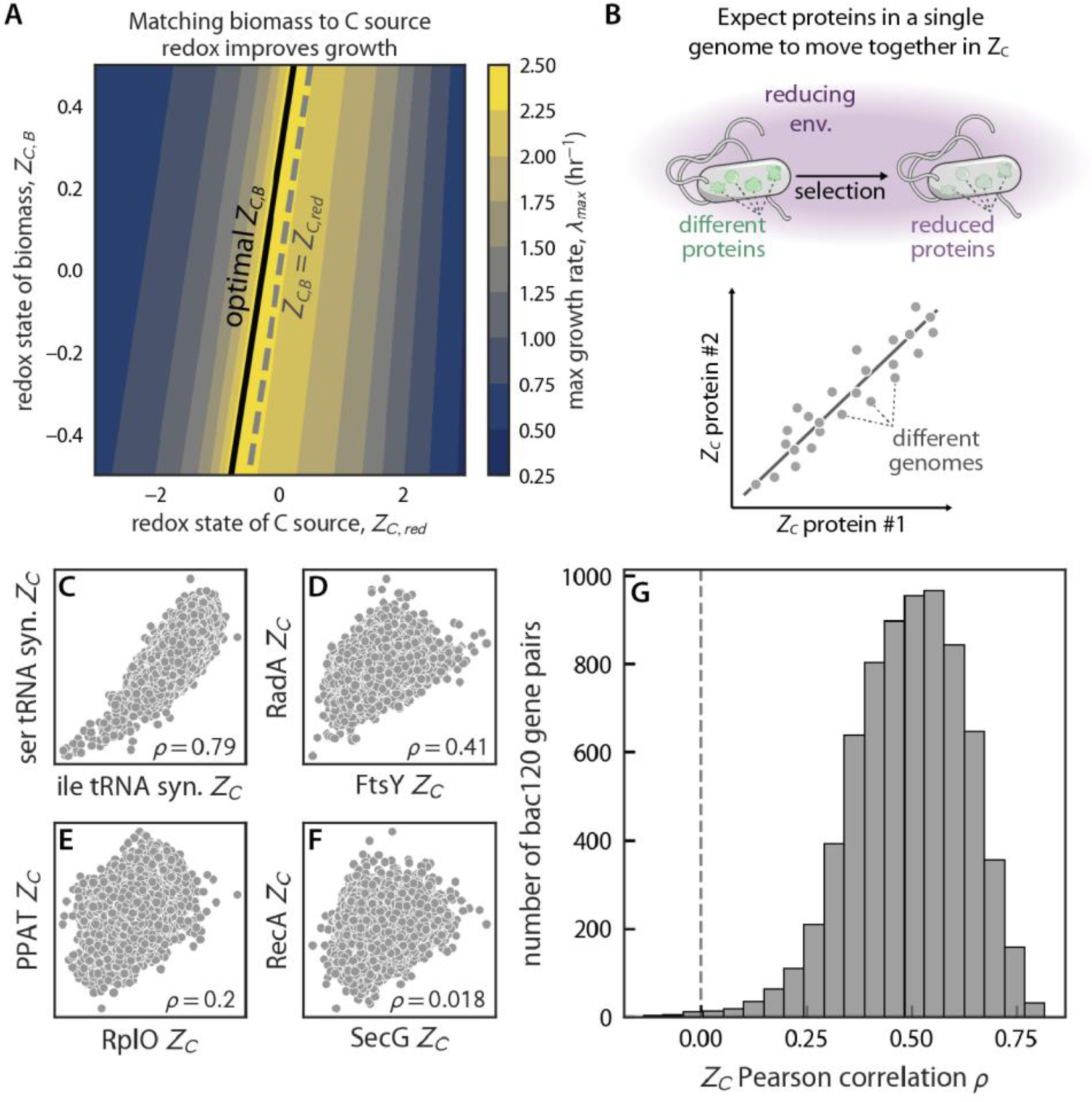
Z*_C_* values of conserved proteins are positively correlated, highlighting a redox-driven mode of protein evolution. (A) Our model indicated that heterotrophic growth can be improved by matching the redox state of biomass (*Z_C,B_*) to the carbon source (*Z_C,red_*). The dashed grey line marks *Z_C,B_* = *Z_C,red_* and the growth rate is maximized (black line) when the C source is somewhat more reduced than biomass. (B) If microbes are found in particular redox environments and/or have preferred metabolisms, we expect their biomass and, therefore, their proteins to take on the redox character of their environments. For example, organisms eating reduced C sources should benefit from making more reduced proteins. Proteins in the same genome should therefore exhibit positively correlated *Z_C_* values. (C-F) *Z_C_* correlations for pairs of conserved single-copy genes across ≈60,000 genomes spanning the bacterial tree of life. Sequences are drawn from the bac120 gene set (*50*). (G) Raw Pearson correlations were almost uniformly positive, with an interquartile range IQR = +0.41-0.59. Controlling for the mean *Z_C_* of coding sequences in each genome eliminated much of this correlation (partial correlation IQR = +0.07-0.21, Fig. S15), indicating that most of the observed effect is genome-wide and not specific to bac120 genes. Gene names: ser tRNA syn (serine tRNA synthetase), ile tRNA syn (isoleucine tRNA synthetase), FtsY (signal recognition particle-docking protein), RadA (DNA repair protein), RplO (ribosomal protein uL15), PPAT (pantetheine-phosphate adenylyltransferase), SecG (preprotein translocase subunit), RecA (DNA repair protein).

where the slope, *K_Z_*, and intercept, *Z°*, are functions of all model parameters and typical *K_Z_* values are positive. We interpret the right-hand side of this equation as an effective environmental redox potential, one that includes C source redox state, the redox potential of the e^-^ acceptor (via the reductive ATP yield, *S_3_*), and the magnitudes of intracellular fluxes. We also noted that the penalty of deviating from *Z\** by Δz is approximately quadratic in Δz (Figs. 4A and S9). So adapting *Z_C,B_* to the local environment is desirable — it reduces excess expenditures of energy and catalytic capacity, maximizing *λ* in rich environments.

If microbial lineages have characteristic habitats — environments with typical *Z\** values — and experience surplus with sufficient frequency, we expect evolutionary selection to push *Z_C,B_* towards *Z\**. Yet microbes frequently experience growth-limiting nutrient and/or energy deprivation in the wild — whether in soils (*40*, *41*), sediments (*42*, *43*) or elsewhere — so it is worth clarifying why we nonetheless assume selection for faster growth in rich conditions. Evolutionary selection and drift proceed over generations, not over absolute time (*44*). If microbes episodically experience surplus (e.g. “feast or famine” dynamics (*45*)) fast growth can account for a sizable fraction of generations even if it represents a minuscule fraction of the time window considered (*46*). As such, selection for increased maximum growth rate is fully consistent with the widespread limitation of microbial growth.

We therefore hypothesized that microbes’ biomass would evolve to match their environments (*47*), with biomass becoming more reduced when *Z_C,B_* - Z° > *K_Z_ · Z_C,red_* (Fig. 4B) or more oxidized when *Z_C,B_* - Z° < *K_Z_ · Z_C,red_*. Proteins, for example, can typically accept a variety of amino acid substitutions while still folding to achieve similar structures and biochemical activities (*48*, *49*). Indeed, across the tree of life, aminoacyl tRNA synthetases span nearly as wide a range of *Z_C_* values as all globular proteins in the *E. coli* genome (Fig. S13). Amino acid substitutions that move *Z_C,B_* towards optimality should therefore be both permitted and favored, for example substituting arginine (*Z_C_* = +⅓) with lysine (*Z_C_* = -⅔) in reducing conditions. As microbial genomes encode thousands of proteins, a generic selection pressure on *Z_C,B_* should affect all proteins, promoting substitutions moving in the same direction in *Z_C_* units (whether oxidizing or reducing) and resulting in positive correlations between *Z_C_* values of proteins in the same genome (Fig. 4B).

Importantly, while our analysis presumes balanced growth — i.e. time-invariant intracellular concentrations — we do not require chemical equilibrium. Rather, our model describes intracellular processes (oxidation, reduction, and anabolism) that must all carry net forward flux for growth to occur. Directional fluxes are a hallmark of non-equilibrium systems. In contrast with equilibrium descriptions (*47*), the magnitudes of these intracellular fluxes alter the optimal biomass redox state (*Z\**) in our model.

### Positive correlations in protein C redox state across the tree of life

To test our hypothesis, we examined a set of ≈60,000 genomes from across the bacterial tree curated by the Genome Taxonomy Database (GTDB, (*50*)). GTDB also identifies sequences for 120 nearly-universal single copy genes in the *bac120* gene set. Using single copy genes enabled our comparison of protein *Z_C_* values without the ambiguity arising from paralogs, e.g. genes with shared ancestry that have evolved different functions. We found that Pearson correlations (*ρ*) between *bac120 Z_C_* values were strongly biased in the positive direction, with an interquartile range (IQR) of +0.41-0.59 and a 99% confidence interval (CI) on the mean of +0.491-0.499 (Fig. 3C-F).

Given their ubiquity, it is not surprising that *bac120* genes perform essential functions like transcription (6 genes), translation (70), genome replication (18), and protein secretion (6). So, in addition to recording the influence of environmental redox conditions on coding sequences, positive correlations might also reflect the prominent and mostly anabolic roles *bac120* genes play in all bacteria. Indeed, expression of many anabolic proteins — especially those involved in protein synthesis — correlates positively with **λ** (*51–54*). Because increasing ribosome content results in the relative oxidation of biomass (Fig. 1C), one might expect such anabolic enzymes to accrue reducing amino acid substitutions that compensate for oxidation, thereby preserving *Z_C,B_* ≈ *K_Z_ ·Z_C,red_* - Z° as **λ** changes.

While both explanations are compatible with our model, we attempted to control for environmental effects, which should affect the entire genome, by calculating partial correlation coefficients controlling for the mean *Z_C_* of protein coding sequences in each genome. Partial correlations of *bac120 Z_C_* values were attenuated, but remained significantly biased towards positive values (IQR = +0.07-0.21, 99% CI = +0.135-0.141, Fig. S16), indicating that both physiology and environment affect the redox state of protein carbon.

### Redox driven protein sequence evolution

Proteins are typically considered to evolve towards improved individual function so long as the organism benefits from the improvement (*48*, *49*, *55*). Based on the correlations documented in Fig. 4F, we propose an additional mode of protein evolution wherein mutations neutral or even weakly deleterious to individual proteins’ biochemical or structural function might be selected due to an organismal benefit — they cause *Z_C,B_* to better align with the redox chemistry of the host’s typical environment (*47*). The affected proteins can be involved in any cellular structure or biochemical process, but large and highly-expressed proteins will affect *Z_C,B_* more.

Substantial changes to amino acid *Z_C_* often require more than one nucleotide substitution, however. To understand why this is, recall that the genetic code conserves hydrophobicity — single nucleotide substitutions tend to produce small changes in amino acid hydrophobicity metrics (*56*, *^57^*). Because hydrophobicity correlates with *Z_C_*, substitutions altering amino acid *Z_C_* also typically require multiple nucleotide changes (Fig. S17). Put differently, the rate of mutations affecting protein *Z_C_* is likely lower than the average nucleotide substitution rate, and so larger selection coefficients are needed to drive these sequence changes to fixation (*44*).

### Faster growth induces coordinated shifts in the redox state of proteomes

One principle arising from resource allocation logic is that, in conditions of surplus, *λ* is often limited by protein translation (*5*, *6*, *58*). Even in nutrient limited conditions, when faster growth requires greater supply of a nutrient like nitrogen, higher *λ* typically coincides with greater ribosome activities and larger RNA:protein ratios (*59*, *60*). Since ribosomes are approximately two-thirds RNA by mass (*6*, *9–12*, *61*) and RNA is about 1e^-^/C more oxidized than protein (*4*, *13*), the need for translation to keep pace with growth entails a relative oxidation of biomass at faster growth rates (Fig. 1C).

We estimated the scale of these biomass redox (*Z_C,B_*) changes by noting that slower-growing *E. coli* cultures are ≈10% RNA and ≈60% protein by mass (*λ* ≈ 0.4 hr^-1^), while fast growing cultures (*λ* ≈ 2 hr^-1^) are ≈20% RNA and ≈40% protein (*12*). Assuming *Z_C_* ≈ +0.9 for RNA and ≈ -0.15 for protein (*13*), *Z_C,B_* must increase by ≈0.03 for each unit increase in *λ* (hr^-1^ units, Fig. 5A, see Methods for detailed calculation). This value may seem small, yet, as noted above, deviations from optimal *Z_C,B_* lead to quadratic decreases in *λ*_max_ (Fig. 2B). As the molecular composition of translation cycle catalysts (ribosomes, tRNAs) leads to biomass oxidation we predicted that the remainder of biomass — proteins, lipids, storage polymers — should shift by a similar degree, but in the direction of reduction, to compensate.

**Figure 5:**
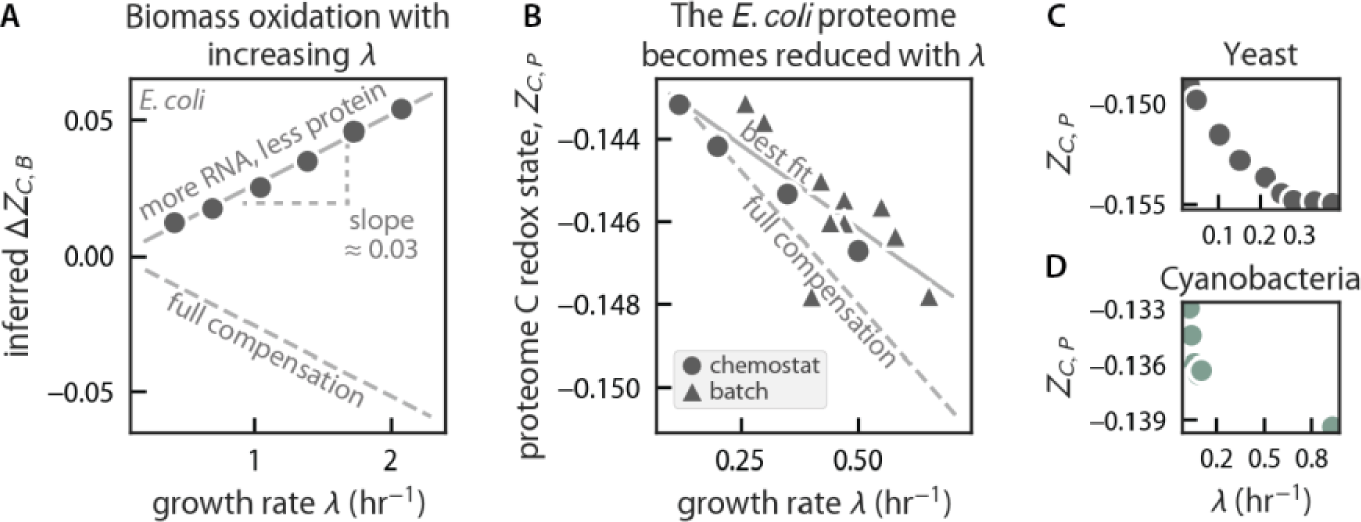
Microbial proteomes compensate for the oxidation of biomass by becoming reduced during fast growth. (A) Reflecting the composition of the ribosome, faster-growing cells contain more RNA and less protein. Given typical *Z_C_* values for biomass constituents (*13*), we estimated the changes in biomass redox state, Δ*Z_C,B_*, due to coordinated shifts in biomass composition with λ. This calculation implies a substantial oxidation of biomass of ≈0.03 *Z_C_* units per unit λ increase (hr^-1^ units), comparable to the standard deviation of all globular proteins in the *E. coli* genome (*σ* = 0.04 *Z_C_* units, Fig. S14A). Given the importance of *Z_C,B_* to attaining maximal growth rates in our model, we expected cells would compensate for this oxidation by making more reduced molecules during faster growth. (B) Quantitative proteomes from *E. coli* (*51*) demonstrate that the proteins produced during fast growth are more reduced. The observed reduction of the *E. coli* proteome is of the same order as, but falls short of complete compensation for (dashed line), background oxidation calculated in panel A. (E-F) Similar results are evident in the proteomes of model yeast (*53*) and cyanobacteria (*52*), suggesting this is a generic effect independent of metabolic mode or taxonomy.

In principle, *Z_C,B_* can be calculated from the elemental composition of biomass (*4*, *62*). While C:N:P ratios (called “Redfield ratios”) are commonly measured (*63*), accurate C:H:O:S ratios are also required to calculate *Z_C,B_* and these are challenging to measure. We noted, however, that quantitative proteomic surveys (*51–53*) report on the abundance and elemental composition of translated proteins and, therefore, the average redox state protein C, *Z_C,P_*. Consistent with our prediction, a comprehensive *E. coli* dataset (*51*) showed a systematic trend where the proteome was ≈0.005 units more reduced at the fastest measured growth rates (Fig. 5B). This work manipulated *λ* in chemostats and also by varying the C source in batch culture (*51*). These manipulations produced similar *λ*-*Z_C,P_* plots, suggesting that *λ*, rather than C source chemistry, is the primary determinant of *Z_C,P_*. Assuming that all biomass constituents participate equally in compensating for RNA-induced oxidation, we estimate that the relative reduction of *E. coli* protein C (≈ 0.01 *Z_C_* units per *λ* unit) compensates for ≈60% of the total effect.

To see if this observation is general, we examined additional *E. coli*, yeast (*53*) and Cyanobacterial proteomic datasets (*52*). Though these microbes perform highly distinct metabolisms and are separated by billions of years of evolutionary divergence (*64*), all datasets displayed *λ*-dependent *Z_C,P_* changes of the same direction and magnitude (Figs. 4C-D and S17). This effect appeared to be diffuse and widely shared across the proteome because explaining *Z_C,P_* variance required a linear model tracking multiple groups of proteins with at least 5 distinct biological functions (Fig. S19). Furthermore, *Z_C,P_* values calculated from expressed proteins differed significantly from those estimated from coding sequences (Fig. S20), indicating that (i) regulation of gene expression substantially affects the redox state of protein C and (ii) genomes and environmental metagenomes alone provide an incomplete picture of proteome redox states (*13*, *47*, *62*).

These observations add to a multi-decadal literature documenting *λ*-dependent changes in microbial form and composition. In *E. coli*, for example, fast growing cells are larger (*10*), have greater mass (*65*), lower surface area to volume ratios (*66*), contain more ribosomes and, consequently, more RNA, less protein and less lipid as a fraction of total biomass (*6*, *10–12*). Similar trends are documented for other microbes, especially yeast (*59*). These large-scale rearrangements can be rationalized via resource allocation logic (*5*, *6*, *8*, *10*) and all lead to biomass becoming more oxidized (e^-^ poor) during fast growth.

Beyond the observed changes in the redox state of protein C, we know of only one other example of biomass constituents becoming more reduced (e^-^ rich) during fast growth: the relative reduction of lipids with increased temperature or *λ* (*67–69*). Over the past two decades, several redox-based lipid biomarkers (e.g. TEX_86_) have been empirically established as paleothermometers and are now widely used to infer ancient sea surface temperatures and climate over the past 100 million years of Earth history. The mechanisms controlling lipid reduction remain debated, but recent work indicates that lipid biomarker chemistry is primarily controlled by growth rate (*68*). Our integrated model explains this relationship as optimizing microbial growth by compensating for the relative oxidation of biomass due to production of ribosomes during faster growth. In Fig. S21 we estimate the *Z_C_* of total *E. coli* lipids (*67*), finding *λ*-dependent changes of similar magnitude to the proteome, suggesting a common mechanism.

## Discussion

We developed a quantitative physiological model of microbial metabolic flexibility (Fig. 1) — microbes’ ability to extract energy and grow through a diversity of coupled redox processes — and explored the implications of our models’ results. This approach bridges the chemical environment — the nutrients and energy sources are available — with intracellular physiology, encoding both the chemical basis of bioenergetics and the simple fact that biomass is both a product and catalyst of cellular metabolism.

Prior efforts to integrate redox and resource allocation logic relied on genome-scale metabolic models (*21*, *22*). While these networks encode bioenergetic principles in a redox framework, it is challenging to extract principles from models describing thousands of enzymes, metabolites, and reactions. Below we review the lessons learned from our compact model, and, where appropriate, compare with those gleaned from other approaches.

Sustained growth at any rate requires the simultaneous balancing of the flows of energy (ATP), reductant (e^-^ carried on ECH), and matter (e.g. C atoms). Cells balance these fluxes through regulatory mechanisms that modulate intracellular activities, for example by inhibiting or activating ribosomes (*8*). Through our model, we found that microbes can regulate several aspects of their physiology to balance fluxes and grow in a diversity of settings. For example, in nutrient-limited conditions (e.g. lacking nitrogen or sulfur) microbes can balance their metabolism by accumulating C atoms on storage compounds like glycogen (*Φ_O_* ∼ 1/*λ* in Fig. 2A). Consistent with this prediction, microbes typically accumulate C storage molecules during nutrient-limited growth — e.g. glycogen and polyhydroxybutyrate in bacteria (*23–26*), lipid bodies in green algae (*27*, *28*) — with the amount proportional to the inverse growth rate 1/*λ*.

As the language of redox chemistry is generic, our model can represent any metabolism (Fig. S2). This enabled our comparison of heterotrophy and autotrophy to probe why the fastest known heterotrophs grow so much faster than similarly-distinguished autotrophs (Fig. 3). We showed that this discrepancy arises because autotrophs produce reduced carbon (*C_red_*) intracellularly. Intracellular flux balance (i.e. mass conservation) forces autotrophs to align production and utilization of *C_red_*, which substantially reduces *λ*_max_ (Fig. S13). This constraint does not apply to heterotrophs because they can deplete environmentally-supplied *C_red_*. Indeed, photoheterotrophic growth of green algae — a mixed metabolism where cells photosynthesize and consume extracellular organics at the same time — is notably faster than growth by photosynthesis alone (*39*, *70*). It appears, therefore, that the requirement to balance intracellularly-produced *C_red_* limits growth rates both in theory and in nature.

Metabolically-versatile microbes face a particular challenge: they must grow while consuming nutrients that are distinct from their biomass in redox state, i.e. with *Z_C,red_* ≠ *Z_C,B_*. Through our model, we discovered that cellular metabolism can be arranged to match a particular environment via modulation of stoichiometries and, thereby, *Z_C,B_* (Figs. 2B-D and 4A). Exact matching maximizes the growth rate, but large mismatches make growth infeasible because it becomes impossible to achieve simultaneous balancing of C, e^-^, and ATP flows. This led us to posit analogs of cyclic e^-^ flow for heterotrophy — regulated processes that bring supply of ATP and e^-^ supply fluxes in balance with anabolic demand. Introduction of such a process (regulated ATP hydrolysis) enabled growth across a broad range of *Z_C,red_* and *Z_C,B_* values.

Microbes can achieve flux balance in a variety of ways. *E. coli*, for example, can modulate ATP yields by performing alternative glycolytic pathways (*34*), altering the composition of its respiratory chain (*35*, *36*), or by secreting acetate instead of making CO_2_ (*7*). Genome-scale metabolic networks encode this flexibility by representing multiple pathways of C source oxidation and acceptor reduction with different products and ATP yields (*71*, *72*). Moreover, genome-scale models often allow excess maintenance ATP consumption (*b* in our model), implicitly enabling balancing by ATP hydrolysis. Notably, these degrees of freedom appear redundant: in a recent *E. coli* model (*71*) pathway switching is sufficient to achieve maximal growth rates across a wide variety of C sources and e^-^ acceptors (Fig. S3).

Regulated mechanisms of flux balancing can enable growth on a wide range of C sources, however these mechanisms come at a cost, consuming energy and requiring that cells dedicate resources to the synthesis of enzymes and regulators. Reducing this expenditure by generating biomass that matches the environment produced faster maximum growth rates (Fig. 4A), which led us to posit the existence of a generic selection pressure for microbes to match the redox state of biomass (*Z_C,B_*) to the chemical character of their habitats (*47*). Consistent with our hypothesis, we found strong and widespread positive correlations between the *Z_C_* values of conserved proteins (Fig. 3C-G), revealing that proteins are subject to shared evolutionary pressures along a compositional dimension not directly related to their individual functions. Amino acid substitutions that alter C redox are not small and it takes substantial evolutionary work (i.e. nucleotide substitutions) to affect such changes (Fig. S17). Yet we nonetheless observed diffuse accumulation of such changes throughout microbial proteomes. Additional support for this unexpected mode of protein evolution is given by our finding that, across diverse metabolic modes, microbes express comparatively reduced proteins during fast growth (Fig. 5), apparently using the proteome as an e^-^ acceptor to compensates for the relative oxidation of biomass due to the RNA content of ribosomes.

All of our results here derive from having integrated two potent, but disparate, views of cellular physiology — merging redox with resource allocation logic. This integration couples intracellular processes — regulatory, structural, and catalytic — to the extracellular redox environment, whether in a lake, an animal gut, a tumor matrix, or a soil pore. How should the character of the chemical environment affect metabolism and physiology? Our integrated, quantitative approach opens the door to asking and answering such questions across domains of life.

## Supporting information

Supplementary Table 1 -- survey of growth capabilities and rates.

Supplementary Table 2 -- chemical properties of amino acids.

Supplementary Table 3 -- GTDB bac120 sequences and Z_C values.

Supplementary Table 4 -- correlations between bac120 genes.

Supplementary Table 5 -- protein C redox states calculated from proteomics.

Supplementary Table 6 -- E. coli total lipid redox analysis.

## Acknowledgements

We are grateful to E. Afik, L. Aristilde, J. Ciemniecki, A. Duarte, J. Goldford, S. Hirokawa, R. Murali, T. Roeschinger, and G. Salmon for useful discussions and to Y.M. Bar-On, G. Chure, D. LaRowe, and R. Milo for detailed comments on the manuscript.

## Funding

NSF PHY-1748958 to the Kavli Institute for Theoretical Physics (AIF, AG)

Jane Coffin Childs Memorial Fund postdoctoral fellowship 61-1772 (AIF).

Gordon and Betty Moore Foundation grant GBMF4513 (AG)

Govt of India’s Ramalingaswami Fellowship (AG).

Caltech Center for Evolutionary Sciences (WWF)

National Institutes of Health grant 1R01AI127850-01A1 (DKN)

The Rosen Center at Caltech (RP)

National Institutes of Health MIRA grant 1R35 GM118043 (RP)

## Author contributions

Conceptualization: AIF

Methodology: AIF, AG

Investigation: AIF, AG, WWF, DKN, RP

Visualization: AIF, AG

Funding acquisition: AIF, AG, WWF, DKN, RP

Supervision: WWF, DKN, RP

Writing – original draft: AIF

Writing – review & editing: AIF, AG, WWF, DKN, RP

## Data and materials availability

All code and data are available either as supplementary tables or at https://github.com/flamholz/redox-proteome.

## References

1. C. G. Friedrich, B. Friedrich, B. Bowien, Formation of enzymes of autotrophic metabolism during heterotrophic growth of Alcaligenes eutrophus. Microbiology. 122, 69–78 (1981).

2. S. Gleizer, R. Ben-Nissan, Y. M. Bar-On, N. Antonovsky, E. Noor, Y. Zohar, G. Jona, E. Krieger, M. Shamshoum, A. Bar-Even, R. Milo, Conversion of Escherichia coli to Generate All Biomass Carbon from CO2. Cell. 179, 1255–1263.e12 (2019).

3. F. M. Harold, The vital force: a study of bioenergetics (1987), (available at https://core.ac.uk/download/pdf/82700792.pdf).

4. D. E. LaRowe, P. Van Cappellen, Degradation of natural organic matter: A thermodynamic analysis. Geochim. Cosmochim. Acta. 75, 2030–2042 (2011).

5. D. Molenaar, R. van Berlo, D. de Ridder, B. Teusink, Shifts in growth strategies reflect tradeoffs in cellular economics. Mol. Syst. Biol. 5, 323 (2009).

6. M. Scott, C. W. Gunderson, E. M. Mateescu, Z. Zhang, T. Hwa, Interdependence of cell growth and gene expression: origins and consequences. Science. 330, 1099–1102 (2010).

7. M. Basan, S. Hui, H. Okano, Z. Zhang, Y. Shen, J. R. Williamson, T. Hwa, Overflow metabolism in Escherichia coli results from efficient proteome allocation. Nature. 528, 99– 104 (2015).

8. G. Chure, J. Cremer, An optimal regulation of fluxes dictates microbial growth in and out of steady state. Elife. 12 (2023), doi:10.7554/eLife.84878.

9. S. Reuveni, M. Ehrenberg, J. Paulsson, Ribosomes are optimized for autocatalytic production. Nature. 547, 293–297 (2017).

10. M. Schaechter, O. Maaloe, N. O. Kjeldgaard, Dependency on medium and temperature of cell size and chemical composition during balanced grown of Salmonella typhimurium. J. Gen. Microbiol. 19, 592–606 (1958).

11. F. C. Neidhardt, B. Magasanik, Studies on the role of ribonucleic acid in the growth of bacteria. Biochim. Biophys. Acta. 42, 99–116 (1960).

12. H. Bremer, P. P. Dennis, Modulation of Chemical Composition and Other Parameters of the Cell at Different Exponential Growth Rates. EcoSal Plus. 3, 1–49 (2008).

13. J. M. Dick, M. Yu, J. Tan, A. Lu, Changes in Carbon Oxidation State of Metagenomes Along Geochemical Redox Gradients. Front. Microbiol. 10, 120 (2019).

14. P. G. Falkowski, T. Fenchel, E. F. Delong, The microbial engines that drive Earth’s biogeochemical cycles. Science. 320, 1034–1039 (2008).

15. Z. Shi, S. Crowell, Y. Luo, B. Moore 3rd, Model structures amplify uncertainty in predicted soil carbon responses to climate change. Nat. Commun. 9, 2171 (2018).

16. P. Mitchell, Chemiosmotic coupling in oxidative and photosynthetic phosphorylation. Biol. Rev. Camb. Philos. Soc. 41, 445–502 (1966).

17. S. Manzoni, P. Taylor, A. Richter, A. Porporato, G. I. Ågren, Environmental and stoichiometric controls on microbial carbon-use efficiency in soils. New Phytol. 196, 79–91 (2012).

18. R. L. Sinsabaugh, S. Manzoni, D. L. Moorhead, A. Richter, Carbon use efficiency of microbial communities: stoichiometry, methodology and modelling. Ecol. Lett. 16, 930–939 (2013).

19. E. J. Zakem, M. F. Polz, M. J. Follows, Redox-informed models of global biogeochemical cycles. Nat. Commun. 11, 5680 (2020).

20. F. Tao, Y. Huang, B. A. Hungate, S. Manzoni, S. D. Frey, M. W. I. Schmidt, M. Reichstein, N. Carvalhais, P. Ciais, L. Jiang, J. Lehmann, Y.-P. Wang, B. Z. Houlton, B. Ahrens, U. Mishra, G. Hugelius, T. D. Hocking, X. Lu, Z. Shi, K. Viatkin, R. Vargas, Y. Yigini, C. Omuto, A. A. Malik, G. Peralta, R. Cuevas-Corona, L. E. Di Paolo, I. Luotto, C. Liao, Y.-S. Liang, V. S. Saynes, X. Huang, Y. Luo, Microbial carbon use efficiency promotes global soil carbon storage. Nature, 1–5 (2023).

21. A. Goelzer, J. Muntel, V. Chubukov, M. Jules, E. Prestel, R. Nölker, M. Mariadassou, S. Aymerich, M. Hecker, P. Noirot, D. Becher, V. Fromion, Quantitative prediction of genome-wide resource allocation in bacteria. Metab. Eng. 32, 232–243 (2015).

22. M. Mori, T. Hwa, O. C. Martin, A. De Martino, E. Marinari, Constrained Allocation Flux Balance Analysis. PLoS Comput. Biol. 12, e1004913 (2016).

23. J. Preiss, Bacterial glycogen synthesis and its regulation. Annu. Rev. Microbiol. 38, 419– 458 (1984).

24. T. Holme, G. Westöö, L. Svennerholm, A. Magnéli, A. Magnéli, H. Pestmalis, S. Åsbrink, Continuous culture studies on glycogen synthesis in Escherichia coli B. Acta Chem. Scand. 11, 763–775 (1957).

25. H. Lees, J. R. Postgate, The behaviour of Azotobacter chroococcum in oxygen- and phosphate-limited chemostat culture. J. Gen. Microbiol. 75, 161–166 (1973).

26. J. D. Linton, R. E. Cripps, The occurrence and identification of intracellular polyglucose storage granules in Methylococcus NCIB 11083 grown in chemostat culture on methane. Arch. Microbiol. 117, 41–48 (1978).

27. Z. T. Wang, N. Ullrich, S. Joo, S. Waffenschmidt, U. Goodenough, Algal lipid bodies: stress induction, purification, and biochemical characterization in wild-type and starchless Chlamydomonas reinhardtii. Eukaryot. Cell. 8, 1856–1868 (2009).

28. A. J. Klok, D. E. Martens, R. H. Wijffels, P. P. Lamers, Simultaneous growth and neutral lipid accumulation in microalgae. Bioresour. Technol. 134, 233–243 (2013).

29. D. I. Arnon, Conversion of light into chemical energy in photosynthesis. Nature. 184, 10– 21 (1959).

30. A. Burlacot, Quantifying the roles of algal photosynthetic electron pathways: a milestone towards photosynthetic robustness. New Phytol. 240, 2197–2203 (2023).

31. J. B. McKinlay, C. S. Harwood, Carbon dioxide fixation as a central redox cofactor recycling mechanism in bacteria. Proc. Natl. Acad. Sci. U. S. A. 107, 11669–11675 (2010).

32. W.-H. Lin, C. Jacobs-Wagner, Connecting single-cell ATP dynamics to overflow metabolism, cell growth, and the cell cycle in Escherichia coli. Curr. Biol. 32, 3911– 3924.e4 (2022).

33. L. Gómez-Consarnau, J. A. Raven, N. M. Levine, L. S. Cutter, D. Wang, B. Seegers, J. Arístegui, J. A. Fuhrman, J. M. Gasol, S. A. Sañudo-Wilhelmy, Microbial rhodopsins are major contributors to the solar energy captured in the sea. Sci Adv. 5, eaaw8855 (2019).

34. A. Flamholz, E. Noor, A. Bar-Even, W. Liebermeister, R. Milo, Glycolytic strategy as a tradeoff between energy yield and protein cost. Proceedings of the National Academy of Sciences. 110, 10039–10044 (2013).

35. M. Bekker, S. de Vries, A. Ter Beek, K. J. Hellingwerf, M. J. T. de Mattos, Respiration of Escherichia coli can be fully uncoupled via the nonelectrogenic terminal cytochrome bd-II oxidase. J. Bacteriol. 191, 5510–5517 (2009).

36. V. B. Borisov, R. Murali, M. L. Verkhovskaya, D. A. Bloch, H. Han, R. B. Gennis, M. I. Verkhovsky, Aerobic respiratory chain of *Escherichia coli* is not allowed to work in fully uncoupled mode. Proc. Natl. Acad. Sci. U. S. A. 108, 17320–17324 (2011).

37. H. W. Jannasch, C. O. Wirsen, D. C. Nelson, L. A. Robertson, Thiomicrospira crunogena sp. nov., a Colorless, Sulfur-Oxidizing Bacterium from a Deep-Sea Hydrothermal Vent. Int. J. Syst. Bacteriol. 35, 422–424 (1985).

38. D. Jaiswal, A. Sengupta, S. Sohoni, S. Sengupta, A. G. Phadnavis, H. B. Pakrasi, P. P. Wangikar, Genome Features and Biochemical Characteristics of a Robust, Fast Growing and Naturally Transformable Cyanobacterium Synechococcus elongatus PCC 11801 Isolated from India. Sci. Rep. 8, 16632 (2018).

39. H. Treves, O. Murik, I. Kedem, D. Eisenstadt, S. Meir, I. Rogachev, J. Szymanski, N. Keren, I. Orf, A. F. Tiburcio, R. Alcázar, A. Aharoni, J. Kopka, A. Kaplan, Metabolic Flexibility Underpins Growth Capabilities of the Fastest Growing Alga. Curr. Biol. 27, 2559–2567.e3 (2017).

40. T. A. Caro, J. McFarlin, S. Jech, N. Fierer, S. Kopf, Hydrogen stable isotope probing of lipids demonstrates slow rates of microbial growth in soil. Proc. Natl. Acad. Sci. U. S. A. 120, e2211625120 (2023).

41. S. J. Blazewicz, B. A. Hungate, B. J. Koch, E. E. Nuccio, E. Morrissey, E. L. Brodie, E. Schwartz, J. Pett-Ridge, M. K. Firestone, Taxon-specific microbial growth and mortality patterns reveal distinct temporal population responses to rewetting in a California grassland soil. ISME J. 14, 1520–1532 (2020).

42. M. A. Lever, K. L. Rogers, K. G. Lloyd, J. Overmann, B. Schink, R. K. Thauer, T. M. Hoehler, B. B. Jørgensen, Life under extreme energy limitation: a synthesis of laboratory- and field-based investigations. FEMS Microbiol. Rev. 39, 688–728 (2015).

43. T. M. Hoehler, B. B. Jørgensen, Microbial life under extreme energy limitation. Nat. Rev. Microbiol. 11, 83–94 (2013).

44. T. Ohta, The Nearly Neutral Theory of Molecular Evolution. Annu. Rev. Ecol. Syst. 23, 263–286 (1992).

45. J. Merritt, S. Kuehn, Frequency- and Amplitude-Dependent Microbial Population Dynamics during Cycles of Feast and Famine. Phys. Rev. Lett. 121, 098101 (2018).

46. F. J. Bruggeman, B. Teusink, R. Steuer, Trade-offs between the instantaneous growth rate and long-term fitness: Consequences for microbial physiology and predictive computational models. Bioessays. 45, e2300015 (2023).

47. J. M. Dick, D. Meng, Community- and genome-based evidence for a shaping influence of redox potential on bacterial protein evolution. mSystems, e0001423 (2023).

48. M. A. DePristo, D. M. Weinreich, D. L. Hartl, Missense meanderings in sequence space: a biophysical view of protein evolution. Nat. Rev. Genet. 6, 678–687 (2005).

49. M. Soskine, D. S. Tawfik, Mutational effects and the evolution of new protein functions. Nat. Rev. Genet. 11, 572–582 (2010).

50. D. H. Parks, M. Chuvochina, P.-A. Chaumeil, C. Rinke, A. J. Mussig, P. Hugenholtz, A complete domain-to-species taxonomy for Bacteria and Archaea. Nat. Biotechnol. (2020), doi:10.1038/s41587-020-0501-8.

51. A. Schmidt, K. Kochanowski, S. Vedelaar, E. Ahrné, B. Volkmer, L. Callipo, K. Knoops, M. Bauer, R. Aebersold, M. Heinemann, The quantitative and condition-dependent Escherichia coli proteome. Nat. Biotechnol. 34, 104–110 (2016).

52. T. Zavřel, M. Faizi, C. Loureiro, G. Poschmann, K. Stühler, M. Sinetova, A. Zorina, R. Steuer, J. Červený, Quantitative insights into the cyanobacterial cell economy. Elife. 8 (2019), doi:10.7554/eLife.42508.

53. J. Xia, B. J. Sánchez, Y. Chen, K. Campbell, S. Kasvandik, J. Nielsen, Proteome allocations change linearly with the specific growth rate of Saccharomyces cerevisiae under glucose limitation. Nat. Commun. 13, 2819 (2022).

54. R. Balakrishnan, M. Mori, I. Segota, Z. Zhang, R. Aebersold, C. Ludwig, T. Hwa, Principles of gene regulation quantitatively connect DNA to RNA and proteins in bacteria. Science. 378, eabk2066 (2022).

55. L. Noda-Garcia, W. Liebermeister, D. S. Tawfik, Metabolite–Enzyme Coevolution: From Single Enzymes to Metabolic Pathways and Networks. Annu. Rev. Biochem. 87, 187–216 (2018).

56. L. Shenhav, D. Zeevi, Resource conservation manifests in the genetic code. Science. 370, 683–687 (2020).

57. D. Haig, L. D. Hurst, A quantitative measure of error minimization in the genetic code. J. Mol. Evol. 33, 412–417 (1991).

58. N. M. Belliveau, G. Chure, C. L. Hueschen, H. G. Garcia, J. Kondev, D. S. Fisher, J. A. Theriot, R. Phillips, Fundamental limits on the rate of bacterial growth and their influence on proteomic composition. Cell Syst. 12, 924–944.e2 (2021).

59. E. Metzl-Raz, M. Kafri, G. Yaakov, I. Soifer, Y. Gurvich, N. Barkai, Principles of cellular resource allocation revealed by condition-dependent proteome profiling. Elife. 6 (2017), doi:10.7554/eLife.28034.

60. S. H.-J. Li, Z. Li, J. O. Park, C. G. King, J. D. Rabinowitz, N. S. Wingreen, Z. Gitai, Escherichia coli translation strategies differ across carbon, nitrogen and phosphorus limitation conditions. Nat. Microbiol. 3, 939–947 (2018).

61. V. Ramakrishnan, Ribosome structure and the mechanism of translation. Cell. 108, 557–572 (2002).

62. J. M. Dick, Average oxidation state of carbon in proteins. J. R. Soc. Interface. 11, 20131095 (2014).

63. C. C. Cleveland, D. Liptzin, C: N: P stoichiometry in soil: is there a “Redfield ratio” for the microbial biomass? Biogeochemistry. 85, 235–252 (2007).

64. S. Kumar, M. Suleski, J. M. Craig, A. E. Kasprowicz, M. Sanderford, M. Li, G. Stecher, S. B. Hedges, TimeTree 5: An Expanded Resource for Species Divergence Times. Mol. Biol. Evol. 39 (2022), doi:10.1093/molbev/msac174.

65. B. R. K. Roller, C. Hellerschmied, Y. Wu, T. P. Miettinen, A. L. Gomez, S. R. Manalis, M. F. Polz, Single-cell mass distributions reveal simple rules for achieving steady-state growth. MBio, e0158523 (2023).

66. L. K. Harris, J. A. Theriot, Surface Area to Volume Ratio: A Natural Variable for Bacterial Morphogenesis. Trends Microbiol. 26, 815–832 (2018).

67. A. G. Marr, J. L. Ingraham, Effect of temperature on the composition of fatty acids in Escherichia coli. J. Bacteriol. 84, 1260–1267 (1962).

68. S. J. Hurley, F. J. Elling, M. Könneke, C. Buchwald, S. D. Wankel, A. E. Santoro, J. S. Lipp, K.-U. Hinrichs, A. Pearson, Influence of ammonia oxidation rate on thaumarchaeal lipid composition and the TEX86 temperature proxy. Proc. Natl. Acad. Sci. U. S. A. 113, 7762–7767 (2016).

69. H. C. Holm, H. F. Fredricks, S. M. Bent, D. P. Lowenstein, J. E. Ossolinski, K. W. Becker, W. M. Johnson, K. Schrage, B. A. S. Van Mooy, Global ocean lipidomes show a universal relationship between temperature and lipid unsaturation. Science. 376, 1487–1491 (2022).

70. G. Lalibertè, J. de la Noüie, Auto-, hetero-, and mixotrophic growth of chlamydomonas Humicola (Chlorophyceae) on acetate. J. Phycol. 29, 612–620 (1993).

71. J. M. Monk, C. J. Lloyd, E. Brunk, N. Mih, A. Sastry, Z. King, R. Takeuchi, W. Nomura, Z. Zhang, H. Mori, A. M. Feist, B. O. Palsson, iML1515, a knowledgebase that computes Escherichia coli traits. Nat. Biotechnol. 35, 904–908 (2017).

72. W. Gottstein, B. G. Olivier, F. J. Bruggeman, B. Teusink, Constraint-based stoichiometric modelling from single organisms to microbial communities. J. R. Soc. Interface. 13 (2016), doi:10.1098/rsif.2016.0627.

