## Supplementary Text for "The proteome is a terminal electron acceptor"

### Supplementary Materials
### The proteome is a terminal electron acceptor

**Avi I. Flamholz**[1], **Akshit Goyal**[2,3], **Woodward W. Fischer**[4], **Dianne K. Newman**[1,4], and **Rob Phillips**[1,5]

[1]Division of Biology and Biological Engineering, California Institute of Technology, Pasadena, CA 91125
[2]Physics of Living Systems, Department of Physics, Massachusetts Institute of Technology, Cambridge 02139.
[3]International Centre for Theoretical Sciences, Tata Institute of Fundamental Research, Bengaluru 560089.
[4]Division of Geological and Planetary Sciences, California Institute of Technology, Pasadena, CA 91125
[5]Department of Physics, California Institute of Technology, Pasadena, CA 91125.
*Correspondence:

### List of Tables

1 Proteomics datasets and reference proteomes used in this study. . . . . . . . . . . . . . . . . . . . . . . . 5
2 Stoichiometric coefficients used in our metabolic models. . . . . . . . . . . . . . . . . . . . . . . . . . . . 18

### 1 Materials and Methods

#### 1.1 A chemical resource-allocation model of microbial growth.

We developed a chemical resource allocation model of microbial growth both for autotrophy and heterotrophy. To understand the behavior of this model we explored its analytic limits, testing limiting behavior with optimizations and simulations implemented in Python. Model equations, plausible ranges of parameter values, and simulations are detailed in the supplementary text below.

#### 1.2 Evaluation of *E. coli* metabolic range.

Figure 1A describes *E. coli*'s capacity to grow on pairs of (carbon source, $e^-$ acceptor). Supplementary Table 1 gives references supporting observations of growth in various conditions. To evaluate whether the *E. coli* genome contains the enzymes and pathways to grow on a given pair in principle, we performed flux balance analysis on a genome-scale model of *E. coli* metabolism, iML1515 (*1*), using the COBRApy package for Python. We provided the model with a minimal medium containing only the chosen carbon source and terminal acceptor, and a positive result was recorded when the predicted maximum growth rate exceeded 0.01 $hr^{-1}$. This analysis is summarized in Supplementary Table 1 and Fig. S3.

In some cases, we could not find experimental confirmation for positive model results e.g. predicted fermentative growth on L-asparagine as the sole source of carbon. This could indicated that (i) lab strains will grow fermentatively on L-asparagine if tested, (ii) that lab strains could do so after laboratory evolution (*2*) or (iii) that there is an error in the iML1515 model. In other cases, negative model predictions were contradicted by observations, e.g. of fermentative growth on L-cysteine (*3*), likely indicating some error in the model. We therefore suggest that experimental validation of predicted growth capabilities be more widely used in evaluating metabolic models.

#### 1.3 Estimating the redox effects of changes in macromolecular composition

We considered changes in the macromolecular composition of *E. coli* as a function of $\lambda$. The data supporting these plots was drawn from (*4*), which reports $\mu$g DNA, RNA and protein per cell as a function of $\lambda$ as well as total cell mass. We converted these values to mass fractions and calculated a residual mass fraction associated with other molecules, which are likely dominated by lipids and polysaccharides. In ref. (*4*) the growth rate is defined as $\mu = 1/\tau$, where $\tau$ is the doubling time, while in our model $\lambda = \ln(2)/\tau$. This explains the discrepancy between $\lambda$ values plotted here and $\mu$ values given in (*4*).

To convert $\lambda$-dependent biomass composition to a change in redox state, we estimated the C mass fraction and C redox state for DNA, RNA and protein. For protein we assumed the atomic formula of $C_{100}$ $H_{159}$ $N_{26}$ $O_{32}$ $S_{0.7}$ drawn from Bionumbers (BNID 109413) to estimate a C mass fraction of $\approx$ 53 %. For RNA ($\approx$ 31%) and DNA ($\approx$ 34%) we averaged the C mass fractions nucleotide monophosphates and deoxynucleotide monophosphoates respectively. For the residual we assumed a C mass fraction of 60% because of its lipid content and also so that the total C mass is $\approx$ 50% carbon. From Fig. S20 we see that $Z_C \approx -0.15$ for *E. coli* proteins. For DNA and RNA we took values at 50% GC content from Figure 1 of (*5*), namely $Z_C \approx +0.9$ for RNA and $Z_C \approx +0.6$ for DNA. While $Z_C$ is a direct function of GC content for double stranded DNA, (*5*) assumed equal representation of G and C and of A and U nucleotides to make an RNA estimate. Since we are calculating changes in $Z_{C,B}$ due to measured macromolecular components, we set the residual compartments' $Z_C$ to 0 and then took the C-weighted average of these $Z_C$ values in each condition to arrive at $Z_{C,B}$ estimates as a function of $\lambda$.

#### 1.4 Calculations of the C redox state for biological molecules

The C redox state of a molecule — its $Z_C$ value — is the average formal charge of carbon on that molecule. So the $Z_C$ of a protein is the C-weighted average of its constituent amino acids. $Z_C$ values for amino acids are given in Supplementary Table 2. In section 2 of the supplementary text we give detailed description of how these values are calculated in general with particular focus on proteins.

#### 1.5 Analysis of bac120 protein sequences from the Genome Taxonomy Database (GTDB).

Release 207 (r207) of the Genome Taxonomy Database (*6*) catalogs ≈300,000 prokaryotic genomes. The genomes are clustered by sequence identity and each cluster of closely related species is assigned a representative genome that is high-quality and, preferably, a type strain with a published name. r207 contained 62,291 representative genomes. GTDB performs bacterial phylogenetic analyses by comparing the concatenated sequences of 120 protein sequences that are nearly always found in single copy, the so-called "bac120" genes (*6*). We used these genes to ask if proteins encoded in the same genome have correlated $Z_C$ values, as predicted by our model. The nearly-universal and single-copy nature of bac120 genes is advantageous here because, for each gene, we can unambiguously associate a single sequence with each representative genome and, thereby, construct a vector of 120 $Z_C$ values describing every genome.

We downloaded the bac120 protein sequences as well whole genome sequences for all representative genomes from GTDB release 207. To assess the physiological roles of bac120 genes, we manually mapped each protein to a high-level functional category from the COG database (*7*). We then calculated $Z_C$ values for all annotated protein coding sequences in each representative genome, $\langle Z_C \rangle_G$, as described above. Raw correlations between bac120 $Z_C$ as well as partial correlations controlling for $\langle Z_C \rangle_G$ were calculated using the pingouin package for Python. $Z_C$ values of bac120 sequences are reported in Supplementary Table 3 while pairwise correlations are in Supplementary Table 4.

#### 1.6 Genetic code analyses.

The genetic code is known to be conservative for various amino acid properties, especially measures of hydrophobicity. As such, single mutations are unlikely to alter measures of hydrophobicity substantially (*8*, *9*). To assess whether the genetic code is also conservative for amino acid $Z_C$, we tested if (i) hydrophobicity indices are correlated with $Z_C$ (Fig. S17A-B) and (ii) how many mutations are required to alter $Z_C$ on average (Fig. S17C). For the former analysis we calculated correlations of amino acid $Z_C$ with polar requirement (roughly hydrophillicity) and hydropathy index (roughly hydrophobicity), drawing values for these properties from (*8*). For the latter analysis, we considered all possible pairs of codon substitutions, for example replacing CGA (arginine) with AAA (lysine). This example requires two nucleotide substitutions and results in a $Z_C$ change of $\Delta Z_C = -1$. We then binned the codon transitions by their $\Delta Z_C$ values and used linear regression to estimate the relationship between $\Delta Z_C$ and the required number of nucleotide substitutions (Fig. S17C).

#### 1.7 Reference proteomes for quantitative proteomics datasets

We considered quantitative proteomics data from *E. coli*, brewers yeast, and a model cyanobacteria. To retrieve metadata for each organism, we downloaded and parsed the relevant XML formatted "reference proteome" from the NCBI RefSeq database (Table 1). Not to be confused with the quantitative proteomes, "reference proteomes" list all known protein coding sequences in a genome along with metadata such as gene names, unique identifiers, functional annotations (e.g., KEGG (*10*) and COG (*7*) databases), and annotation of transmembrane segments (*11*). Secondary isoforms (e.g. due to splicing, translational slippage, etc.) were extracted from auxiliary files provided by RefSeq. Proteins with non-specific amino acid identifiers in their sequence were ignored. $N_C$ and $Z_C$ values were then calculated for each protein isoform from amino acid sequences as the sum and C-weighted average of amino acid values respectively (*12*, *13*). These values are reported in Supplementary Table 5.

| species and strain | NCBI RefSeq ID | Dataset References |
|---|---|---|
| *E. coli* MG1655 K12 | UP000000625 | (*14–16*) |
| *S. cerevisiae* S288c | UP000002311 | (*17*) |
| *Synechocystis* sp. PCC 6803 | UP000001425 | (*18*) |

**Table 1.** Proteomics datasets and NCBI RefSeq identifiers for reference proteomes used in this study.

#### 1.8 Quantitative proteomic data and calculation of whole proteome redox state, $Z_{C,P}$.

We use "proteome" to describe the mean expression levels of proteins in a microbial culture. Such values are typically measured by mass spectrometry, though sometimes by other means, e.g., ribosome profiling (*19*) or tagging individual proteins via genetic manipulation (*20*). Since the redox state of expressed proteins, $Z_{C,P}$, is a C-weighted average of all expressed proteins, only relative expression levels are required, not absolute values with real units (e.g. copies/cell, fg/gDW). Expression measurements from high-quality proteomics surveys of *E. coli*, *S. cerevisiae*, and the Cyanobacterium *Synechocystis* sp. PCC 6803 (Table 1) were mapped to reference proteomes via unique identifiers (protein accessions or *E. coli* b-numbers), verifying that all or nearly all measured protein C could be mapped to a specific protein with a known sequence. Proteome-wide $Z_{C,P}$ values were then calculated as the C-weighted average of expressed proteins

$$Z_{C,P} = \frac{\sum_i N_{C,i} \cdot Z_{C,i} \cdot \eta_i}{\sum_i N_{C,i} \cdot \eta_i}. \tag{1}$$

Here $N_{C,i}$ and $Z_{C,i}$ give the number of C atoms and the formal oxidation state of C in protein $i$ while $\eta_i$ gives the (relative or absolute) expression level of the same protein. $Z_{C,i}$ values are calculated from the amino acid sequence of each protein as described in section 2. Supplementary Table S5 gives a full listing of protein $Z_C$ values for all three model organisms, reproduces protein expression data and reports the derived $Z_{C,P}$ value for all experimental conditions considered here.

#### 1.9 Calculation of $Z_C$ using genomic coding sequences.

Recent work has investigated whether environmental redox conditions affect the average $Z_C$ of genomic coding sequences (*13*, *21*). We calculated the C-weighted average of coding sequence $Z_C$ values by setting $\eta_i = 1$ in equation 1, i.e., treating all proteins as if they are equally expressed. We term this value $\langle Z_C \rangle_G$. To estimate a confidence interval for $\langle Z_C \rangle_G$ we sampled 1000 coding sequences from each genome, repeating this procedure $10^4$ times. This sampling procedure attempts to replicate variation in $\langle Z_C \rangle_G$ that might occur due to inadequate sequencing depth, horizontal gene transfer, or errors in genome reconstruction. To estimate the uncertainty associated with unknown protein expression — i.e., $\eta_i$ are unknown when working with genomes — we performed the same sampling procedure, but sampled $\eta_i$ from a log-normal distribution with $\sigma = 2.3$ fit from *E. coli* expression data. This results in $\eta_i$ varying over roughly 6 orders of magnitude and permits a very wide range of $Z_{C,P}$ estimates (Fig. S20).

#### 1.10 Sparse reconstruction of $Z_{C,P}$ variation with LASSO regression

$Z_{C,P}$ is the C-weighted average of expressed proteins. Inspecting equation 1 we see that each protein $i$ contributes

$$x_{i,j} = \frac{N_{C,i} \cdot Z_{C,i} \cdot \eta_i^j}{\sum_i N_{C,i} \cdot \eta_i^j}$$

to the total $Z_{C,P}^j$, where $j$ is an index marking the experiment (e.g., glucose or succinate media). Since $Z_{C,P}^j = \sum_i x_{i,j}$ it follows that 100% of $Z_{C,P}$ variation across conditions $j$ can be accounted for by the matrix $X = [x_{i,j}]$. In other words, ordinary linear regression fitting $Z_{C,P}^j = \beta X$ would always recover a $\beta$ that perfectly reconstructs $Z_{C,P}^j$. Indeed $\beta = \vec{1}$ does this trivially.

To understand the dimensionality of the $Z_{C,P}$ trend, i.e. how many groups of proteins drive the trend, we asked: how many 'basis' proteins $i$ are required to reconstruct 100% of $Z_{C,P}$ variation? To achieve this, we used the Python sklearn implementation of LASSO regression (setting 'fit_intercept' to False), which minimizes a regularized loss function

$$\sum_j \left( Z_{C,P}^j - \beta_i x_{i,j} \right)^2 - \alpha \sum_i |\beta_i|$$

that penalizes solutions where entries of $\beta$ are large. The first term is the $l_2$ norm of residuals, while the second term imposes a penalty on the weights $\beta$. Large values of $\alpha$ prefer sparser solutions that are less accurate. By varying $\alpha$, we ask how many individual proteins are required to reconstruct a set of $Z^j_{C,P}$ values. Setting $\alpha = 10^{-8}$ produced near-perfect reconstructions of $Z_{C,P}$ trends in all three organisms (see Fig. S19). Because groups of proteins are often co-expressed — i.e., their expression levels are correlated — the proteins chosen by the LASSO regression may represent a larger set of proteins with correlated $x_{i,j}$ values. As such, we inspected COG functional categories associated with the chosen proteins (*7*) to determine if they perform similar or different biological functions (Fig. S19D).

##### 1.11 *E. coli* total lipid $Z_C$ analysis.

Figure S21 reports the growth-rate dependence of the C redox state of *E. coli* total lipids. The temperature-dependent composition of *E. coli* lipids was drawn from (*22*). The number of C atoms ($N_C$) and C redox state ($Z_C$) of individual lipids was manually calculated. Their measured proportions were then used to estimate the C redox state of *E. coli*lipid C, $Z_{C,L}$, as a function of temperature during *E. coli* growth in minimal glucose medium. Since ref. (*22*) did not report the growth rate $\lambda$, the relationship between temperature and $\lambda$ was drawn from (*23*). This reference grew *E. coli* in a similar media to (*22*), fitting

$$\sqrt{\lambda} = b\,(T - T_{min}) \cdot (1 - \exp\,(c \cdot (T - T_{max})))$$

with fit values $b = 0.0262$, $c = 0.298$, $T_{min} = 4.9$ °C, $T_{max} = 47.3$ °C. This relationship was used to plot $Z_{C,L}$ against $\lambda$ in Fig. S21. We truncated the plot at $T = 35$ °C because the data of (*23*) indicated that $\lambda$ begins decreasing around $T = 40$ °C. Source data, inferred $Z_{C,L}$ values and estimated growth rates are given in Supplementary Table 6.

##### 1.12 Source code and data availability.

Source code and most data required for analyses and figure generation is available at `https://github.com/flamholz/redox-proteome`. Files containing representative genomes and bac120 sequences were too large to host on GitHub but are available for download from GTDB directly at `https://gtdb.ecogenomic.org/downloads` (r207). Summary files derived from those are available in the GitHub repository.

#### 2 An introduction to the formal oxidation state of carbon, $Z_C$

Dry biomass is approximately 50% carbon (C) by mass and so much of central metabolism involves transformations of C atoms. Of course synthesis of biological macromolecules also requires hydrogen, nitrogren, oxygen, sulfur, phosphorus (collectively abbreviated CHNOPS), and many trace elements, especially metals. In this primer we focus on carbon.

In this section we explain how to calculate the nominal oxidation state of carbon ($Z_C$) for a molecule. We then show how this value can be used to calculate the number of reducing equival $e^-$ — i.e. electron carriers — required for a redox transformation. These calculations depend on assumptions, but when the assumptions hold they are exact.

Let's take the example of pyruvate as in Fig. S1. This key central metabolite is a glycolytic product and is a substrate for synthesis of several amino acids. $Z_C$ gives the average number valence $e^-$ 'associated' with C atoms in a molecule, where this value is calculated relative to a neutral C atom with 4 valence $e^-$. There are two ways to calculate $Z_C$ for pyruvate: (i) by counting valence electrons on a per-C basis or (ii) with a formula based on its elemental composition.

**Option (i): calculating $Z_C$ by enumerating C atoms**

For each C atom in pyruvate, we consider all of its bonds and ask if the $e^-$ pair is shared evenly across the bond (as in a C-C bond), primarily resides on the C atom (as in a C-H bond) or primarily resides on the other atom (as in a C-O or C-N bond). This is equivalent to asking which atom is more electronegative. If the $e^-$ pair "resides" on the C, then we add -1 to its ledger, if it resides on the other atom we add +1, and if it shared then we add 0. After summing these values across all C atoms (+2 for pyruvate) we divide by the number of C atoms $N_C$ (3 for pyruvate) to get the $Z_C$ (= +2/3 for pyruvate, Fig. S1).

**Balancing C and $e^-$ in reactions**

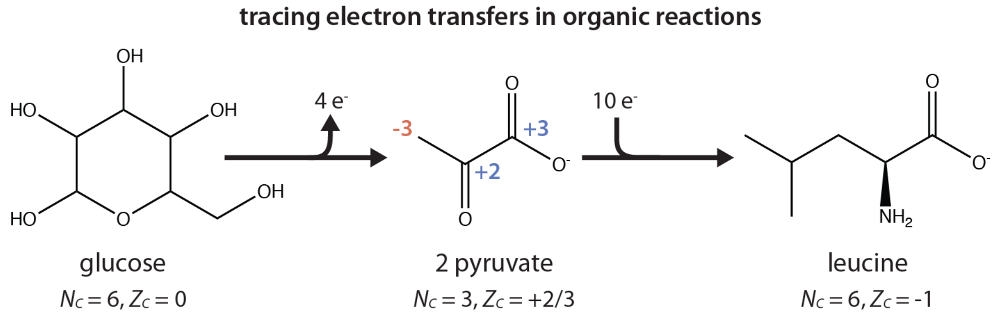

**Figure S1. Tracing electron transfers in organic reactions using the average redox state of carbon, $Z_C$.** For the central example of pyruvate (2-oxopropanoate) each carbon is marked with a formal charge. The terminal hydrocarbon has a formal charge of -3 due to 3 bonding hydrogens which are less electronegative than C. The central carbonyl group (C=O) has a formal charge of +2 since two electron pairs are shared with C atoms and two more reside on an O atom that is more electronegative than the focal C ($0+2=+2$). By the same logic the terminal carboxylate ($COO^-$) has a formal charge of +3. Summing these values ($-3+2+3=+2$) and dividing by the number of C atoms in pyruvate ($N_C = 3$) gives the nominal oxidation state of C atoms on pyruvate, $Z_C = +2/3$. Values for glucose (left) and leucine (right) can be calculated via the same procedure. Using $Z_C$ values we can now calculate the number of electrons produced in the oxidation of glucose → 2 pyruvate. This reaction already balances C atoms, so the change in formal charge is $6\times\frac{2}{3}-6\times 0 = 4$ e$^-$. A similar calculation for producing 1 leucine ($N_C = 6$) from 2 pyruvate gives $6\times -1-6\times\frac{2}{3} = -10$ e$^-$ indicating that this is a reductive process requiring input of 10 electrons per leucine.

If we perform the same operation for glucose we get $N_C = 6$ and $Z_C = 0$. Since pyruvate has $N_C = 3$ C atoms, we infer that glycolysis produces 2 pyruvate per glucose, meaning that there is a net excess of

$$2\cdot\text{pyruvate}\times\frac{2\ \text{C}}{\text{pyruvate}}\times\frac{+2}{3}\frac{\text{valence e}^-}{\text{pyruvate C}}-\frac{6\ \text{C}}{\text{glucose}}\times 0\frac{\text{valence e}^-}{\text{glucose C}} = 4\ \text{valence e}^- \quad (2)$$

in glucose relative to pyruvate. Since we consider 2 e$^-$ carriers here (e.g. NADH), we conclude that a reaction scheme producing 2 pyruvate from glucose (e.g. glycolysis) should produce 2 reduced carriers (NADH, generically termed ECH here) from 2 oxidized carriers ($NAD^+$ or $EC^+$) to balance e$^-$, as indeed is the case for all glycolytic pathways producing pyruvate.

**Assumptions in calculating $Z_C$ and balacing e$^-$**

Before exploring option (ii), it's useful to reflect on why this $Z_C$ calculation correctly predicts e$^-$ production/consumption in organic reaction systems. When we assign e$^-$ to the more electronegative bonding atom, what we are really doing is counting valence e$^-$ on C atoms relative to the neutral state of 4. If we break a C-C bond and we want the resulting C atoms to be neutrally charged (as they typically are in biological molecules), we need to add 2 e$^-$ to the system, one for each C. If we had broken a C-O bond, our scheme tells us that both of those e$^-$ would be added to the C atom rather than O (this is more detail than we need here).

Our $Z_C$-based electron-balancing calculation is premised on two assumptions. First we assume that C atoms are neutrally charged, i.e. that carbanions and carbocations are unstable in water, in accordance with biochemical intuition and measurements. Second, we assumed that the transformation of glucose to pyruvate is entirely determined by the C atoms in those molecules, i.e. that the 'metabolic story' of organic molecules is entirely told through the bonding patterns of C atoms. If, for example, we created and destroyed O-N or O-P bonds in this reaction scheme, tracking $Z_C$ would not be enough to track all the e$^-$: we would need to track $Z_N$ and $Z_P$ as well. Fortunately, these two assumptions are essentially correct when considering the core of metabolism.

**Option (ii): calculating $Z_C$ by formula**

The process of counting the formal charge of each carbon can be formalized with an equation. This formula applies to any organic molecule so long as it contains only C, H, N, O, P and S atoms.

$$Z_C = 4 - \frac{1}{N_C}\left(-q + 4N_C + N_H - 3N_N - 2N_O + 5N_P - 2N_S\right) \tag{3}$$

The above formula assumes that only C changes its oxidation state through metabolism, but allows for molecules with net charge $q$. See reference (*12*) for full explanation.

We can simplify equation (3) by noticing that $4 - \frac{4N_C}{N_C} = 0$ and that changes in charge $q$ that are due to protonation (i.e. adding 1 to $N_H$) have equal and opposite contribution to the equation. Therefore, for a molecule like an amino acid that carries no charge when fully protonated, we can simply consider the $q = 0$ protonation state in calculating $Z_C$.

$$Z_C(q=0) = \frac{3N_N - N_H + 2N_O - 5N_P + 2N_S}{N_C} \tag{4}$$

Finally, when considering specific classes of molecules, we can further simplify. For example, translated proteins[1], glucose, and pyruvate all lack phosphorus (P), yielding further simplification for proteins

$$Z_C(q=0) = \frac{3N_N - N_H + 2N_O + 2N_S}{N_C}. \tag{5}$$

Let's take the example of pyruvate, for which we previously manually calculated $Z_C = +2/3$. Pyruvate refers to the charged (deprotonated) form of pyruvic acid. The neutral ($q = 0$) acid has the formula $C_3H_4O_3$. If we apply equation 5 to pyruvic acid we find

$$Z_C(q=0) = \frac{-4 + 2\times 3}{3} = +\frac{2}{3}. \tag{6}$$

We can perform a similar calculation for the amino acid leucine, which has $Z_C = -1$ as shown in Fig. S1. The neutrally charged form of leucine has the atomic formula $C_6H_{13}NO_2$. Applying equation 5 gives

$$Z_C(q=0) = \frac{3\times 1 - 13 + 2\times 2}{6} = -1. \tag{7}$$

**Option (iib): calculating protein $Z_C$ from amino acids**

One useful lesson from equations (3): addition or removal of $H_2O$ has no effect on $Z_C$ since $N_H - 2N_O = 0$. This means that peptide bond formation/hydrolysis are $Z_C$ neutral, allowing us to calculate the $Z_C$ of a protein or proteome by taking the C-weighted average of constituent amino acids

$$Z_C = \frac{\sum_{\text{amino acids } i} N_{C,i} \cdot Z_{C,i}}{\sum_{\text{amino acids } i} N_{C,i}} = \frac{\text{protein C valence e}^-}{\text{total protein C atoms}} \tag{8}$$

as we do throughout this work. To see this equation in action, consider a dipeptide of leucine ($N_C = 6$, $Z_C = -1$) and glycine ($N_C = 3$, $Z_C = 1$). The carbon-weighted average of these values is

$$Z_C = \frac{6\times -1 + 3\times 1}{6+3} = -\frac{1}{3}. \tag{9}$$

[1] i.e., as opposed to post-translationally phosphorylated ones.

### 3 Derivation of the integrated redox + resource allocation model

In this section our goal is derive our integrated model from simple principles, which include conservation of mass and allocation of catalytic activity. In 3.1-3.2 we derive the relationship between the growth rate of a culture, $\lambda$, and the flows of matter (carbon, electrons) through metabolism. Then in 3.3 we define expressions for the fluxes through each intracellular metabolic process — oxidation, reduction, and anabolism — and connect these fluxes to both the resource allocation — the amount of each type of catalyst — and the growth rate. In the next section (**??**) we bring these derivations together to build a generic framework coupling any cellular metabolism (defined in redox terms) to a resource allocation framework.

#### 3.1 Definitions of growth rate $\lambda$

Since we are examining the relationship between growth, physiology and metabolic chemistry, we begin by defining the growth rate. This technical discussion may be repetitious for those familiar with Flux Balance Analysis (*24*) and related models (*25*), but is useful for readers new to the topic.

Several definitions of growth are used in the literature. Common definitions include growth in total cell number, total cell volume (collective volume of all cells), total cell mass (collective mass of all cells including water), total dry mass, and total carbon (C) mass. Notably, many standard methods of determining culture growth rates are direct or indirect measurements of the dry (water-free) mass of cell material (biomass). Optical densities, for example, are more consistently correlated with dry mass than with cell counts (*26*).

Considering the total mass ($M$) growth rate ($\lambda_M$) of an exponentially growing culture, we can see that $\lambda_M$ is the mass-specific time derivative of the total cell mass.

$$M(t) = M_0 \exp(\lambda_M t), \tag{10}$$

$$\frac{dM}{dt} = \lambda_M M_0 \exp(\lambda_M t), \tag{11}$$

$$\lambda_M = \frac{1}{M(t)} \frac{dM}{dt}. \tag{12}$$

If we assume that cells have constant density, i.e. $\rho = M/V$ is a constant (*27*), then we find that the volumetric and mass-specific growth rates are equal ($\lambda_V = \lambda_M$, (*24*)).

$$\begin{aligned} \frac{1}{M}\frac{dM}{dt} &= \frac{1}{M}\frac{dM}{dV}\frac{dV}{dt} \\ &= \frac{\rho}{M}\frac{dV}{dt} \\ &= \frac{1}{V}\frac{dV}{dt} = \lambda_V \end{aligned}$$

Likewise, if we assume a constant protein mass fraction $f_P = M_P/M$, as in (*28*), then $\lambda_P = \frac{f_P}{M_P}\frac{dM}{dt} = \lambda_M$. So when we make an appropriate compositional assumption — e.g. constant protein mass fraction — various definitions of $\lambda$ become equivalent.

Since we are tracing carbon atoms through metabolism here, it is convenient for us to focus on the carbon mass of the culture $M_C$. As such, we assume that the carbon mass fraction $f_C = M_C/M$ is constant, and so the growth rate of culture carbon mass $\lambda_C = \lambda_M$.

#### 3.2 Deriving the flux balance condition

Consider some carbon-bearing intracellular molecule with index $i$, e.g. pyruvate or ATP. We are interested in describing how its concentration, $C_i$, changes due to metabolic processes and growth. Metabolic processes can produce or consume $i$, but growth consumes $i$ by dilution through volumetric expansion or cell division.

The concentration $C_i$ is typically defined as number/volume, i.e., $N_i/V$ where $N_i$ is a number of moles. Yet we are interested in the flows of carbon atoms here, so we will now apply our compositional assumption ($f_C = M_C/M$ is constant) to transform number/volume concentrations into C mass/C mass concentrations.

Let $M_i$ be the total mass of C found as $i$ in the culture (gram units) and $m_i$ to be the mass of C in a mole of $i$ (g C/mol). It follows that that $N_i = M_i/m_i$ in units of moles. As such,

$$C_i \equiv \frac{N_i}{V} = \frac{M_i}{m_i V}. \tag{13}$$

Additionally, $\frac{1}{V} = \frac{\rho}{M} = \frac{f_C \cdot \rho}{M_C}$. As such, we see that

$$C_i = \frac{f_C \cdot \rho}{M_C} \frac{M_i}{m_i}. \tag{14}$$

We apply the product rule to to calculate the time derivative of $C_i$:

$$\frac{dC_i}{dt} = \frac{f_C \cdot \rho}{M_C} \frac{1}{m_i} \frac{dM_i}{dt} - \frac{f_C \cdot \rho}{M_C^2} \frac{M_i}{m_i} \frac{dM_C}{dt}. \tag{15}$$

The C specific growth rate $\lambda_C \equiv \frac{1}{M_C} \frac{dM_C}{dt}$ by definition. Combining this definition with equations (15-14), we find that

$$\frac{dC_i}{dt} = \left(\frac{dC_i}{dt}\right)_{M_C} - (C_i \lambda_C)_{M_i}. \tag{16}$$

That is, two terms contribute to the concentration dynamics of molecule $i$ – changes to $C_i$ at constant $M_C$ due to reactions in the cell, and changes to $C_i$ at constant $M_i$ due to changes in total (carbon) mass, i.e. due to dilution by growth.

Equation 14 tells us that

$$\frac{dC_i}{dt} = \frac{f_C \cdot \rho}{m_i} \left(\frac{d}{dt} \frac{M_i}{M_C}\right) \tag{17}$$

where $\frac{f_C \cdot \rho}{m_i}$ is assumed to be a constant. By factoring this constant term out of both sides of eq. (15) we find that

$$\frac{d}{dt}\left(\frac{M_i}{M_C}\right) = \left(\frac{1}{M_C}\frac{dM_i}{dt}\right)_{M_C} - \left(\frac{M_i}{M_C}\frac{1}{M_C}\frac{dM_C}{dt}\right)_{M_i}, \tag{18}$$

$$= \left(\frac{1}{M_C}\frac{dM_i}{dt}\right)_{M_C} - \left(\frac{M_i}{M_C}\lambda_C\right)_{M_i}, \tag{19}$$

Where we once again applied the definition of $\lambda_C \equiv \frac{1}{M_C}\frac{dM_C}{dt}$. For the remainder of this document, we will use $\lambda$ to refer to $\lambda_C$.

Equation 19 relates the change in a C mass fraction (a sort of concentration) to mass-specific reaction rates ($\frac{1}{M_C}\frac{dM_i}{dt}$) and dilution by growth ($\frac{M_i}{M_C}\lambda_C$). Note that while no factors of $\rho$ or $f_C$ appear, we had to assume these are constants independent of $M_i, M_C$ and $\lambda$ to factor them out.

By noting that (i) $M_i = N_i \cdot m_i$ and (ii) $m_i$ is a constant, we can write fluxes in terms of concentrations expressed as (number/$M_C$) fractions:

$$m_i \frac{d}{dt}\left(\frac{N_i}{M_C}\right) = m_i \left(\frac{1}{M_C}\frac{dN_i}{dt}\right)_{M_C} - m_i \left(\frac{N_i}{M_C}\lambda\right)_{N_i}. \tag{20}$$

This is convenient because biochemical reactions are commonly written with stoichiometries reflecting relative molecule numbers and not relative carbon masses. Hereafter we refer to the number/C mass concentration of molecule $i$ as $\chi_i$ so that the above becomes

$$\frac{d\chi_i}{dt} = \left(\frac{d\chi_i}{dt}\right)_{M_C} - (\chi_i \lambda)_{N_i}. \tag{21}$$

###### 3.2.1 Typical $\chi_i$ values

Typical metabolite concentrations range from 1 $\mu$M - 10 mM. Converting $C_i$ in mol/L units to mol/gC we get

$$C_i \frac{\text{mol}}{\text{L}} \times \frac{1}{1100} \frac{\text{L}}{\text{g cells}} \times \frac{1}{0.3} \frac{\text{g cells}}{\text{gDW}} \times 2 \frac{\text{gDW}}{\text{gC}} = 6 \times 10^{-3} C_i \frac{\text{mol}}{\text{gC}},$$

where $\rho = \frac{M}{V} \approx 1100$ g/L is the bouyant density of cells. So for a molecule $i$ with a 1 mM = $10^{-3}$ mol/L concentration, $\chi_i \approx 6 \times 10^{-6}$ mol/gC.

##### 3.3 Fluxes through metabolic processes

Above we derived a relationship (eq. 21) between the time derivative of an intracellular concentration $\chi_i$ (number/C mass) and the metabolic and growth processes affecting molecule $i$. In this section we define expressions for the fluxes through metabolic processes — e.g. oxidation or reduction — that affect concentrations of intracellular molecules (e.g. ATP) by producing or consuming them.

The first term in equation (21) describes to changes $\chi_i$ at constant $M_C$. These changes are due to various intracellular reactions that produce and consume $i$. We assign these reactions an index $\alpha$ and number flux $v_\alpha$, the latter having units of [mol/s]. Since process fluxes are defined in units of our choosing (e.g. C atoms/s) we must also define stoichiometric coefficients $S_{\alpha,i}$ translating process fluxes (units of our choosing) into fluxes of $\chi_i$. As such, the change in $\chi_i$ due to process $\alpha$ is

$$\underbrace{\frac{1}{M_C}}_{\chi_i \text{ unit conversion}} \times \underbrace{S_{\alpha,i}}_{\text{stoichiometry}} \times \underbrace{v_\alpha}_{\text{process flux}} \tag{22}$$

Since a single concentration can be affected by multiple metabolic processes $\alpha$, we sum these effects:

$$\frac{d\chi_i}{dt} = \left(\sum_\alpha \frac{1}{M_C} S_{\alpha,i} v_\alpha\right) - \chi_i \lambda. \tag{23}$$

To connect the flows of matter to the cellular allocation of catalytic resources, we must relate fluxes $v_\alpha$ to the composition of biomass. That is, cells must make enzymes to catalyze a process $\alpha$ (e.g. oxidation or anabolism). We now introduce variables $\phi_\alpha$ that describe the fraction of biomass carbon dedicated to catalysis of each process.

We treat $v_\alpha$ as if it is an irreversible reaction catalyzed by a single enzyme with Michaelis-Menten kinetics.

$$v_\alpha = \underbrace{\gamma_\alpha}_{\text{kinetics}} \times \underbrace{M^E_\alpha}_{\text{enzyme mass}} \times \underbrace{f(\vec{\chi})}_{\text{reactant conc. effect}} \tag{24}$$

Here $M^E_\alpha$ is the carbon mass of catalyst (enzyme) for process $\alpha$ and $\gamma_\alpha$ is the maximum rate of a mass unit of catalyst. $\gamma_\alpha$ has units of [mol $i$/gC catalyst/s]. The function $f_\alpha(\vec{\chi})$ describes the dependence of $v_\alpha$ on all metabolite concentrations $\vec{\chi}$. Since enzymes can consume multiple substrates, we use $f_\alpha(\vec{\chi})$ to indicate a unitless function of all concentrations. This will usually be a saturating function of a few $\chi_i$, e.g. $f_\alpha(\vec{\chi}) = \chi_i \cdot (K_M + \chi_i)^{-1}$ in the case of single substrate Michaelis-Menten rate law.

Converting this expression for $v_\alpha$ into units of $\chi_i$ flux following equation 22 yields

$$\frac{1}{M_C} S_{\alpha,i} v_\alpha = S_{\alpha,i} \gamma_\alpha \underbrace{(M^E_\alpha / M_C)}_{\phi_\alpha} f_\alpha(\vec{\chi}). \tag{25}$$

Here we defined $\phi_\alpha \equiv M^E_\alpha / M_C$ as the C mass fraction that is catalyst of process $\alpha$. The expression $S_{\alpha,i}\gamma_\alpha\phi_\alpha f(\vec{\chi})$ therefore has units of [mol $i$/gC/s], i.e., the same units as $\chi_i \lambda$.

##### 3.3.1 Approximate values of kinetic constants $\gamma_\alpha$

To estimate $\gamma_\alpha$ for a process catalyzed by a single enzyme we relate it to the enzyme $k_{cat}$ — the per-active-site maximum catalytic rate — by dimensional analysis. $\gamma_\alpha$ has units of [mol $i$/gC catalyst/s] while $k_{cat}$ has units of [mol product/mol enzyme/s]. As such,

$$\gamma_\alpha = \frac{1}{Y_{\alpha,i}} \frac{1}{m_\alpha} k_{cat}, \tag{26}$$

where $m_\alpha$ is carbon mass of a mole of catalyst and $Y_{\alpha,i}$ is the yield of product from substrate molecule $i$ in process $\alpha$ in units of [moles product/moles metabolite $i$]. For central metabolic reactions, $k_{cat} \approx 100\ \text{s}^{-1}$ ((*29*), Fig. S5). Enzymes have characteristic molar masses of a 50-100 kDa of which $\approx 50\%$ is C by mass (*30*). So $m_\alpha \approx 2-5 \times 10^4$ gC/mol. Further, typical $Y_{\alpha,i} \approx 1$. Therefore,

$$\gamma_{\alpha,i} \approx (2-5) \times 10^{-5} \times k_{cat} \tag{27}$$

in units of [mol product (g C)$^{-1}$ s$^{-1}$]. Since we assumed a generic value for $Y_{\alpha,i}$ we can omit the $i$ subscript from $\gamma_\alpha$.

In cases where we require a $\gamma_\alpha$ value for a metabolic pathway we assume that pathways are composed of $\approx 10$ enzymatic steps, each with equivalent kinetics. If we take $k_{cat} = 100\ \text{s}^{-1}$ (*29*), then we find $\gamma_\alpha \approx (2-5) \times 10^{-4}$ mol/gC/s.

##### 3.3.2 Calculating kinetic constants $\gamma_\alpha$ based on the ribosome

Another way to estimate $\gamma_\alpha$ values for pathways is to consider protein translation. Translation is catalyzed by the ribosome, a mega-complex of catalytic RNA with tens of small proteins having a $k_{cat} \approx 20\ \text{s}^{-1}$ and molar mass of $\approx 2.5 \times 10^6$ g/mol. Neglecting the masses of all other components of the translation cycle — e.g., initiation factors, tRNAs, and tRNA synthetases — then we get an upper bound of $\gamma_{tl} \lessapprox 10^{-5}$ mol amino acids per gram catalyst per second. As discussed below, our convention is to write reactions on a per C basis, e.g., transforming 1 C of amino acid into 1 C of protein. Since amino acids in typical proteins have $\approx 5$ C atoms, $\gamma_{tl} \lessapprox 5 \times 10^{-5}$ mol C per gram catalyst per second.

##### 3.3.3 Calculating of kinetic constants $\gamma_\alpha$ based on rubisco

For autotrophs using the Calvin-Benson cycle, Ribulose Bisphosphate Carboxylase/Oxygenase, or rubisco, is commonly considered to be a bottleneck that limits rates of growth (*31*). As such, many bioengineering efforts aim to improve rubisco itself or the physiology supporting its operation (*32*). Here we will estimate $\gamma_{rb}$, the value of $\gamma$ that we would use to describe rubisco, and compare to the ribosome.

Rubisco is the focal enzyme of the Calvin-Benson cycle, a cyclic metabolic pathway comprising $\approx 10$ enzymes. The Form I rubisco common to plants, algae and Cyanobacteria is a large protein complex with a mass of $\approx 500 \times 10^3$ [g/mol] and 8 active sites, or $\approx 60 \times 10^3$ [g/mol/site] (*33*). The rubisco carboxylation reaction adds a single $CO_2$to the five carbon organic substrate, ribulose 1,5-bisphosphate with a $k_{cat} \approx 1-10$ s$^{-1}$ depending on the isoform. Considering a substrate-saturate rubisco in isolation, therefore, gives an upper-bound of

$$\gamma_{rb} \lessapprox 10 \frac{\text{reactions}}{\text{s}} \times 1 \frac{\text{C}}{\text{reaction}} \times \frac{1}{60 \times 10^3} \frac{\text{mol active sites}}{\text{g rubisco}} \approx 2 \times 10^{-4} \frac{\text{mol C}}{\text{g rubisco} \times \text{s}}. \tag{28}$$

This value is a few times large than we estimated for the ribosome. Yet it is certainly too high for a number of reasons. First, we used the top-end of rubisco $k_{cat}$ values. Second, we neglected rubisco's promiscuous reactivity with $O_2$, which has the net effect of reducing the rate of carboxylation (*32*, *33*). Finally, we neglected all other enzymes in the cycle. These factors could, taken together, reduce our estimate by a factor of ten or more, yet the result value would still be similar to our estimate for the ribosome.

##### 3.3.4 Caveats associated with coarse-graining metabolic pathways

While the calculation treats pathways as if they are catalyzed by a single enzyme, biological processes like anabolism are catalyzed by multiple enzymes in sequential, cyclical or parallel pathways. So we are "coarse-graining" entire pathways as single enzymes. This is commonly done in resource allocation models (*25*, *28*, *34*), but has not yet been justified on empirical or theoretical bases to our knowledge. Justification could include (i) demonstrating consistency of this coarse graining with physiological measurements across conditions or (ii) deriving effective expressions for whole pathways from the kinetic properties of constituent enzymes. Noor and Liebermeister have made some recent progress on this second direction in (*35*).

#### 4 A generic redox resource allocation framework coupling metabolism and growth

In section 3 we derived equations relating changes in concentration to growth and metabolic processes in cells. We further connected those equations to expressions relating the quantity of intracellular enzymes to the flux through metabolic processes like anabolism. Here we collect these derivations into a generic model coupling metabolism, defined in redox terms, to growth. We will then describe how this generic model can be made to represent a specific metabolism, e.g. one of those depicted in Fig. S2.

We assume that each process $\alpha$ has a flux proportional to the resources allocated to it, i.e., the C mass fraction $\phi_\alpha$ defined in equation 25. The reaction representing each process also includes stoichiometric coefficients $S_{\alpha,i}$ for each participating metabolite $i$. Our convention here is to write $S_{\alpha,i}$ relative to carbon so that each reaction transforming organic molecules is written as transforming a single C atom, as in this example of glucose oxidation:

$$\frac{1}{6} \underbrace{\text{glucose}}_{C_6H_{12}O_6} + H_2O + 2\ \text{NAD+} \rightarrow\ CO_2 + 2\ \text{NADH}.$$

Writing atomically and electronically balanced reactions, as above, ensures that steady-state solutions of our model will conserve atoms and electrons both. The mass-flux through each process is given by

$$v_\alpha = \gamma_\alpha M_\alpha^E f(\vec{\chi}) \tag{29}$$

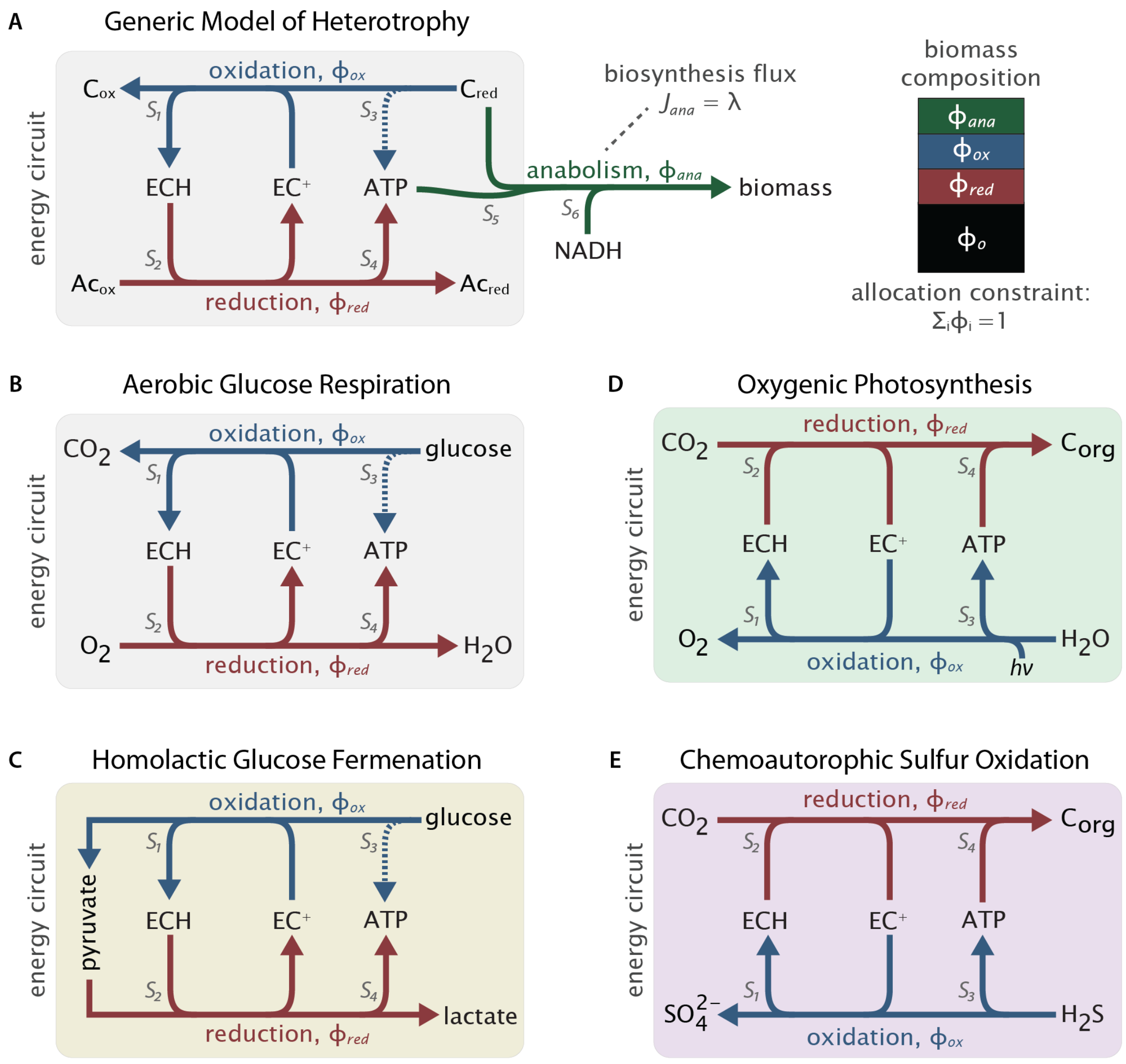

**Figure S2. Integrated redox + resource allocation models can describe any metabolism with equivalent equations.** Panel (A) shows a generic model or respiratory heterotrophy, oxidizing $C_{red}$ to $C_{ox}$ while reducing a terminal $e^-$ acceptor $Ac_{ox}$ to $AC_{red}$. Because acceptor reduction is coupled to ATP synthesis, this "energy circuit" has the net effect of coupling $e^-$ flow to synthesis of ATP from ADP and inorganic phosphate. In all of our models we assume that $e^-$ are carried between processes via a generic two-electron redox couple ECH/$EC^+$ denoting "electron carrier." Anabolism typically consumes ATP and, depending on the redox state of biomass carbon ($Z_{C,B}$) might also consume electron carriers. All of these cellular processes are catalyzed by enzymes, so the flux $J_\alpha$ through each process is proportional to $\phi_\alpha$, the fraction of biomass carbon that is catalyst for process $\alpha$. So, for example, $\phi_{ana}$ denotes the fraction that catalyzes anabolism. Panel (B) gives a concrete example of glucose respiration, (C) shows a fermentation of glucose, (D) photosynthetic $CO_2$fixation, and (E) chemoautotrophic sulfur oxidation. Anabolism and the allocation constraint are omitted from panels (B-E) for space.

and the concentration dynamics for metabolite $i$ are

$$\frac{d\chi_i}{dt} = \left(\sum_\alpha \frac{1}{M_C} S_{\alpha,i} \nu_\alpha\right) - \chi_i \lambda \tag{30}$$

$$= \left(\sum_\alpha S_{\alpha,i} \gamma_\alpha \phi_\alpha f(\vec{\chi})\right) - \chi_i \lambda \tag{31}$$

$$= \left(\sum_\alpha S_{\alpha,i} J_\alpha\right) - \chi_i \lambda, \tag{32}$$

where we've defined mass-specific mass fluxes

$$J_\alpha \equiv \frac{\nu_\alpha}{M_C} = \gamma_\alpha \phi_\alpha f(\vec{\chi}) \tag{33}$$

in units of [mol product $gC^{-1}$ $s^{-1}$]. If we assume that catalysts are substrate-saturated then $f(\vec{\chi}) = 1$. If we instead assume first-order reaction kinetics (the linear low-concentration regime of a Michaelis-Menten rate law), then

$$f(\vec{\chi}) = \prod_{i=1}^{N_{sub}} \left(\frac{C_i}{K_{M,i}}\right)^{Q_{\alpha,i}}. \tag{34}$$

Where $K_{M,i}$ is the half-saturating concentration of metabolite $i$ for process $\alpha$. Note that the exponents $Q_{\alpha,i} \neq S_{\alpha,i}$ in general. $S_{\alpha,i}$ are stoichiometric coefficients used for mass-balancing and have arbitrary scale of our choosing. $Q_{\alpha,i}$ values are called 'molecularities.' These values are related to the reaction mechanisms of enzymes catalyzing specific pathway steps where a certain number of substrates $i$ and $j$ come together in enzyme active sites. For typical enzymatic reactions, $Q_{\alpha,i} = 1$, as we assume below.

Notice also that the product runs over $N_{sub}$, the number of substrates in the reaction. This reflects our assumption metabolic processes $\alpha$ are irreversible. In some low energy environments this assumption may be invalid; we hope future work explores reversible versions of such models.

In addition to the cellular processes $\alpha$, we admit that some fraction of biomass C is non-catalytic, e.g. lipids and storage molecules like glycogen. We term this C mass fraction $\phi_O$ and enforce the carbon mass allocation constraint

$$\left(\sum_\alpha \phi_\alpha\right) + \phi_O = 1. \tag{35}$$

While we have written process fluxes $J_\alpha$ in a generic form above, individual processes are distinguished by their specific chemistry and role in any specific metabolism as we discuss below and diagram in Fig. S2B-E.

Two redox reactions, termed 'oxidation' and 'reduction' here, together form an 'energy circuit' whose net effect is to generate a flow of ATP by extracting electrons from a donor and conveying them to a terminal acceptor by way of an electron carrier (Fig. S2A). Throughout we model a single 2 $e^-$ carrier that we call ECH/$EC^+$ for "electron carrier." ECH denotes the reduced ($e^-$ carrying) form and $EC^+$ the oxidized ($e^-$ poor) form. Since cells use more than one carrier (*36*) modeling a single carrier is a simplification. A third redox reaction, anabolism, represents the synthesis of biomass C from nutrients, ATP and ECH. Fluxes $J_{ox}$, $J_{red}$, $J_{ana}$ are proportional to C mass fractions $\phi_{ox}$, $\phi_{red}$, and $\phi_{ana}$. Given these definitions, the allocation constraint is

$$\phi_{ox} + \phi_{red} + \phi_{ana} + \phi_O = 1. \tag{36}$$

#### 4.1 Converting $J_{ana}$ to $\lambda$

Having defined the relationships between metabolic fluxes, concentrations and dilution, we now turn to the growth rate, $\lambda$. Anabolism produces new biomass carbon with flux $J_{ana}$. We defined growth as the production of new carbon mass with $\lambda \equiv \frac{1}{M_C}\frac{dM_C}{dt}$. Yet $J_{ana} = \nu_{ana}/M_C$ is a mass-specific number flux in units of [mol C / gC / s] and $\lambda$ has [s$^{-1}$] units. The conversion factor is exactly the molar mass of carbon:

$$\lambda = m_C \cdot J_{ana}, \tag{37}$$

where $m_C = 12$ gC/mol C, giving correct units of [gC/gC/s] for $\lambda$. If we assume a value for $\gamma_{ana}$ we can use this relationship to estimate the growth rate from the anabolic flux:

$$\lambda = m_C \cdot \gamma_{ana} \cdot \phi_{ana} \cdot f(\vec{\chi}).$$

Above we estimated $\gamma_{tl} \lessapprox 5 \times 10^{-5}$ mol C/g C/s for translation. If we use this value for $\gamma_{ana}$, assume $f(\vec{\chi}) = 1$ and note that $\phi_{ana} \approx 0.1$ we estimate[2]

$$\begin{aligned}\lambda &\approx 12 \frac{\text{g}}{\text{mol C}} \times 5 \times 10^{-5} \frac{\text{mol C}}{\text{g enz c}} \times 0.1 \frac{\text{g enz C}}{\text{g tot C}} \\ &\approx 6 \times 10^{-5}\ \text{s}^{-1} \\ &\approx 0.2\ \text{hr}^{-1}.\end{aligned}$$

Though the fastest measured heterotrophic growth rates are a few generations per hour ($\lambda = 2-4$ hr$^{-1}$, Fig. 3C), typical growth rates of laboratory cultures in minimal media are indeed $0.1-1$ hr$^{-1}$ (*16*).

#### 4.2 Maintenance energy

The minimum energy expenditure of living matter is usually termed $m$ and denominated in energetic units. Due to our use of $m$ and $M$ for masses, we will use $b$ instead of $m$ for 'basal.' Values of $b$ are typically reported in millimoles of ATP per gram dry weight per hour ([mmol ATP/gDW/hr]) and give the minimum ATP flux required to sustain living matter. A minimum maintenance energy is enforced by subtracting $b$ from the ATP mass balance:

$$\frac{d\chi_{\text{ATP}}}{dt} = \left(\sum_{\alpha=1}^{N_{proc}} S_{\alpha,i} J_\alpha\right) - \lambda\chi_{\text{ATP}} - b. \tag{38}$$

Empirical values of $b \approx 10$ mmol ATP/gDW/hr, which are easily converted to $b \approx 5 \times 10^{-6}$ mol ATP/gC/s by assuming C makes up $\approx 1/2$ of the total dry mass. Note that $b$ accounts for non-growth associated maintenance. Growth-associated maintenance — ATP costs proportional to $\lambda$ — can be accounted for by increasing the ATP requirements of anabolism ($S_5$ in Fig. S2).

##### 4.2.1 Comments on the maintenance energy

Before moving on, two comments on the maintenance energy. First, mounting evidence over decades indicates that maintenance energies estimated from laboratory cultures are several orders too large to account for microbial populations in the wild, e.g. in deep sediments (*37*). This may be due to the method of measurement, which usually involves projecting metabolic rates to zero growth from relatively fast growth rates on order 0.1 hr$^{-1}$ (*38*, *39*). The discrepancy between lab measurements and inferences from environmental constraints strongly suggests that microbes in the wild access low energy physiological states that are qualitatively unlike lab cultures.

[2] $\phi_{ana}$ cannot realistically exceed 0.6 if protein + RNA make up no more than 60% biomass (*4*) as shown in Fig. 5B.

Secondly, the standard treatment of maintenance energy assumes ATP units (*38*, *39*). However, some maintenance, e.g. repair of oxidized proteins or metabolites, might be properly denominated in redox (NADH) units (*40*). These units are not directly interconvertible, as is sometimes assumed (*41*), since converting NADH into ATP requires catalyst, i.e. requires increasing some $\phi_\alpha$. Future work should consider the implications of denominating maintenance in both energy and reductant units.

#### 4.3 Cycling co-factors like ATP and NADH

Since our models are simplified they do not explicitly describe the biosynthesis of co-factors like ATP and ECH (the redox co-factor in our model). Since oxidation of 1 ECH always produces 1 $EC^+$, these molecules have equal and opposite reaction fluxes

$$0 = \left( \sum_{\alpha=1}^{N_{proc}} S_{\alpha,\mathrm{EC}^+} J_\alpha \right) - \left( \sum_{\alpha=1}^{N_{proc}} S_{\alpha,\mathrm{ECH}} J_\alpha \right).$$

Note that this would not be true if we explicitly represented ECH biosynthesis, as the pathway would produce one of the two forms (say $EC^+$) and so the fluxes would not be equal and opposite. Rather, their difference would equal the flux of $EC^+$ biosynthesis.

If we enforce steady-state mass balance ($\frac{d\mathrm{ECH}}{dt} = \frac{d\mathrm{EC}^+}{dt} = 0$) in the absence of explicit biosynthesis, we find that

$$\lambda \chi_{\mathrm{NAD}^+} = \lambda \chi_{\mathrm{NADH}}.$$

It follows that reduced and oxidized forms have equal concentrations. Yet there is substantial evidence to the contrary, with $NAD^+$/NADH $\approx 10$ being fairly typical (*30*, *42*, *43*). A similar problem arises for ATP and ADP. To address this challenge one could either (i) explicitly encode biosynthesis into the model, as in genome-scale flux balance models (*24*, *44*), (ii) enforce mass balance for only one of the two cycling forms, e.g., only for ATP and NADH. We pursued the latter approach because it is simpler.

| coefficient | process | description | typical sign in het. | in autotrophy |
|---|---|---|---|---|
| $S_1$ | oxidation | per-C ECH produced | + | + |
| $S_2$ | reduction | per-C ECH produced | - | - |
| $S_3$ | oxidation | per-C ATP produced | + | + |
| $S_4$ | reduction | per-C ATP produced | + | - |
| $S_5$ | anabolism | per-C ATP produced | +/- | +/- |
| $S_6$ | anabolism | per-C ECH produced | +/- | +/- |

**Table 2.** Description of stoichiometric coefficients used in our metabolic models. By convention, we absorb signs in $S_i$ so that the mass balance can be written as a sum, as in eq. (31). Typical signs for heterotrophic and autotrophic metabolisms are given. Note that certain coefficients can be positive or negative in realistic settings, e.g., $S_3, S_5$, and $S_6$. In heterotrophy, $S_3 < 0$ arises if carbon source oxidation requires ATP investment. $S_5 < 0$ connotes ATP-producing anabolism, which arises when making biomass from energy-rich carbon sources, and $S_6 < 0$ occurs when the C source is more oxidized than biomass.

#### 4.4 Maximizing the growth rate with linear programming

The above integrated redox + resource allocation model induces an optimization problem to maximize the exponential growth rate

$$\begin{aligned}
\max \quad & \lambda = m_C J_{ana} \\
\text{s.t.} \quad & 0 = \left(\sum_\alpha S_{\alpha,\text{ATP}} J_\alpha\right) - \chi_{\text{ATP}}\lambda - b \text{ for ATP} \\
& 0 = \left(\sum_\alpha S_{\alpha,i} J_\alpha\right) - \chi_i \lambda \text{ for all other balanced mets. } i \\
& 0 = \left(\sum_\alpha \phi_\alpha\right) + \phi_O - 1 \\
\text{where} \quad & J_\alpha = \gamma_{\alpha,i}\phi_\alpha f(\vec{\chi})
\end{aligned}$$

Since dilution terms multiply two variables ($\lambda\chi_i$) and rate laws might also multiply variables (e.g. if $f(\vec{\chi})$ is non-trivial) this is a non-linear optimization problem. We can linearize by holding concentrations $\chi_i$ constant, permitting maximization of the growth rate at fixed $\chi_i$ as a function of $\phi_\alpha$. This is akin to a coarse-grained Resource Balance Analysis (*45*) or Constrained Allocation Flux Balance Analysis (*46*). To corroborate intuition drawn from analytics and simulations, we sweep wide ranges of values for each $\chi_i$ (several orders-of-magnitude). This is tractable because our models consider only 2-3 internal metabolites (e.g., ATP, NADH), and these are the only ones for which mass-balancing constraints apply.

### 5 A model of heterotrophic respiration

In this section we will describe the model of heterotrophic respiration used in the main text. This model is a specific instantiation of our generic model described in 4.

Our models include three processes: oxidation (mass-specific mass flux $J_{ox}$, associated mass-fraction $\phi_{ox}$), reduction ($J_{red}$, $\phi_{red}$) and anabolism ($J_{ana}$, $\phi_{ana}$), represented with per C atom stoichiometric coefficients $S_1,\ldots,S_6$ as labeled in Fig. S2. In heterotrophic respiration, the carbon source $C_{red}$ is oxidized to $C_{ox}$ (e.g. $CO_2$) to reduce the terminal acceptor (e.g. $O_2$) and generate ATP.

Let $A = \chi_{\text{ATP}}$ and $N = \chi_{\text{ECH}}$ for brevity. Mass-balances for ATP and ECH are then given by

$$\frac{dN}{dt} = (S_1 J_{ox} + S_2 J_{red} + S_6 J_{ana}) - \lambda N, \tag{39}$$

$$\frac{dA}{dt} = (S_3 J_{ox} + S_4 J_{red} + S_5 J_{ana}) - \lambda A - b, \tag{40}$$

where $S_i$ may be positive or negative depending on the specific metabolism, as described in Table (2). Our choice to absorb the signs in $S_i$ means that identical equations apply for autotrophs, as we discuss below.

At steady-state the above derivatives equal 0. By noting $\lambda = m_C J_{ana}$ we arrive at two different expressions for $\lambda$ which can be combined to simplify by eliminating either $J_{ox}$ or $J_{red}$. If we eliminate $J_{red}$ then

$$\lambda = -\frac{m_C\left(S_2\left(b - S_3 J_{\text{ox}}\right) + S_1 S_4 J_{\text{ox}}\right)}{m_C\left(AS_2 - NS_4\right) - S_2 S_5 + S_4 S_6} \tag{41}$$

The allocation constraint is as written in eq. (36) and carbon mass-specific process rates are be written as follows

$$J_{ox} = \gamma_{ox}\phi_{ox}f_{ox}(\vec{\chi}), \tag{42}$$
$$J_{red} = \gamma_{red}\phi_{red}f_{red}(\vec{\chi}), \tag{43}$$
$$J_{ana} = \gamma_{ana}\phi_{ana}f_{ana}(\vec{\chi}). \tag{44}$$

#### 5.1 An comparable model of photosynthesis

A great advantage of redox-based models is that all cells perform oxidation, reduction and anabolism. As such, the same framework that represents respiration can also describe photosynthesis with nearly the same equations. Indeed, oxygenic photosynthesis is roughly the opposite metabolism of respiration. While in oxygen respiration organic matter is oxidized (e.g. to $CO_2$) to reduce $O_2$ to $H_2O$, photosynthetic organisms oxidizes $H_2O$ to $O_2$ and use the extracted $e^-$ to reduce $CO_2$ to organic molecules (Fig. S2). So while processes with all the same names are present in photosynthesis, they represent very different underlying biochemistry.

Given that oxidation, reduction, and anabolism take place in photosynthesis, we can use the equations in 5 to describe photosynthesis or chemoautotrophy as well. Two modifications are required. First, reasonable values of stoichiometric coefficients differ between photosynthesis and respiration. In particular, the ATP yield of reduction is a positive number in respiration (reduction of the terminal acceptor produces ATP). In contrast, the reductive process in photosynthesis is $CO_2$ fixation which requires input of ATP. As such, $S_4$ is negative in models of autotrophic growth (Table 2).

Secondly, autotrophs produce and use reduced organic carbon (termed $C_{red}$ here) intracellularly while heterotrophs draw $C_{red}$ from the extracellular environment. In other words, $C_{red}$ is an intracellular metabolite in autotrophs and we must track its concentration dynamics. If $\chi_C$ is the number/C mass concentration of $C_{red}$, then

$$\frac{d\chi_C}{dt} = J_{red} - J_{ana} - \lambda\chi_C. \tag{45}$$

We can omit stoichiometric coefficients from this expression because we write reduction and anabolism fluxes per C atom. Demanding intracellular mass conservation as in 4.4 therefore requires that fluxes producing and consuming $C_{red}$ balance, i.e. that $\frac{d\chi_C}{dt} = 0$. This constraint on steady-state autotrophic growth is absent from models of heterotrophy and, as discussed in the main text and described in Fig. S13, always reduces autotrophic growth rates.

#### 5.2 Solutions when fluxes $J_\alpha$ are concentration independent

For the remainder of this section we will return to the model of heterotrophic respiration described in section 5. If we assume a functional form for $f_\alpha(\vec{\chi})$, we can derive expressions for $\lambda$ that include biomass fractions $\phi_\alpha$. For simplicity, if $f_\alpha(\vec{\chi}) = 1$ then we find that

$$\lambda = \frac{\gamma_{\text{ana}} m_C \left(S_1\gamma_{\text{ox}}\left(b + S_4\left(\phi_O - 1\right)\gamma_{\text{red}}\right) - S_2\gamma_{\text{red}}\left(b + S_3\left(\phi_O - 1\right)\gamma_{\text{ox}}\right)\right)}{\gamma_{\text{red}}\left(\gamma_{\text{ana}}\left(m_C\left(AS_2 - NS_4\right) - S_2S_5 + S_4S_6\right) + \left(S_2S_3 - S_1S_4\right)\gamma_{\text{ox}}\right) + \gamma_{\text{ana}}\gamma_{\text{ox}}\left(m_C\left(NS_3 - AS_1\right) + S_1S_5 - S_3S_6\right)} \tag{46}$$

If we assume that $\gamma_{ox} = \gamma_{red} = \gamma_{ana} = \gamma$ then this simplifies to

$$\lambda = \frac{m_C\left(S_2\left(b + \gamma S_3\left(\phi_O - 1\right)\right) - S_1\left(b + \gamma S_4\left(\phi_O - 1\right)\right)\right)}{m_C\left(AS_1 - AS_2 - NS_3 + NS_4\right) + S_1\left(S_4 - S_5\right) + S_2\left(S_5 - S_3\right) + \left(S_3 - S_4\right)S_6}.$$

Notice that we must treat ATP and NADH concentrations as constants here because they appear only in dilution terms in the zero-order case where $J_\alpha$ are concentration-independent.

One deficiency of this simple model is that it fails to grow when $S_6$ is too extreme. As discussed in the main text, some $S_i$ are constants given a choice of metabolic substrates and products (NADH stoichiometries $S_1, S_2$) and others are subject to thermodynamic constraints (ATP stoichiometries $S_3, S_4, S_5$). Here we vary $S_6$ because its value is determined by the $Z_C$ difference between the organic C source and biomass, and is therefore determined in part by the regulated and $\lambda$-dependent composition of biomass. By calculating the values of $S_6$ at which $\phi_{ox}$ and $\phi_{red}$ cross 0 we can calculate upper- and lower-bounds on feasible $S_6$.

$$S_6 \geq \frac{AbS_2\gamma_{\text{ana}}m_C - bS_2S_5\gamma_{\text{ana}} + bS_2S_4\gamma_{\text{red}}}{S_4\gamma_{\text{ana}}\left(b + S_4\phi_O\gamma_{\text{red}} - S_4\gamma_{\text{red}}\right)} + \frac{-AS_2m_C + NS_4m_C + S_5S_2}{S_4} \tag{47}$$

$$S_6 \leq \frac{AbS_1\gamma_{\text{ana}}m_C - bS_1S_5\gamma_{\text{ana}} + bS_1S_3\gamma_{\text{ox}}}{S_3\gamma_{\text{ana}}\left(b + S_3\phi_O\gamma_{\text{ox}} - S_3\gamma_{\text{ox}}\right)} + \frac{-AS_1m_C + NS_3m_C + S_5S_1}{S_3} \tag{48}$$

These bounds depend on $\phi_O$ and concentrations $A, N$. While $0 \leq \phi_O \leq 1$ is not a constant, physiological considerations puts limits on its value. Protein and RNA are, to a large degree, to only catalytic components of biomass. In *E. coli*, these together make up $\leq 60\%$ of dry mass (*4*). So we can reasonably conclude that $\phi_O \geq 0.4$. These biological constraints on $\phi_O$ affect the values of $S_6$ compatible with growth.

Electron carrier stoichiometries ($S_1$ and $S_6$) are simple functions of the redox states of the C source ($Z_{C,red}$) biomass ($Z_{C,B}$) and the metabolic product ($Z_{C,ox}$). For a heterotroph, these relationships are

$$S_1 = \frac{1}{2}\left(Z_{C,ox} - Z_{C,red}\right), \tag{49}$$

$$S_6 = \frac{1}{2}\left(Z_{C,B} - Z_{C,red}\right). \tag{50}$$

As such, bounds on $S_6$ can be rewritten as bounds on $Z_{C,red}$. Here we reproduce bounds for the case that maintenance $b = 0$.

$$Z_{C,red} \geq Z_{C,B} - \frac{2\left(-AS_2m_C + NS_4m_C + S_5S_2\right)}{S_4}, \tag{51}$$

$$Z_{C,red} \leq \frac{Z_{C,ox}\left(Am_C - S_5\right)}{Am_C + S_3 - S_5} - \frac{S_3\left(2Nm_C - Z_{C,B}\right)}{Am_C + S_3 - S_5} \tag{52}$$

Note that these only apply in the absence of a homeostatic process balancing ATP and ECH production fluxes with their anabolic consumption. In the presence of such a process (e.g. regulated ATP hydrolysis, flux $J_H = \gamma_H\phi_H f_H(\vec{\chi})$) growth becomes feasible across the full range of $S_6$ or $Z_{C,red}$ values as we discuss in main text. Full expressions are given in the relevant Mathematica notebook.

If we assume constant ATP, NADH, $\phi_O$ and $\phi_H$, we can use Lagrange multipliers to maximize $\lambda$. The Lagrangian for the zero-order case is

$$\mathscr{L} = \lambda - \beta_1 g - \beta_2 h - \beta_3 i, \tag{53}$$

$$g = \phi_{ox} + \phi_{red} + \frac{\lambda}{m_C\gamma_{ana}} + \phi_H + \phi_O - 1, \tag{54}$$

$$h = S_1\gamma_{ox}\phi_{ox} + S_2\gamma_{red}\phi_{red} + S_6\frac{\lambda}{m_C} - \lambda\cdot N, \tag{55}$$

$$i = S_3\gamma_{ox}\phi_{ox} + S_4\gamma_{red}\phi_{red} + S_5\frac{\lambda}{m_C} - \gamma_H\phi_H - \lambda\cdot A - b \tag{56}$$

where $\beta_i$ are Lagrange multipliers and we've used $\lambda = m_C J_{ana} = m_C\gamma_{ana}\phi_{ana}$ to eliminate $\phi_{ana}$ from $g, h,$ and $i$. Solving $\nabla\mathscr{L}(\lambda, \phi_{ox}, \phi_{red}, \beta_1, \beta_2, \beta_3) = 0$ gives

$$\lambda_{max} = -\frac{(S_2\gamma_{\text{red}} - S_1\gamma_{\text{ox}})(-b - \gamma_H\phi_H + S_3\gamma_{\text{ox}}(-\phi_H - \phi_O + 1)) - S_1\gamma_{\text{ox}}(-\phi_H - \phi_O + 1)(S_4\gamma_{\text{red}} - S_3\gamma_{\text{ox}})}{(S_2\gamma_{\text{red}} - S_1\gamma_{\text{ox}})\left(-A - \frac{S_3\gamma_{\text{ox}}}{\gamma_{\text{ana}}m_C} + \frac{S_5}{m_C}\right) - (S_4\gamma_{\text{red}} - S_3\gamma_{\text{ox}})\left(-\frac{S_1\gamma_{\text{ox}}}{\gamma_{\text{ana}}m_C} + \frac{S_6}{m_C} - N\right)} \tag{57}$$

If ATP and NADH concentrations are roughly constant, e.g., due to thermodynamic and kinetic constraints (*36*, *42*, *43*, *47*), then $\lambda_{max}$ is a function of $\phi_O$ and $\phi_H$ alone. Therefore, $\phi_O$, $\phi_H$, or both must be variable if cells are to grow at a variety of rates in response to environmental conditions, as we concluded in the main text.

We verified that this expression for $\lambda_{max}$ corresponds exactly to linear programming solutions to the corresponding zero-order system with dilution. Furthermore, the analytic limits on $S_6$ and $Z_{C,red}$ correspond exactly to the points at which a zero-order model without flux-balancing ATP homeostasis fails to grow. Simulations of higher-order models give comparable results for $\lambda_{max}$ and also exhibit finite limits on growth as a function of $Z_{C,red}$ as we discuss below.

##### 5.3 Optimizing the redox state of biomass for growth

Using Lagrange multipliers we were also able to solve for the growth rate maximizing value of $Z_{C,B}$. The optimum value $Z^*_{C,B}$ can be expressed as a linear function of $Z_{C,red}$

$$Z^*_{C,B} = K_Z \cdot Z_{C,red} + Z^\circ \tag{58}$$

where the slope $K_Z$ and intercept $Z^\circ$ are functions of all model parameters and, therefore, related to intracellular fluxes $J_\alpha$. For a zero-order model of heterotrophy,

$$\begin{aligned}
K_Z = {} & \frac{Am_C\gamma_{\text{ox}}}{S_3\gamma_{\text{ox}} - S_4\gamma_{\text{red}}} + \frac{S_4\gamma_{\text{red}}(\gamma_{\text{ana}} - \gamma_{\text{ox}})}{\gamma_{\text{ana}}(S_4\gamma_{\text{red}} - S_3\gamma_{\text{ox}})} + \frac{bm_C\gamma_{\text{ox}}}{\lambda_{max}(S_3\gamma_{\text{ox}} - S_4\gamma_{\text{red}})} + \frac{m_C\gamma_H\phi_H\gamma_{\text{ox}}}{\lambda_{max}(S_3\gamma_{\text{ox}} - S_4\gamma_{\text{red}})} - \\
& \frac{S_4m_C\phi_H\gamma_{\text{ox}}\gamma_{\text{red}}}{\lambda_{max}(S_4\gamma_{\text{red}} - S_3\gamma_{\text{ox}})} + \frac{S_4m_C\phi_O\gamma_{\text{ox}}\gamma_{\text{red}}}{\lambda_{max}(S_3\gamma_{\text{ox}} - S_4\gamma_{\text{red}})} + \frac{S_4m_C\gamma_{\text{ox}}\gamma_{\text{red}}}{\lambda_{max}(S_4\gamma_{\text{red}} - S_3\gamma_{\text{ox}})} + \frac{(S_3 - S_5)\gamma_{\text{ox}}}{S_3\gamma_{\text{ox}} - S_4\gamma_{\text{red}}}. \\
Z^\circ = {} & (\lambda_{max}\gamma_{\text{ana}}(S_3\gamma_{\text{ox}} - S_4\gamma_{\text{red}}))^{-1} \times (\gamma_{\text{ox}}Z_{C,\text{ox}}(-A\lambda_{max}\gamma_{\text{ana}}m_C - b\gamma_{\text{ana}}m_C - \gamma_{\text{ana}}m_C\gamma_H\phi_H - S_4\gamma_{\text{ana}}m_C\phi_H\gamma_{\text{red}} - \\
& S_4\gamma_{\text{ana}}m_C\phi_O\gamma_{\text{red}} + S_4\gamma_{\text{ana}}m_C\gamma_{\text{red}} + \lambda_{max}S_5\gamma_{\text{ana}} - \lambda_{max}S_4\gamma_{\text{red}})) + (2(A\lambda_{max}S_2\gamma_{\text{ana}}m_C\gamma_{\text{red}} + bS_2\gamma_{\text{ana}}m_C\gamma_{\text{red}} + \\
& S_2S_3\gamma_{\text{ana}}m_C\phi_H\gamma_{\text{ox}}\gamma_{\text{red}} + S_2\gamma_{\text{ana}}m_C\gamma_H\phi_H\gamma_{\text{red}} + \lambda_{max}NS_3\gamma_{\text{ana}}m_C\gamma_{\text{ox}} - \lambda_{max}NS_4\gamma_{\text{ana}}m_C\gamma_{\text{red}} + \\
& S_2S_3\gamma_{\text{ana}}m_C\phi_O\gamma_{\text{ox}}\gamma_{\text{red}} - S_2S_3\gamma_{\text{ana}}m_C\gamma_{\text{ox}}\gamma_{\text{red}} - \lambda_{max}S_2S_5\gamma_{\text{ana}}\gamma_{\text{red}} + \lambda_{max}S_2S_3\gamma_{\text{ox}}\gamma_{\text{red}})),
\end{aligned} \tag{59}$$

where $\lambda_{max}$ is given by eq. 57.

Since adapting $Z_{C,B}$ to match the right hand side of equation 58 increases $\lambda$, we consider $Z^*_{C,B}$ to be an effective environmental redox potential. In contrast with typical approaches to characterizing environmental redox state (*21*), which account only for those molecules that react with the electrode, $Z^*_{C,B}$ accounts for the carbon source $Z_{C,red}$, the metabolic transformations performed (i.e. producing $Z_{C,B}$ and $Z_{C,ox}$) as well as the magnitudes of intracellular fluxes.

#### 6 Models with concentration-dependent fluxes

Thus far, we have restricted ourselves to the simplifying assumption that the fluxes $J_\alpha$ are independent of the concentrations of ATP and the electron carrier ECH (here assumed to be NADH). In this saturated regime, $f(\vec{\chi})$ was always set to 1. This regime allowed us to compute the guaranteed global maximum growth rate $\lambda_{max}$ in different conditions using linear programming and analytics, but restricted us to manually fix the steady state ATP and NADH concentrations. To test the robustness of our results to relaxing this assumption, we used simulations and numerical nonconvex optimization methods, which we detail in this section. These methods allowed the ATP and NADH concentrations to emerge from the dynamics specified by our model.

#### 6.1 Simulations

We performed simulations in two regimes: (1) a zero order model, where fluxes were independent of the ATP and NADH concentrations, but numerically reached their steady state values, and (2) a Michaelis-Menten model, where fluxes depended on the ATP and NADH concentrations according to Michaelis-Menten kinetics.

To simulate both models, we numerically evolved the system of differential equations described by equations 15 (parameters given in Table 2) with their mass fluxes $J_\alpha$ set as per different assumptions for $f(\vec{\chi})$. In the zero order model (Fig. S7a), we assumed $f(\vec{\chi}) = 1$ for all processes. In the Michaelis-Menten model (Fig. S7b), we assumed the following: (a) oxidation was independent of the ATP and NADH concentrations; (b) reduction depended only on the NADH concentration, $f_{red}(NADH) = \frac{NADH}{NADH+K_M}$; (c) anabolism depended on both the ATP and NADH concentrations, $f_{ana}(NADH) = \frac{NADH}{NADH+K_M}$ and $f_{ana}(ATP) = \frac{ATP}{ATP+K_M}$.

We used the same parameters as the concentration-independent versions of the model, with the additional parameter being the half-saturation concentration $K_M$ for different metabolites and processes. For simplicity, and with some support from measurements, we assumed that all the $K_M$ values approximately corresponded to 100 $\mu$M, i.e., $6 \times 10^{-7}$ mol/gC in $\chi$ units (Fig. S5). For initial conditions, we assumed arbitrary starting concentrations of 1 mM for both ATP and NADH, since the initial concentrations did not affect the final steady state concentrations. For each set of parameters, we simulated the system of equations to steady state. We discarded parameter combinations where either ATP or ECH did not reach a steady state, as well as those where the steady state concentrations fell outside the To find the optimal growth rate $\lambda_{max}$, we systematically varied the allocation variables $\phi_\alpha$ along an equispaced mesh along the simplex $\sum_\alpha \phi_\alpha = 1$ to ensure that the allocation constraint was satisfied. We then numerically selected the allocation variables corresponding to the largest numerically observed growth rate as $\lambda_{max}$.

Simulating the model with different values of $Z_{C,red}$, we noticed qualitatively and quantitatively identical trends in both models (Fig. S8): $\lambda_{max}$ increased near $Z_{C,red} = Z_{C,B} = 0$. Similar to the zero order model with fixed concentrations, in this model, too, we observed a similar dependence of $\lambda_{max}$ on $Z_{C,red}$ when we changed the reductive ATP yield $S_4$. In particular, reducing $S_4$ resulted in ATP concentrations not reaching a steady state for the same allocation strategy ($\phi_\alpha$), indicating that the metabolic constraints were infeasible for the same strategy. Lowering $\phi_{ana}$ however, allowed an ATP steady state, albeit resulting in a lower growth rate $\lambda_{max}$.

#### 6.2 Optimization

Since our simulations had to sample a high-dimensional simplex of allocation variables and were computationally expensive, we could not sample the space of solutions finely enough. Due to this, our numerically computed maximum growth rates showed sudden jumps as we changed $Z_{C,red}$. Given that this involves optimization of a non-convex equation system, we took measures to ensure the robustness of our results by performing optimization from a large number of randomly sampled initial conditions. Optimization was done over the allocation strategy $\phi_\alpha$ and the ATP and ECH concentrations to maximize the growth rate, while satisfying all three constraints (ATP and ECH balance as well as the allocation constraint). To stay in the regime of biologically plausible ATP and ECH concentrations, we placed bounds on the final ATP and ECH concentrations to be within a reasonable range ($6 \times 10^{-6}$ to $6 \times 10^{-5}$ mol/gC (1–10 mM) for ATP, and $6 \times 10^{-7}$ to $1.2 \times 10^{-6}$ mol/gC (100–200 $\mu$M) for ECH (NADH) in $\chi$ units).

Using numerical optimization, we sampled different $S_4$ and $S_3$ values for both zero order fluxes ($J_\alpha$ independent of ATP and ECH concentrations, i.e., $f(\vec{\chi}) = 1$) as well as Michaelis-Menten fluxes ($J_\alpha$ dependent on ATP and ECH with $f(\vec{\chi})$ set according to Michaelis-Menten kinetics). The results of these optimizations — $\lambda_{max}$ as a function of $Z_{C,red}$ — are shown in Figs. S8 and S9. In these figures, we first show the optimal growth rate in conditions where no additional ATP homeostasis is allowed ($\phi_h = 0$; Fig. S8), and later when we allow ATP homeostasis to be optimized for maximum $\lambda$ ($\phi_h \geq 0$; Fig. S9). In all conditions, we observe that decreasing the reductive ($S_4$) and oxidative ($S_3$) ATP yields can only broaden, not reduce the viable range of $Z_{C,red}$ where $\lambda > 0$. In these conditions, lowering either ATP yield typically reduces growth rate, as observed in other variants of the model.

| carbon source | $O_2$ | $NO_3^-$ | DMSO | TMAO | fermentation |
|---|---|---|---|---|---|
| alcohol-ethanol | grows | grows | grows | grows | no growth |
| amino acid-alanine | grows | grows | grows | grows | no growth |
| amino acid-arginine | grows | grows | grows | grows | no growth |
| amino acid-asparagine | grows | grows | grows | grows | grows |
| amino acid-aspartate | grows | grows | grows | grows | grows |
| amino acid-cysteine | grows | grows | grows | grows | no growth |
| amino acid-glutamate | grows | grows | grows | grows | grows |
| amino acid-glutamine | grows | grows | grows | grows | grows |
| amino acid-glycine | grows | grows | grows | grows | no growth |
| amino acid-serine | grows | grows | grows | grows | grows |
| amino acid-threonine | grows | grows | grows | grows | no growth |
| amino acid-tryptophan | grows | grows | grows | grows | grows |
| aromatic-phenylpropanoate | grows | no growth | no growth | no growth | no growth |
| aromatic-phenylacealdehyde | grows | no growth | no growth | no growth | no growth |
| fatty acid-decanoate | grows | grows | no growth | no growth | no growth |
| fatty acid-dodecanoate | grows | grows | no growth | no growth | no growth |
| fatty acid-hexadecanoate | grows | grows | no growth | no growth | no growth |
| fatty acid-hexadecenoate | grows | grows | no growth | no growth | no growth |
| fatty acid-hexanoate | grows | grows | no growth | no growth | no growth |
| fatty acid-octadecanoate | grows | grows | no growth | no growth | no growth |
| fatty acid-octadecenoate | grows | grows | no growth | no growth | no growth |
| fatty acid-octanoate | grows | grows | no growth | no growth | no growth |
| fatty acid-tetradecanoate | grows | grows | no growth | no growth | no growth |
| fatty acid-tetradecenoate | grows | grows | no growth | no growth | no growth |
| nucleobase-cytosine | grows | grows | grows | grows | grows |
| nucleobase-guanine | grows | no growth | no growth | no growth | no growth |
| nucleobase-adenine | grows | grows | grows | grows | grows |
| nucleoside-adenosine | grows | grows | grows | grows | grows |
| nucleoside-cytidine | grows | grows | grows | grows | grows |
| nucleoside-guanosine | grows | no growth | no growth | no growth | no growth |
| nucleoside-inosine | grows | no growth | no growth | no growth | no growth |
| nucleoside-uridine | grows | grows | grows | grows | grows |
| nucleoside-xanthosine | grows | no growth | no growth | no growth | no growth |
| organic acid-fumarate | grows | grows | grows | grows | grows |
| organic acid-acetate | grows | grows | no growth | no growth | no growth |
| organic acid-lactate | grows | grows | grows | grows | no growth |
| organic acid-pyruvate | grows | grows | grows | grows | grows |
| organic acid-succinate | grows | grows | no growth | no growth | no growth |
| sugar-trehalose | grows | grows | grows | grows | grows |
| sugar-fructose | grows | grows | grows | grows | grows |
| sugar-rhamnose | grows | grows | grows | grows | grows |
| sugar-xylose | grows | grows | grows | grows | grows |
| sugar-mannose | grows | grows | grows | grows | grows |
| sugar-arabinose | grows | grows | grows | grows | grows |
| sugar-glucose | grows | grows | grows | grows | grows |
| sugar-fucose | grows | grows | grows | grows | grows |
| sugar-maltose | grows | grows | grows | grows | grows |
| sugar acid-gluconate | grows | grows | grows | grows | grows |
| sugar acid-galacturonate | grows | grows | grows | grows | grows |
| sugar acid-glucarate | grows | grows | grows | grows | grows |
| sugar acid-glucuronate | grows | grows | grows | grows | grows |
| sugar alcohol-glycerol | grows | grows | grows | grows | grows |
| sugar alcohol-sorbitol | grows | grows | grows | grows | grows |

terminal $e^-$ acceptor

**Figure S3. Flux Balance Analysis of *E. coli*'s two dimensional metabolic capabilities**. Each row denotes a carbon source, each column a terminal electron acceptor. Black denotes no growth in the iML1515 model (*1*), while light brown indicates that the modeled $\lambda_{max} > 0.01$ hr$^{-1}$. TMAO denotes trimethylamine N-oxide and DMSO dimethyl sulfoxide. In all cases where growth was feasible, maintenance energy consumption did not exceed it's minimum allowed value of 6.86 mmol ATP/gDW/hr.

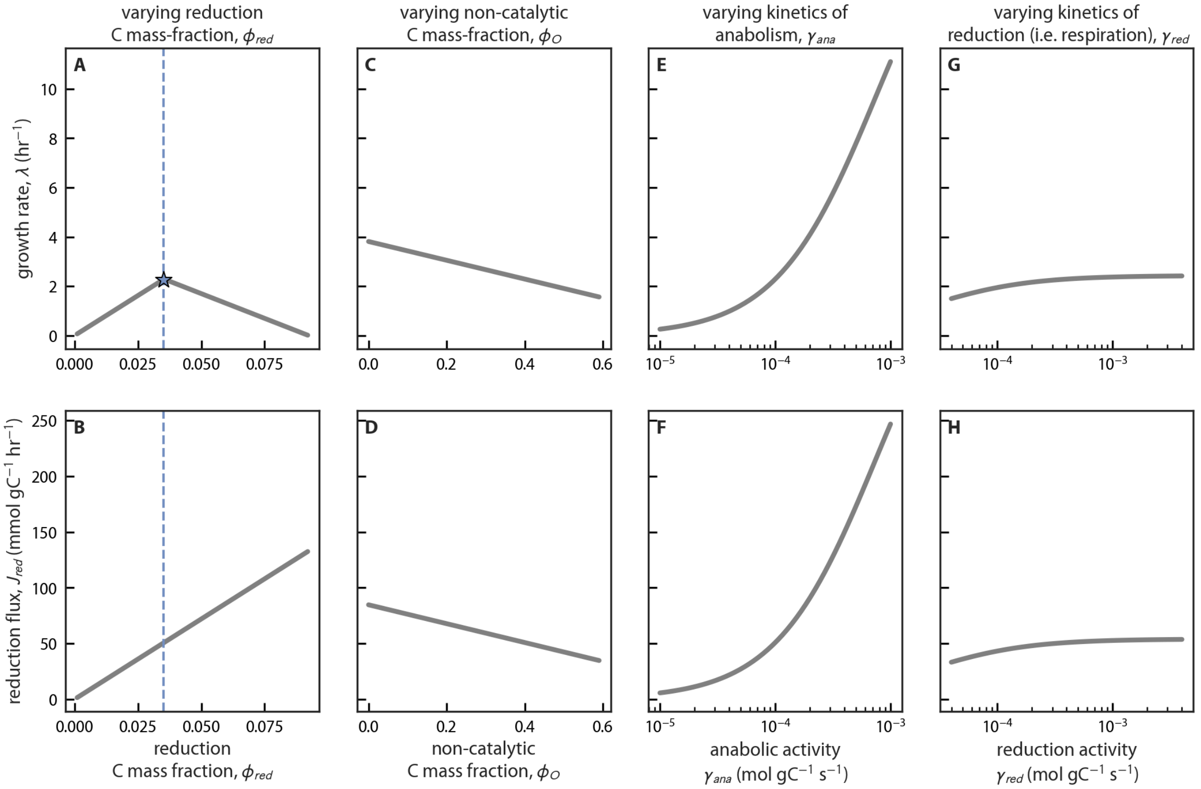

**Figure S4. A redox + resource allocation model of respiration displays a growth-rate maximizing respiratory rate $J^*_{red}$.** Columns of panels vary a model parameter and display the effect on $\lambda_{max}$ and mass-specific respiratory flux $J_{red}$ in a zero-order model optimized using linear methods. Panels (A-B) show that there is a $\lambda$-maximizing value of $J_{red} = \gamma_{red}\phi_{red}f_{red}(\mathbf{c})$ denoted with the star in (A). Panels (C-D) show the effect of increasing the non-catalytic C mass fraction $\phi_O$, (E-F) show the effect of increasing the catalytic activity of anabolic enzyme, $\gamma_{ana}$, and (G-H) the effect of increasing $\gamma_{red}$.

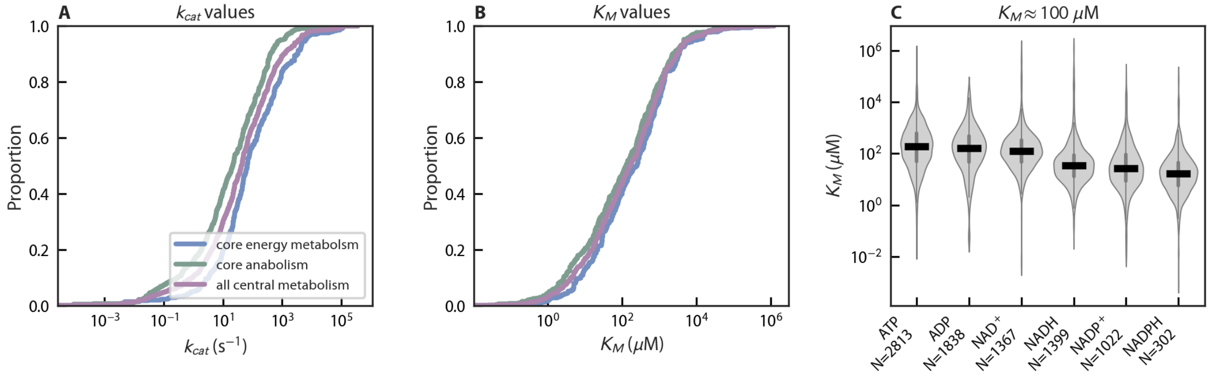

**Figure S5. Typical values of enzyme kinetic constants $k_{cat}$ and $K_M$** (A) CDFs of empirical $k_{cat}$ values for different core metabolic modules. $k_{cat}$ gives the maximum per-active site catalytic rate in mol product/mol enzyme/s. (B) The same, but for $K_M$ in $\mu$M units. (C) Typical $K_M$ values for energy and redox cofactors are $\approx 100$ $\mu$M. Data for all panels from (*29*).

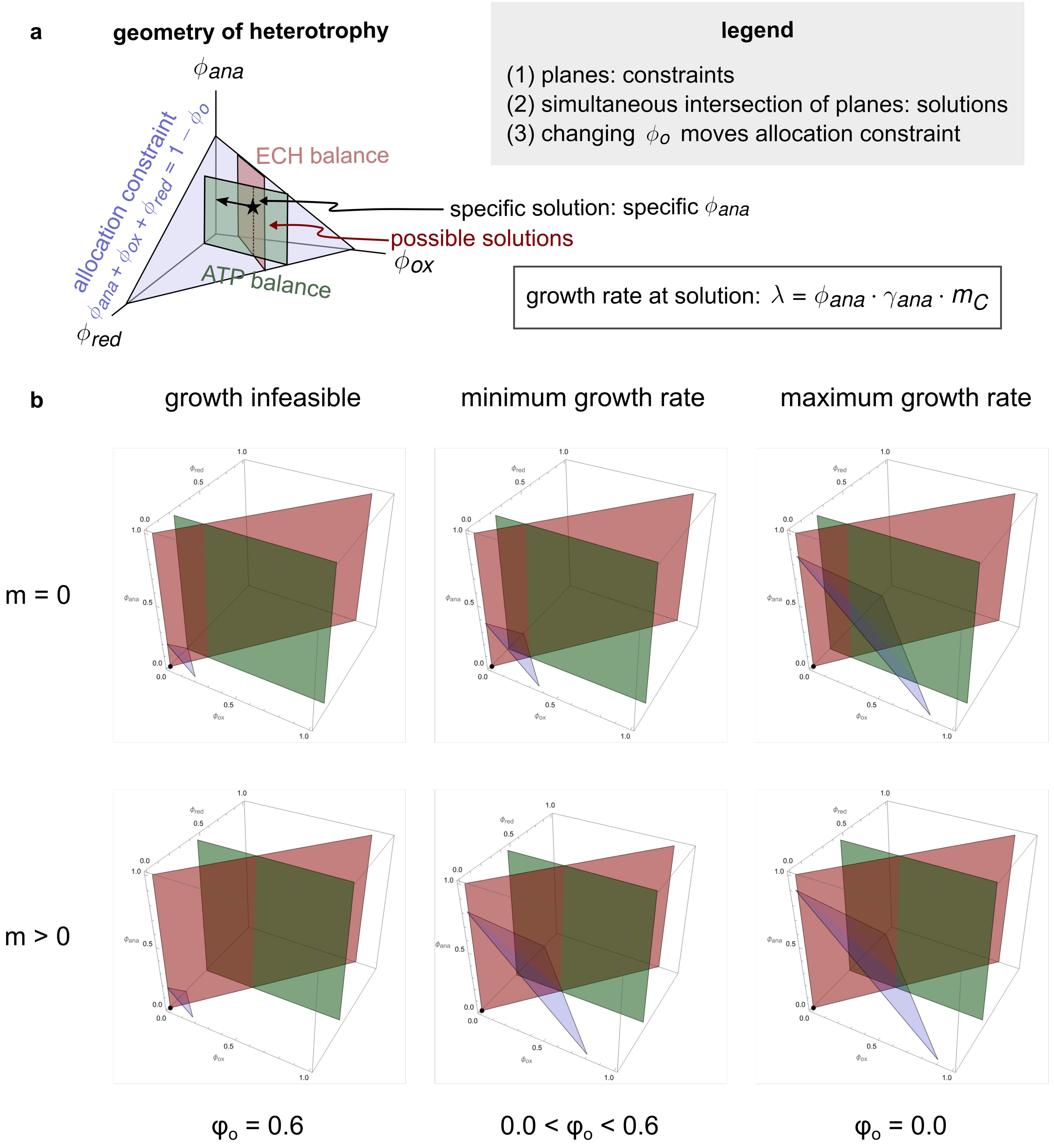

**Figure S6. Geometric visualization of constraints on a zero order model of heterotrophic growth.** (a) Schematic of a 3D space of the allocation variables $\phi_{ana}$, $\phi_{red}$ and $\phi_{ox}$ and the constraints of ECH balance (red), ATP balance (green) and the biomass allocation constraint (blue) as planes in the positive octant. Increasing $\phi_o$ moves the blue allocation plane closer to the origin. If $\phi_o$ is allowed to vary a range of allocation planes and growth rates $\lambda$ are therefore viable. The joint intersection of all three spaces (two mass balance planes and one allocation space) corresponds to possible solutions. The growth rate depends on the $\phi_{ana}$ for the solution (black star) and is given by the formula $\lambda = \phi_{ana} \cdot \gamma_{ana} \cdot m_C$ in our concentration-independent zero order model. (b) Specific model incarnations with no maintenance ($m = 0$, top row) and with a finite maintenance cost ($m = 0.001$, bottom row). In each case, we show three outcomes: one where growth is infeasible (left column, no common intersection); one where growth is just feasible and the growth rate is minimum (middle column; common intersection is a point) and one where the growth rate is maximized (right column, where $\phi_O$ achieves its minimum allowed value). A similar visualization is also valid for higher-order, concentration-dependent, models albeit where planes move with ATP and ECH concentrations in more complex ways.

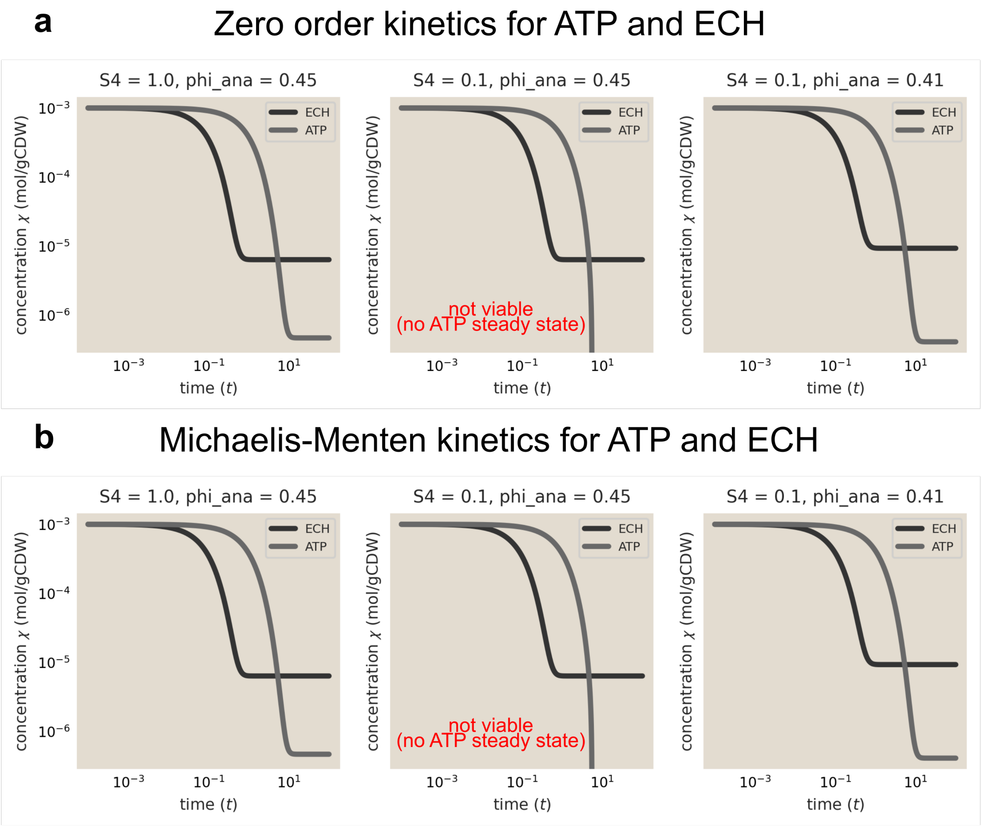

**Figure S7. Simulations highlight the effect of changing ATP yields on growth.** Decreasing the reductive ATP yield $S_4$ affects the viability of growth for the same allocation strategy. Shown here are ATP and ECH dynamics simulated numerically using two model variants (see details in section 6.1). In these simulations, ATP and ECH concentrations dynamically reach steady state, unlike in other versions of the model where it is fixed. (a) shows simulations of the zero order model, where the fluxes $J_\alpha$ are independent of ATP and NADH concentrations; (b) shows simulations where the fluxes depend on the concentration through Michaelis-Menten kinetics. Shown are cases with no additional ATP homeostasis (i.e., $\phi_h = 0$). In both cases, from left to right, we show the following: (left) at $S_4 = 1$, we show a simulation with the optimal allocation strategy with $\phi_{ana} = 0.45$; in this case both ATP and ECH reach a steady state; (middle) at a lower $S_4 = 0.1$, for the same strategy, ATP no longer reaches a steady state and instead dips to arbitrarily small concentrations over time; (right) for the same $S_4 = 0.1$, lowering $\phi_{ana}$ to 0.41 and appropriately adjusting the other allocation variables now allows both ATP and ECH to reach a steady state. This lowering of $\phi_{ana}$ results in a lower optimal growth rate at a lower $S_4$. Time shown on the x-axis is in arbitrary units.

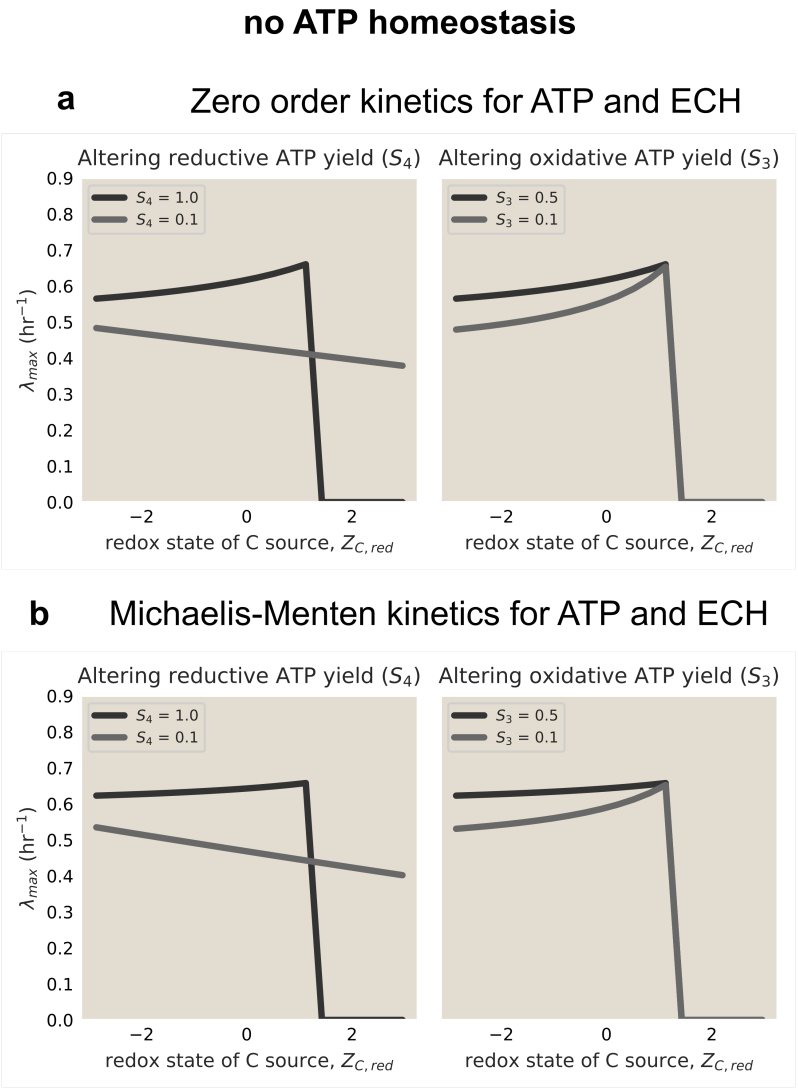

**Figure S8. Optimization without ATP homeostasis shows the effect of changing ATP yields on growth rate $\lambda$.** Decreasing the reductive (left column, $S_4$) ATP yield can broaden the range of viable carbon sources, similar to Figs. 2 and S11. However, decreasing the oxidative ATP yield (right column, $S_3$) only has a quantitative effect of changing growth rates without affecting the breadth of viable carbon sources $Z_{C,red}$. Shown here are results from model variants (detailed in section 6.2) that we numerically optimized. In contrast with Figs. 2 and S11, here ATP and ECH concentrations are not fixed, but optimized maximize $\lambda$ while satisfying all model constraints. In this figure, optimizations disallow allocation towards ATP homeostasis ($\phi_h = 0$). Additionally, we place bounds on the ATP and ECH concentrations over a biologically-plausible range — $6 \times 10^{-6}$ to $6 \times 10^{-5}$ mol/gC (1–10 mM) for ATP, and $6 \times 10^{-7}$ to $1.2 \times 10^{-6}$ mol/gC (100–200 $\mu$M) for ECH. ADP and EC concentrations were assumed to always be a fixed factor $r_a = 0.1$ and $r_e = 10$ times the ATP and ECH concentrations, respectively. Panel (a) shows optimization of the zero order model where fluxes $J_\alpha$ are independent of ATP and NADH concentrations; (b) shows optimization where the fluxes depend on the concentration via Michaelis-Menten kinetics.

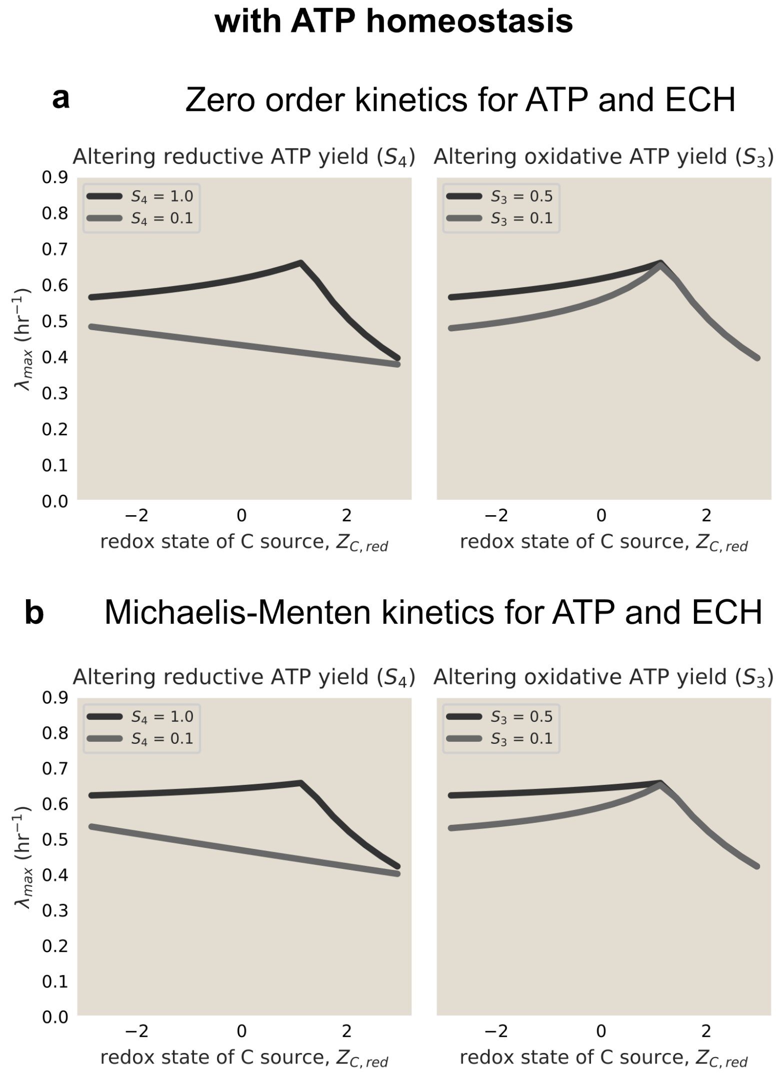

**Figure S9. Flux-balancing ATP homeostasis enables growth across a wide range of C sources also in a nonlinear model.** In contrast with Fig. S8, where flux-balancing ATP homeostasis was disallowed, here we optimize models with non-zero $\phi_H$ and $J_H$. As in the linearized model described in the main text, the addition of flux-balancing homeostasis permits growth on a wide range of carbon sources characterized by $Z_{C,red}$. Decreasing the reductive (left column) and oxidative (right column) ATP yield ($S_4$ and $S_3$ respectively) does not shrink the range of viable carbon sources, but can reduce growth rates, as qualitatively seen in the main text. Shown here are results from model variants (detailed in section 6.2) we numerically optimized, where the ATP and ECH are optimized to achieve maximize $\lambda$ while satisfying the constraints of ATP and ECH balance and biomass allocation. Simulations include the possibility of flux-balancing homeostasis via ATP hydrolysis. Additionally, we placed bounds on the ATP and ECH concentrations over a biologically plausible range ($6 \times 10^{-6}$ to $6 \times 10^{-5}$ mol/gC (1–10 mM) for ATP, and $6 \times 10^{-7}$ to $1.2 \times 10^{-6}$ mol/gC (100–200 $\mu$M) for ECH (NADH) in $\chi$ units). Panel (a) shows optimization of the zero order model, where the fluxes $J_\alpha$ are independent of ATP and ECH concentrations; (b) shows optimization where the fluxes depend on the concentration through Michaelis-Menten kinetics.

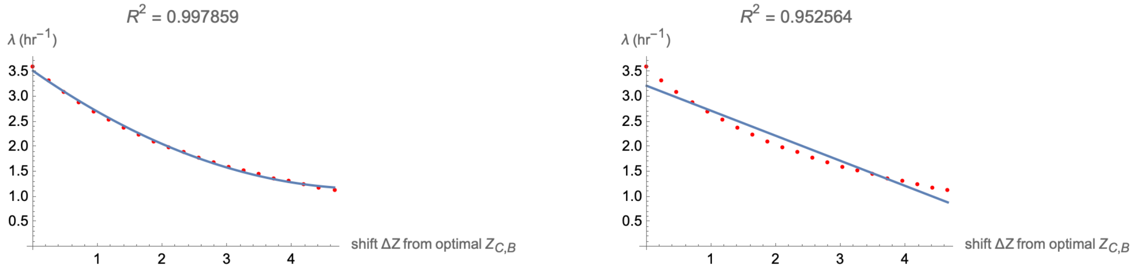

**Figure S10. Growth rate penalty for biomass $Z_{C,B}$ deviating from the optimum is approximately quadratic.** Plot showing the growth rate $\lambda$ as a function of the deviation from optimal $Z_{C,B}$ for the same model of respiratory heterotophy as Fig. 4A. Red points give growth rates obtained by numerically solving our model along a line perpendicular to the optimal line $Z_{C,B} = K_Z Z_{C,red} + Z_0$. The x-axis indicates the distance from the optimal line $\Delta Z$. Blue lines indicate quadratic (left) and linear (right) fits, indicating a clear quadratic trend.

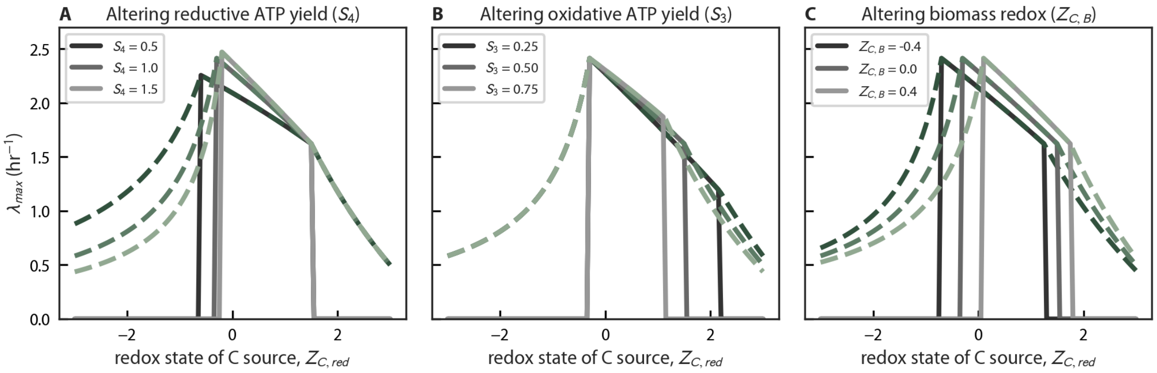

**Figure S11. An intrinsic tension between metabolic flexibility and maximum growth rates.** Plots are for the zero-order model presented in the main text. (A) Increasing the ATP yield of reduction ($S_4$) makes respiration more thermodynamically efficient. This increases the modeled $\lambda_{max}$ at the expense of decreasing the range of reduced C sources that can be consumed without requiring ATP homeostasis (dashed green lines). (B) Likewise, increasing the ATP yield of catabolic C source oxidation ($S_3$) reduces the range of usable C sources. (D) Shifting the redox state of biomass ($Z_{C,B}$), e.g. by making more reduced compounds, affects the ECH stoichiometry of anabolism ($S_6$). Making more reduced biomass shifts the curve in the reduced direction (left) while oxidized biomass shifts it to the right. That is, $Z_{C,B} \approx Z_{C,red}$ is advantageous, permitting near-maximal growth rates without requiring flux-balancing homeostasis mechanisms that inevitably use resources (C atoms, energy, catalytic activity) and thereby reduce the growth rate $\lambda$.

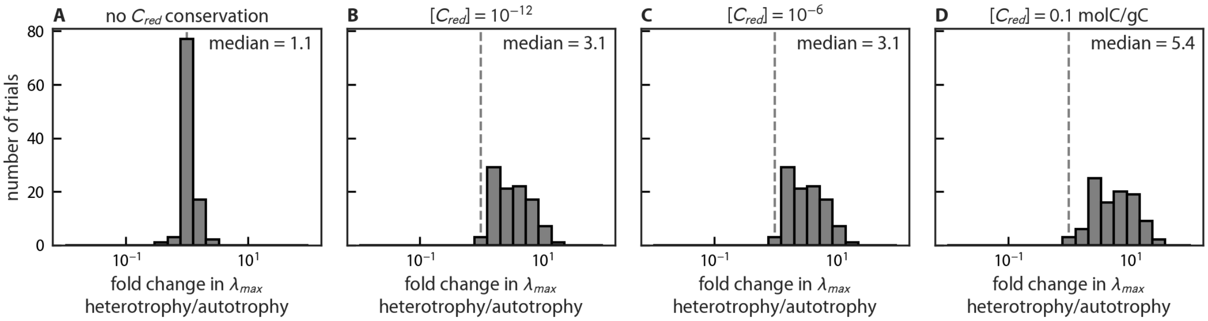

**Figure S12. Enforcing conservation of internally-produced organic carbon ($C_{red}$) limits maximal autotrophic growth rates.** Each panel compares equivalent models of respiration and photosynthesis with identical sampled kinetic parameters ($\gamma_\alpha$) in a zero-order model as in the main text. In (A) the photosynthetic model was not forced to balance production and consumption of $C_{red}$. In this case, photosynthetic and respiratory growth rates are roughly equal. In panels (B-D) conservation was required at different fixed concentrations of $C_{red}$ given in the title and respiratory (heterotrophic) growth rates always exceed autotrophic (photosynthetic) ones. This effect depends on the $C_{red}$ concentration (units of mol C per gram biomass C) when it is very high, as in panel (D). Note that using the same kinetic parameters in both models means that we neglect kinetic difficulties often associated with specific processes, for example $CO_2$ fixation through rubisco in the Calvin-Benson cycle (*32*, *33*). So our analysis here indicates that the fastest-growing autotrophs would grow more slowly than similarly distinguished heterotrophs even if rubisco was a "typical" central metabolic enzyme (*29*).

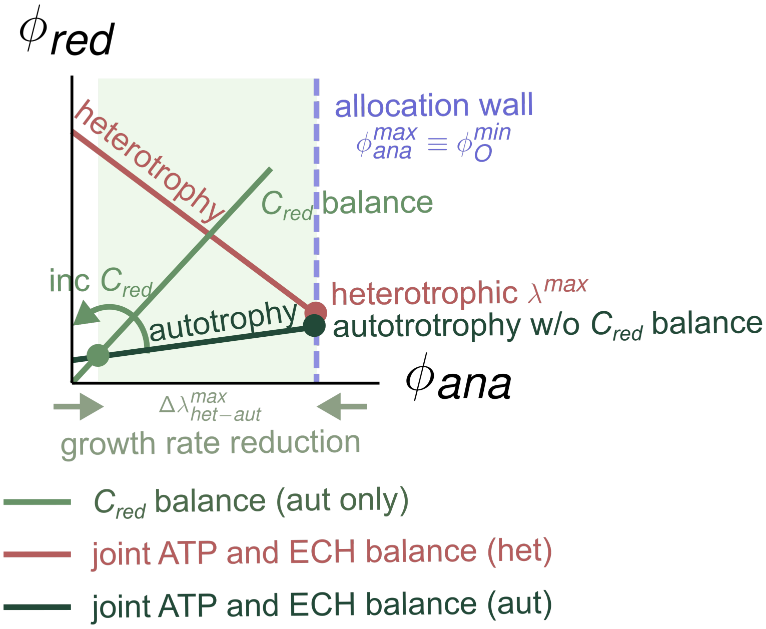

**Figure S13. Geometric explanation as to why balancing intracellularly-produced organic C during autotrophic growth reduces $\lambda$.** Schematic of the 2D space of allocation variables $\phi_{red}$ and $\phi_{ana}$, along with relevant constraints for comparable heterotrophic (red) and autotrophic (green) metabolisms. The solid lines show the constraints on each metabolism: joint ECH and ATP balance in heterotrophy (red), ECH and ATP balance in autotrophy (dark green), $C_{red}$ balance (green) and the biomass allocation constraint (blue). Despite almost all parameter values being the same, ECH and ATP balance constraints in autotrophy and heterotrophy differ in slope due to a sign flip in the ATP stoichiometry of reduction $S_4$. $C_{red}$ balance is an additional constraint that applies only to autotrophy (see Fig. 3 in the main text). This additional constraint involves $\phi_{ana}$ and $\phi_{red}$ and represents a line at a fixed angle depending on the steady-state concentration of reduced carbon $C_{red}$. Without this constraint, the maximum growth rates $\lambda^{\max}$ in both autotrophy and heterotrophy are comparable (filled red and dark green circles) because they are essentially determined by the kinetics of anabolism ($\gamma_{ana}$) which we assumed to be identical. With this constraint, the only viable autrophic growth rate is the one where the $C_{red}$ balance line cuts the ECH and ATP balance line (filled light green circle), which corresponds to a much lower $\phi_{ana}$ and thus a lower autotrophic growth rate. At any non-negligible steady-state $C_{red}$ concentration, the additional $C_{red}$ balance constraint reduces $\phi_{ana}$, yielding a $\approx 3$-fold lower autotrophic growth rate when compared to a model of heterotrophy with identical $\gamma_\alpha$ values, as shown in Fig. 3 and S12.

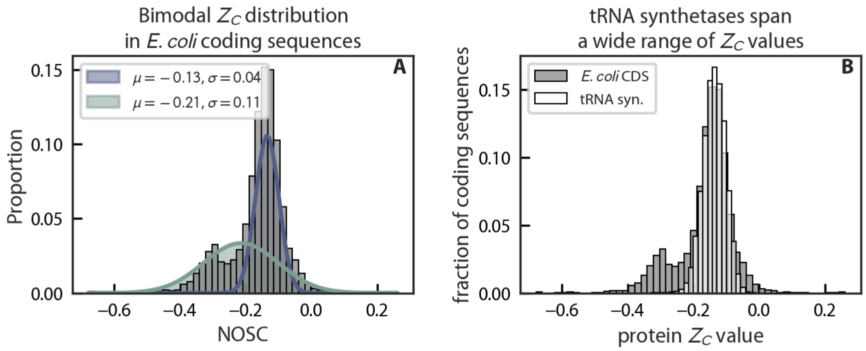

**Figure S14. Aminoacyl tRNA synthetases span a wide range of $Z_C$ values.** (A) *E. coli* protein coding sequence $Z_C$ values are bimodally distributed with two approximately Gaussian components. The blue fitted Gaussian component represents globular proteins while the green represents membrane proteins (*13*). Membrane proteins are more reduced (fit mean $\mu = -0.21$ e$^-$/C and standard deviation $\sigma = 0.11$) than cytosolic proteins ($\mu = -0.13$ e$^-$/C and $\sigma = 0.04$). (B) The grey distribution in the background reproduces the distribution of *E. coli* $Z_C$ values from (A). The distribution of $Z_C$ values of aminoacyl tRNA synthetases across $\approx 60,000$ GTDB representative genomes is given on top in white. The mean and variance of tRNA synthetase $Z_C$ values ($\mu = -0.14$ e$^-$/C and $\sigma = 0.03$) is comparable to soluble proteins in *E. coli*.

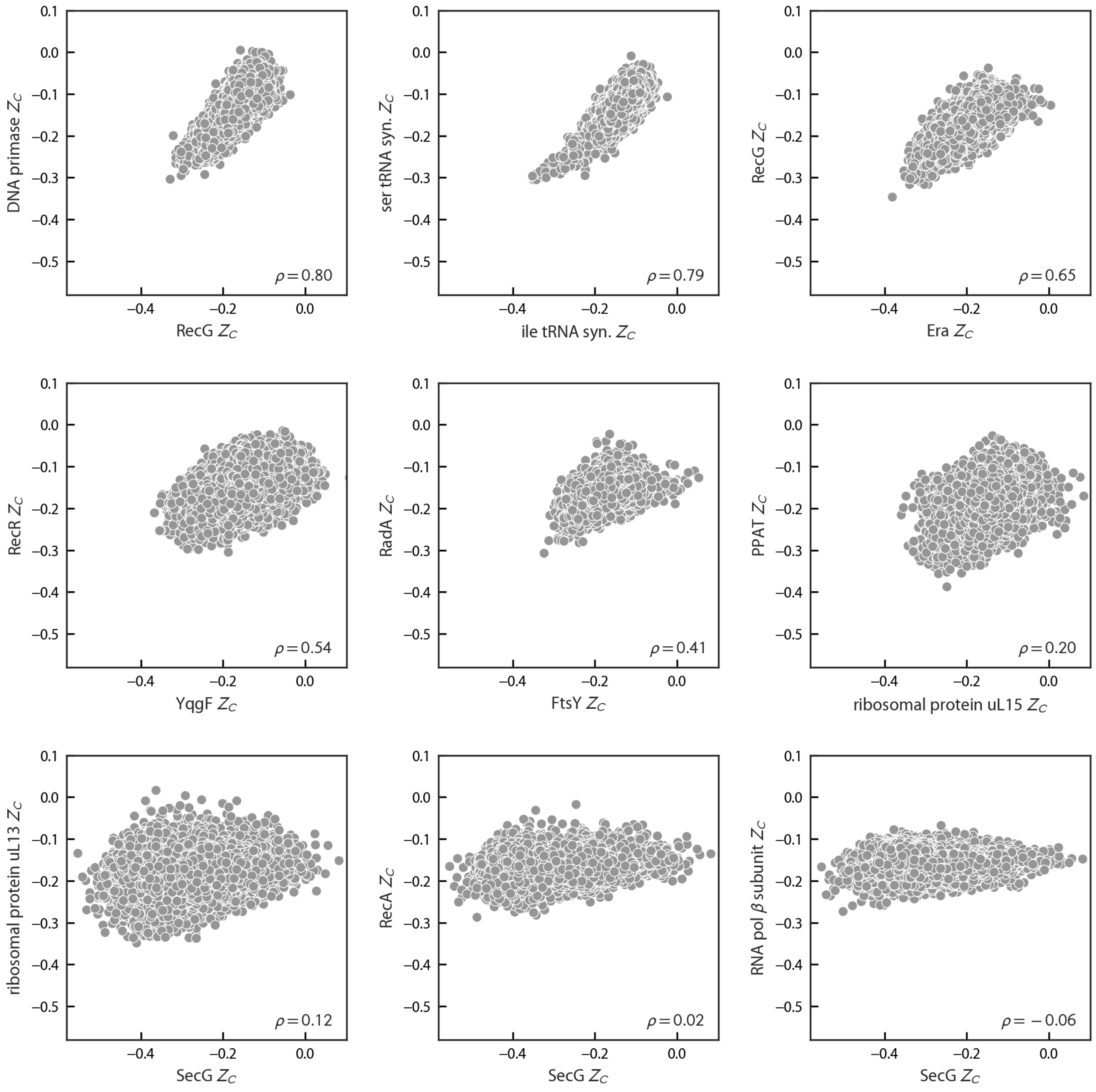

**Figure S15. A range of Pearson correlation coefficients between vectors of $Z_C$ values calculated for bac120 genes in GTDB representative genomes.** This gallery spans the full range of raw correlation coefficients, $\rho$, more than 99% of which were within the interval [0.077, 0.77]. As indicated by Fig. 4, nearly all raw correlations were positive.

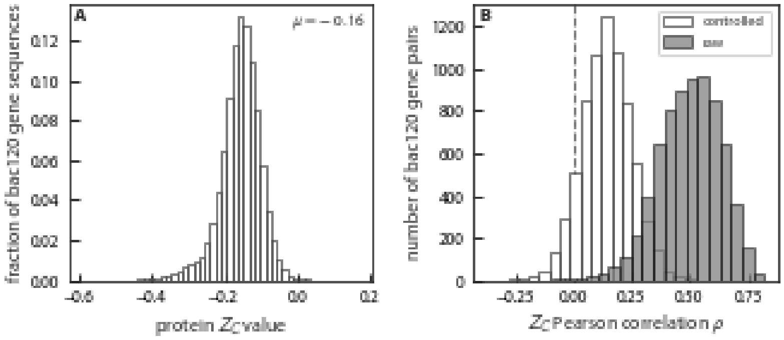

**Figure S16. Controlling for the mean $Z_C$ of protein coding sequences.** Panel (A) gives a histogram of $Z_C$ values for all *bac120* sequences analyzed. This distribution has a mean of $\mu = -0.158$ and a standard deviation of $\sigma = 0.055$. Panel (B) gives raw Pearson correlations between pairs of *bac120* genes (dark grey). These were almost uniformly positive, with an interquartile range IQR = +0.41-0.59. Controlling for the mean $Z_C$ of coding sequences in each genome eliminated much of this correlation (partial correlation IQR = +0.07-0.21, white bars), indicating that most of the effect is genome-wide. The residual positive correlation coincides with our intuition that *bac120* are indeed distinct, given their central and predominantly anabolic roles.

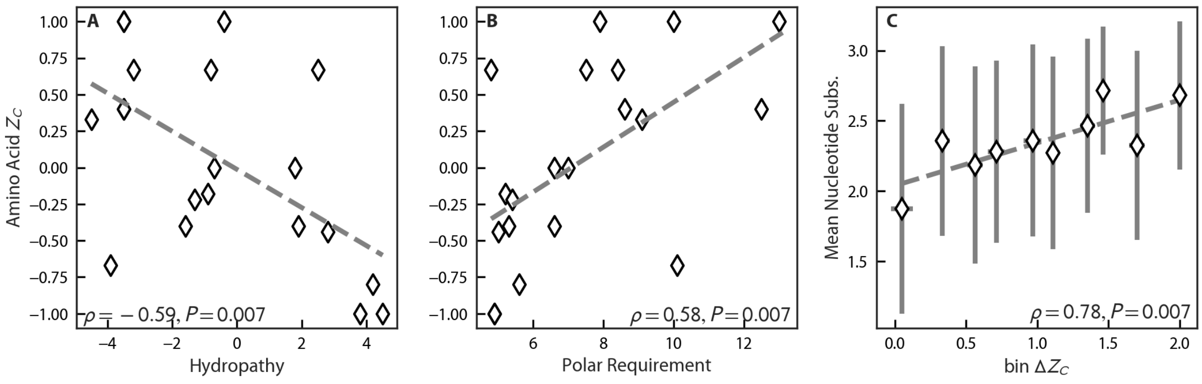

**Figure S17. Altering amino acid $Z_C$ via nucleotide changes typically requires more than one substitution.** The genetic code is conservative for measures of hydrophobicity (*8*, *9*). As such, single nucleotide changes tend to conserve hydrophobicity metrics. Panels (A-B) show that amino acid $Z_C$ is correlated with two different metrics of hydrophobicity — hydropathy and polar requirement (*8*, *9*) – which are rough inverses of each other. For panel (C), pairs of codons were binned by the $Z_C$ difference ($\Delta Z_C$) between their associated amino acids, assuming the standard genetic code. Notice that substitutions that meaningfully alter amino acid $Z_C$ — i.e. lead to a high $\Delta Z_C$ on the right — require a larger number of nucleotide substitutions on average.

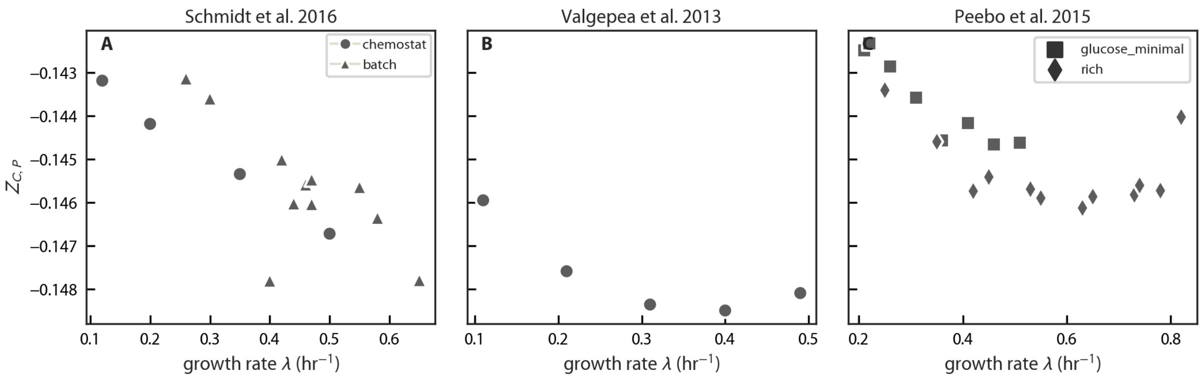

**Figure S18. Various *E. coli* proteomic datasets evidence similar $Z_C$ changes**. Despite differences in extraction, quantification and analysis, three different proteomic datasets show that *E. coli*proteins become more reduced during faster growth rates.

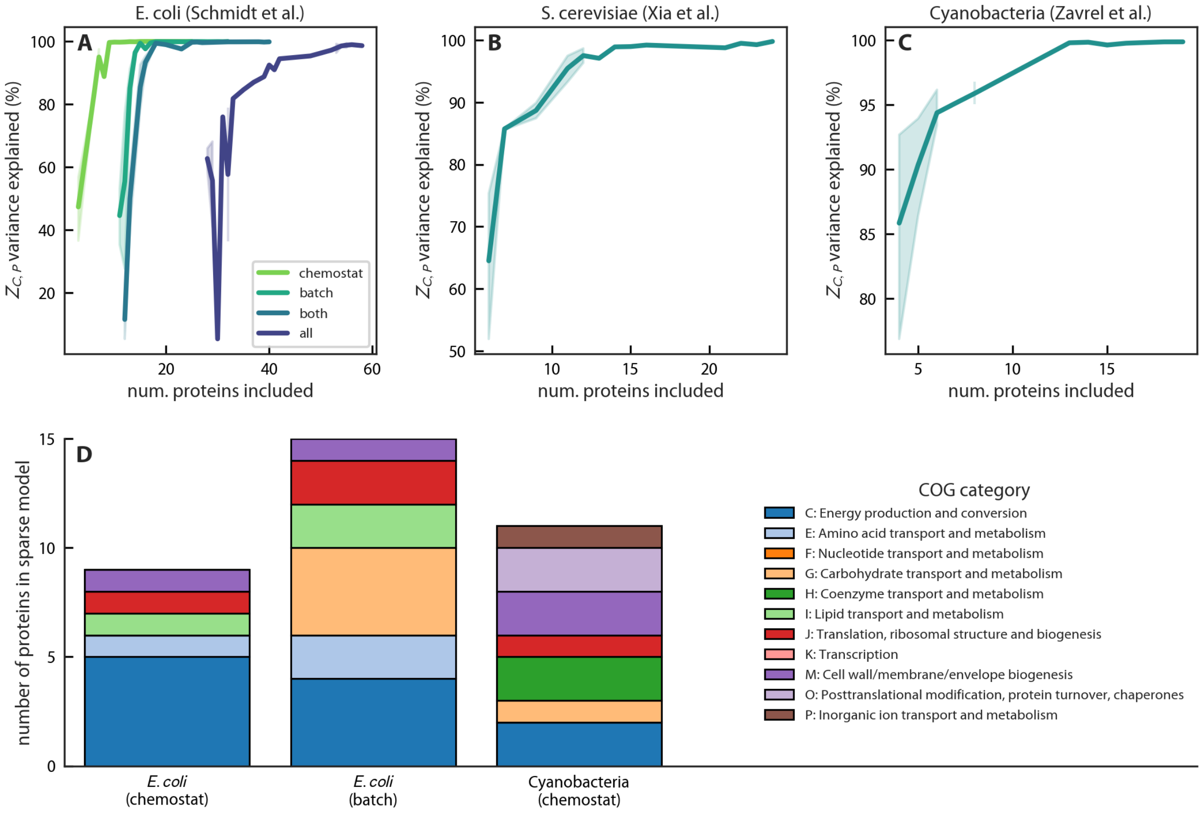

**Figure S19. Trends in proteome $Z_C$ with $\lambda$ are due to many groups of proteins.** Because proteome $Z_{C,P}$ is a carbon-weighted average of individual proteins' $Z_C$ values, a linear model can certainly account for 100% of $Z_{C,P}$ variation with $\lambda$. Yet proteins' expression patterns and $Z_C$ values are correlated, meaning that we need not use the full set of condition-dependent expression levels to reconstruct $Z_{C,P}$ changes. We used sparse regression to ask how many individual proteins' expression changes are required reconstruct $Z_{C,P}(\lambda)$. As shown in panels (A-C), roughly 10 proteins are needed to achieve near-perfect reconstructions approaching 100% variance explained. Panel (D) shows that the proteins selected represent a diversity of protein functional groups. Categories are given for models generated with Lasso regression (Python sklearn package) with the regularization parameter $\alpha$ set to $10^{-8}$, which provided near-perfect reconstruction in all cases. COG functional groups are not available for eukaryotic genomes, so yeast data is omitted from this panel.

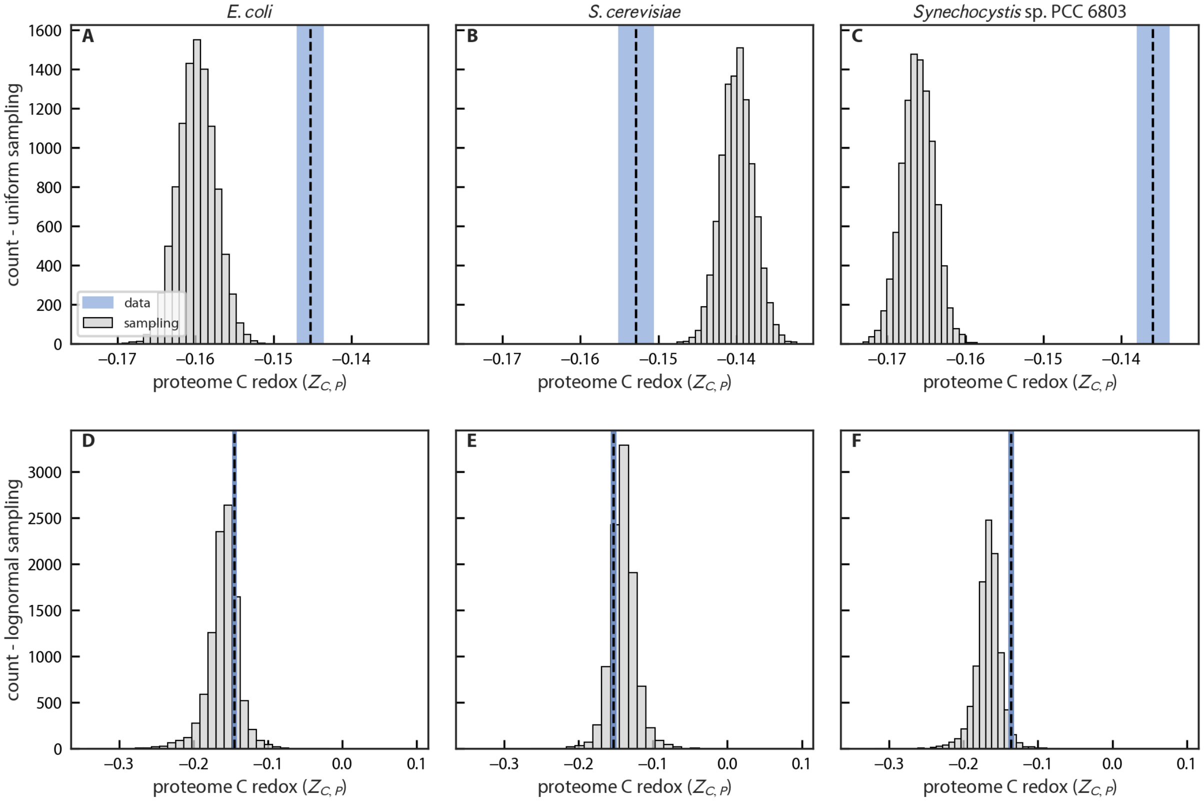

**Figure S20. Protein coding sequences do not accurately estimate the $Z_C$ values of expressed proteins.** In panels (A-C) 1000 individual proteins were repeatedly sampled from the protein coding sequences in each genome to estimate a range of $Z_{C,P}$ values. In all cases, the sampled ranges (gray histograms) did not overlap with the range of measurements (blue range), which account for protein expression. Protein levels are roughly log-normally distributed. Panels (D-F) calculate $Z_{C,P}$ after sampling expression levels from a log-normal distribution spanning roughly 6 orders (i.e. $\sigma \approx 2.3$ estimated from *E. coli* data). This produces much wider ranges of $Z_{C,P}$ values that include measurements.

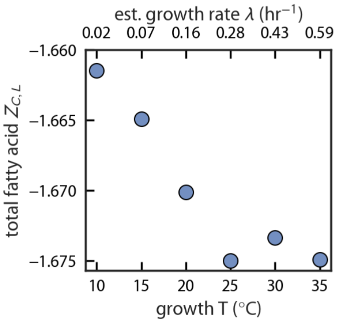

**Figure S21. *E. coli* total lipids $Z_C$ decreases as $\lambda$ increases.** Total lipid composition as a function of growth temperature was reported in (*22*), but the temperature-dependence of the $\lambda$ was not. We estimated $\lambda$ from measurements in similar media from (*23*). This estimate is given on the upper X axis.

#### References

1. J. M. Monk *et al.*, en, *Nat. Biotechnol.* **35**, 904–908 (Oct. 2017).
2. R. U. Ibarra, J. S. Edwards, B. O. Palsson, en, *Nature* **420**, 186–189 (Nov. 2002).
3. Y. Zhou, J. A. Imlay, en, *MBio* **13**, e0296521 (Apr. 2022).
4. H. Bremer, P. P. Dennis, en, *EcoSal Plus* **3**, 1–49 (Sept. 2008).
5. J. M. Dick, M. Yu, J. Tan, A. Lu, en, *Front. Microbiol.* **10**, 120 (Feb. 2019).
6. D. H. Parks *et al.*, en, *Nat. Biotechnol.* (Apr. 2020).
7. M. Y. Galperin *et al.*, en, *Nucleic Acids Res.* **49**, D274–D281 (Jan. 2021).
8. L. Shenhav, D. Zeevi, en, *Science* **370**, 683–687 (Nov. 2020).
9. D. Haig, L. D. Hurst, en, *J. Mol. Evol.* **33**, 412–417 (Nov. 1991).
10. M. Kanehisa, Y. Sato, M. Kawashima, M. Furumichi, M. Tanabe, en, *Nucleic Acids Res.* **44**, D457–62 (Jan. 2016).
11. N. A. O'Leary *et al.*, en, *Nucleic Acids Res.* **44**, D733–45 (Jan. 2016).
12. D. E. LaRowe, P. Van Cappellen, *Geochim. Cosmochim. Acta* **75**, 2030–2042 (Apr. 2011).
13. J. M. Dick, en, *J. R. Soc. Interface* **11**, 20131095 (Nov. 2014).
14. K. Peebo *et al.*, en, *Mol. Biosyst.* **11**, 1184–1193 (Apr. 2015).
15. K. Valgepea, K. Adamberg, A. Seiman, R. Vilu, en, *Mol. Biosyst.* **9**, 2344–2358 (Sept. 2013).
16. A. Schmidt *et al.*, en, *Nat. Biotechnol.* **34**, 104–110 (Jan. 2016).
17. J. Xia *et al.*, en, *Nat. Commun.* **13**, 2819 (May 2022).
18. T. Zavřel *et al.*, en, *Elife* **8** (Feb. 2019).
19. Li GW *et al.*, en, *Cell* **157**, 624–635 (Apr. 2014).
20. Taniguchi Y *et al.*, *Science* **329**, 533–538 (July 2010).
21. J. M. Dick, D. Meng, en, *mSystems*, e0001423 (June 2023).
22. A. G. Marr, J. L. Ingraham, en, *J. Bacteriol.* **84**, 1260–1267 (Dec. 1962).
23. C. O. Gill, D. M. Phillips, *Food Microbiol.* **2**, 285–290 (Oct. 1985).
24. W. Gottstein, B. G. Olivier, F. J. Bruggeman, B. Teusink, en, *J. R. Soc. Interface* **13** (Nov. 2016).
25. M. Basan *et al.*, en, *Nature* **528**, 99–104 (Dec. 2015).
26. A. L. Koch, in *Methods for General and Molecular Microbiology* (ASM Press, Washington, DC, USA, Apr. 2014), pp. 172–199.
27. A. K. Bryan, A. Goranov, A. Amon, S. R. Manalis, *Proc. Natl. Acad. Sci. U. S. A.* **107**, 999–1004 (Jan. 2010).
28. G. Chure, J. Cremer, *Elife* **12**, e84878 (2023).
29. A. Bar-Even *et al.*, en, *Biochemistry* **50**, 4402–4410 (May 2011).
30. R. Milo, R. Phillips, *Cell Biology by the Numbers*, en (Garland Science, Dec. 2015).
31. M. Stitt, D. Schulze, en, *Plant Cell Environ.* **17**, 465–487 (May 1994).
32. N. Prywes, N. R. Phillips, O. T. Tuck, L. E. Valentin-Alvarado, D. F. Savage, en, *Annu. Rev. Biochem.*, arXiv: `2207.10773 (q-bio.BM)` (Apr. 2023).
33. A. I. Flamholz *et al.*, en, *Biochemistry* **58**, 3365–3376 (Aug. 2019).
34. D. Molenaar, R. van Berlo, D. de Ridder, B. Teusink, *Mol. Syst. Biol.* **5**, 323 (Jan. 2009).
35. E. Noor, W. Liebermeister, "Optimal enzyme profiles in unbranched metabolic pathways", en, Sept. 2023.
36. J. E. Goldford, A. B. George, A. I. Flamholz, D. Segrè, en, *Proceedings of the National Academy of Sciences* **119**, e2110787119 (Apr. 2022).
37. T. M. Hoehler *et al.*, *Proceedings of the National Academy of Sciences* **120**, e2303764120 (2023).
38. S. J. Pirt, en, *Proc. R. Soc. Lond. B Biol. Sci.* **163**, 224–231 (Oct. 1965).
39. D. W. Tempest, O. M. Neijssel, en, *Annu. Rev. Microbiol.* **38**, 459–486 (1984).
40. J. a. Imlay, *Nat. Rev. Microbiol.* **11**, 443–454 (2013).
41. C. Kaleta, S. Schäuble, U. Rinas, S. Schuster, *Biotechnol. J.* **8**, 1105–1114 (Sept. 2013).
42. B. D. Bennett *et al.*, en, *Nat. Chem. Biol.* **5**, 593–599 (June 2009).
43. J. O. Park *et al.*, *Nat. Chem. Biol.* (2016).
44. J. D. Orth, I. Thiele, B. Ø. Palsson, en, *Nat. Biotechnol.* **28**, 245–248 (Mar. 2010).
45. A. Goelzer *et al.*, en, *Metab. Eng.* **32**, 232–243 (Nov. 2015).
46. M. Mori, T. Hwa, O. C. Martin, A. De Martino, E. Marinari, en, *PLoS Comput. Biol.* **12**, e1004913 (June 2016).
47. J. J. Hamilton, V. Dwivedi, J. L. Reed, en, *Biophys. J.* **105**, 512–522 (July 2013).
